# NLIMED: Natural Language Interface for Model Entity Discovery in Biosimulation Model Repositories

**DOI:** 10.1101/756304

**Authors:** Yuda Munarko, Dewan M. Sarwar, Anand Rampadarath, Koray Atalag, John H. Gennari, Maxwell L. Neal, David P. Nickerson

**Affiliations:** Auckland Bioengineering Institute, University of Auckland, Auckland, New Zealand; Department of Biomedical Informatics and Medical Education, University of Washington, Seattle, USA; Center for Global Infectious Disease Research, Seattle Children’s Research Institute, Seattle, USA

**Author notes:** Correspondence: Yuda Munarko.

**Keywords:** semantic annotation, ontology class, Physiome Model Repository, Biomodels, NLP, SPARQL, information retrieval

## Abstract

Semantic annotation is a crucial step to assure reusability and reproducibility of biosimulation models in biology and physiology. For this purpose, the COmputational Modeling in BIology NEtwork (COMBINE) community recommends the use of the Resource Description Framework (RDF). This grounding in RDF provides the flexibility to enable searching for entities within models (e.g. variables, equations, or entire models) by utilising the RDF query language SPARQL. However, the rigidity and complexity of the SPARQL syntax and the nature of the tree-like structure of semantic annotations, are challenging for users. Therefore, we propose NLIMED, an interface that converts natural language queries into SPARQL. We use this interface to query and discover model entities from repositories of biosimulation models. NLIMED works with the Physiome Model Repository (PMR) and the BioModels database and potentially other repositories annotated using RDF. Natural language queries are first ‘chunked’ into phrases and annotated against ontology classes and predicates utilising different natural language processing tools. Then, the ontology classes and predicates are composed as SPARQL and finally ranked using our SPARQL Composer and our indexing system. We demonstrate that NLIMED’s approach for chunking and annotating queries is more effective than the NCBO Annotator for identifying relevant ontology classes in natural language queries. Comparison of NLIMED’s behaviour against historical query records in the PMR shows that it can adapt appropriately to queries associated with well-annotated models.

## 1 INTRODUCTION

The Resource Description Framework (RDF) is a standard data model from the semantic web community that is used in semantically annotated biosimulation models such as those formatted in CellML (Cuellar et al., 2003) and Systems Biology Markup Language (SBML) (Hucka et al., 2003) in the Physiome Repository Model (PMR) (Yu et al., 2011) and BioModels Database (Chelliah et al., 2015). These RDF-annotated models can then be discovered by their model entities, such as variables, components, mathematical formula, reactions, compartments, species, and events. Leveraging these semantic annotations when searching the model repositories improves the discoverability of relevant models, thus supporting communities in biology and physiology by guiding modellers to potential starting points for reuse.

The use of RDF in the semantic annotation of biosimulation models has been formalised as the recommended technology by the COmputational Modeling in BIology NEtwork (COMBINE) community (Neal et al., 2019). Furthermore, the community encourages standardised model annotation to ensure interoperability and sharing between modellers using different platforms (Gennari et al., 2021; Welsh et al., 2021). The standard uses composite annotations (Gennari et al., 2011) for more expressive and consistent model annotation. Composite annotations are logical statements linking multiple knowledge resource terms, enabling modellers to precisely define model elements in a structured manner. Methods such as those presented here are able to make use of that structure to go beyond the raw RDF triples with which a model may be annotated.

Because these semantic annotations are encoded with RDF, we can leverage the SPARQL Query Language for RDF, a standard query language to retrieve information from RDF data. SPARQL is a well-defined and high-performance query language, well-suited for triple (subject, predicate, and object) searches over RDF graphs (Pérez et al., 2009). However, writing SPARQL becomes complicated when searching over biological knowledge associated with complex mathematical models. Following the COMBINE standard, model entities often need to be annotated compositely by several ontology classes to accurately represent the underlying biological knowledge inherent to the model. Moreover, the ontology classes can be connected by a series of predicates and objects, creating a tree structure. Hence, knowledge about the ontology classes and tree structure related to the model is critical, requiring expert users to be able to create SPARQL queries that suit their information needs. Thus, the availability of a tool or interface to assist an ordinary user in composing SPARQL is required.

Several studies answering this challenge have been carried out by developing a graphical user interface (GUI) that provides assistance when building SPARQL for a query. Arenas et al. (2016) and Ferré (2014) have created a textual-based GUI using faceted queries where the user can select one or multiple facets and then specify its coverage. For an easier use of SPARQL features, Ferré (2014) complemented their work with natural language verbalism and readability. Another textual-based GUI was SPARQL Query-Builder which generated the WHERE clause based on ontology classes and attributes (Vcelak et al., 2018). Visual-based GUIs such as SparqlBlocks (Ceriani and Bottoni, 2017) and ViziQuer (Čerāns et al., 2019) provided different user experiences by minimising typing activity. SparqlBlocks implemented block-based programming so the user can quickly drag and drop subjects, predicates, objects, and SPARQL features, while ViziQuer presented a Unified Modeling Language (UML) style interface targeting general IT expert and non-IT users. The advantage of a GUI lies in the ability to create a complex SPARQL query, accommodating various features, however, understanding the RDF data structure and SPARQL logic is still required in all of these examples.

Other studies have addressed this challenge by proposing a converter from query to SPARQL. Using Natural Language Processing (NLP), Hamon et al. (2014) developed POMELO (PathOlogies, MEdicaments, aLimentatiOn), a tool that enriches a query with linguistic and semantic features and then abstracts and constructs SPARQL. A similar approach is used by Yahya et al. (2012) and Xu et al. (2014) who have developed tools that identify the subject, predicate, and object associated with a query and eventually generate SPARQL. Marginean (2014) proposed a different approach using manually generated rules and showed that it performs better than POMELO for biomedical data (QALD-4) (Unger et al., 2014); however, this approach is brittle and does not work for newer or more complex queries. Another approach is SimplePARQL (Djebali and Raimbault, 2015) which is a pseudo-SPARQL query language where the query is made in SPARQL form, but the structure, especially the WHERE clause, is in textual form. With SimplePARQL, SPARQL can be more expressive, and multiple SPARQL queries are created based on the available templates. The query to SPARQL converter offers a seamless interface that allows the user to provide keywords and get results directly. These queries to the SPARQL converter are the most similar to Natural Language Interface for Model Entity Discovery (NLIMED); however, they are less suitable for exploring RDF in biosimulation models that use composite annotations. We differentiate our work by applying different approaches for ontology class identification and SPARQL generation. In addition, our work accommodates complex RDF annotations that are not just a set of triples but rather a tree structure with different depths where the leaves are either ontology classes or literals.

In the biosimulation model communities, to the best of our knowledge, the only study of a query to SPARQL conversion is based on semantic queries with a web interface aimed at finding model entities related to epithelial transport (Sarwar et al., 2019b). This interface was utilised by the Epithelial Modelling Platform (EMP), a visual web tool for creating new epithelial transport models based on existing models (Sarwar et al., 2019a). However, for more general discoveries involving several repositories with various biosimulation models, this interface requires a significant modification.

In this manuscript, we introduce NLIMED, a natural language interface for searching semantically annotated model entities from biosimulation model repositories. In order to translate queries into SPARQL, there are two stages. (1) The terms in the query must be identified as ontology classes. This task is similar to named entity recognition, and could be done by off-the-shelf tools like the NCBO annotator (Jonquet et al., 2010). (2) The now-annotated query can be used to generate a SPARQL query. As an alternative to NCBO Annotator, we apply off-the-shelf NLP tools such as Stanford CoreNLP (Manning et al., 2014) and Stanza (Zhang et al., 2020) to ‘chunk’ a query into phrases and then link the phrases to ontology classes using our proposed similarity measure. We first examine entities’ semantic annotation patterns in a repository involving the detected ontology classes to generate SPARQL. For all patterns that have the detected ontology classes, the SPARQL are generated.

We evaluated NLIMED on all CellML models within the PMR that are annotated with RDF. We show that the NLIMED approach for detecting ontology classes in NLQ is more effective than the NCBO Annotator based on precision, recall, and F-measure statistics with margins above 0.13. In addition to the PMR, we have performed preliminary testing of NLIMED with the BioModels database indexing models encoded in the SBML format. This testing supports our belief that NLIMED can be implementable and adaptable for different biosimulation model repositories and modelling formats, and that it is possible to create a generic model entity discovery tool. Also, comparing the behaviour of NLIMED against historical records of actual queries and results in the PMR shows that NLIMED can perform appropriately on well-annotated biosimulation models. Currently, we have implemented NLIMED in the Epithelial Modeling Platform and Model Annotation and Discovery web tools (Sarwar et al., 2019a) to optimise user experiences when searching for model entities. We provide our implementation and experiment setup at https://github.com/napakalas/NLIMED.

## 2 MATERIALS AND METHODS

Our interface consists of two primary modules, the NLQ Annotator and the SPARQL Generator (Figure 1). Both modules are based on data collected from the PMR (Yu et al., 2011), BioModels (Chelliah et al., 2015), and BioPortal (Whetzel et al., 2011). We utilise natural language parsers provided by Stanford CoreNLP (Manning et al., 2014) and Benepar (Kitaev and Klein, 2018) contained in NLTK (Bird et al., 2009), and named entity recognition for biomedical and clinical data by Stanza (Zhang et al., 2020).

**Figure 1.**
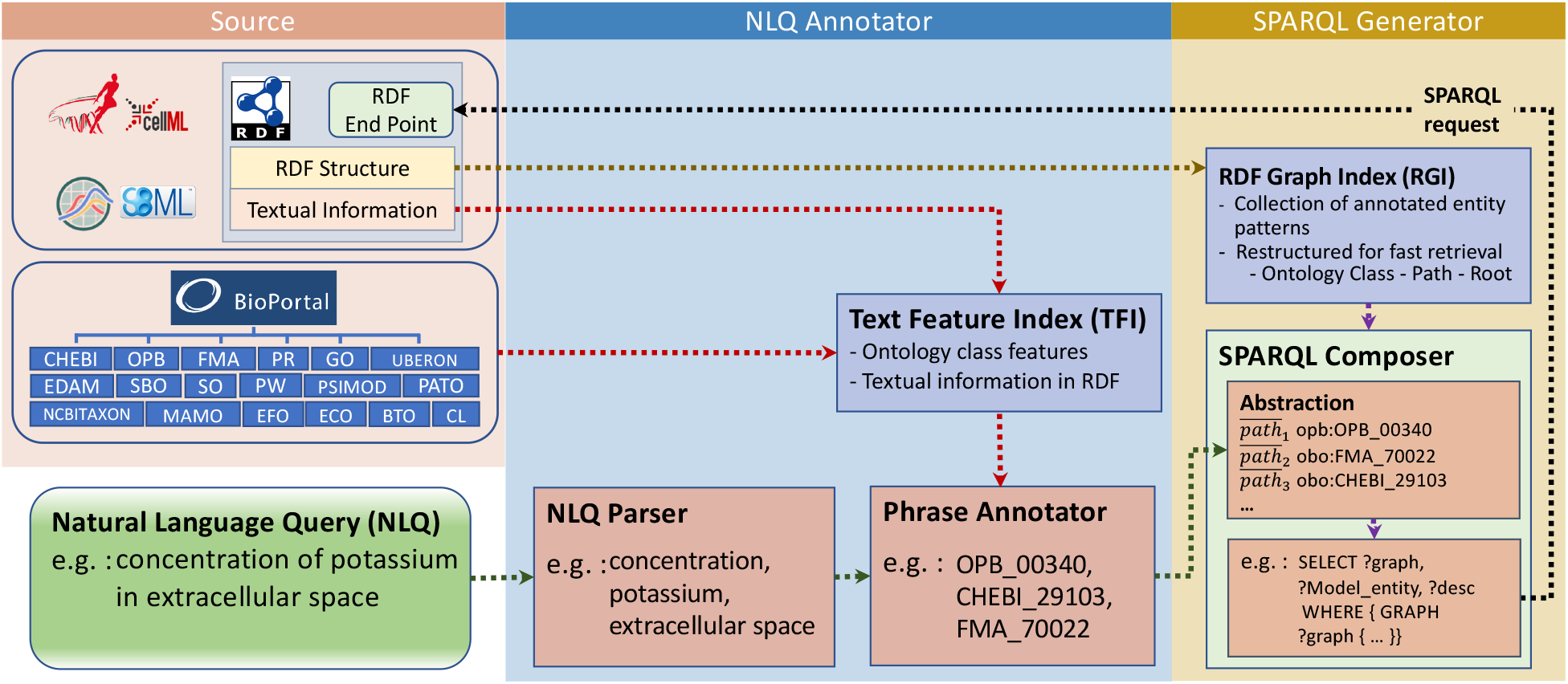
NLIMED workflow. We first create a Text Feature Index (TFI) and an RDF Graph Index (RGI) based on data available in the PMR, BioModels database, and ontology dictionaries. The natural language query is initially annotated into ontology classes in the NLQ Annotator module and then translated into SPARQL in the SPARQL Generator module.

### 2.1 The Physiome Model Repository, BioModels, and BioPortal

The PMR contains more than 800 CellML biosimulation models in which around 25% have been annotated with RDF. Within these annotated models, there are 4,671 model entities annotated described using ontology classes, textual data, and other attributes such as author name, doi, and journal title. Figure 2 is an example of well-annotated entities and shows how this example entity is expressed in RDF, and thus can be viewed as a tree structure, where the leaves are ontology terms. There are 3,472 distinct leaves, 29,755 paths between roots and leaves, and 529 distinct ontology leaves in the models contained in the PMR.

**Figure 2.**
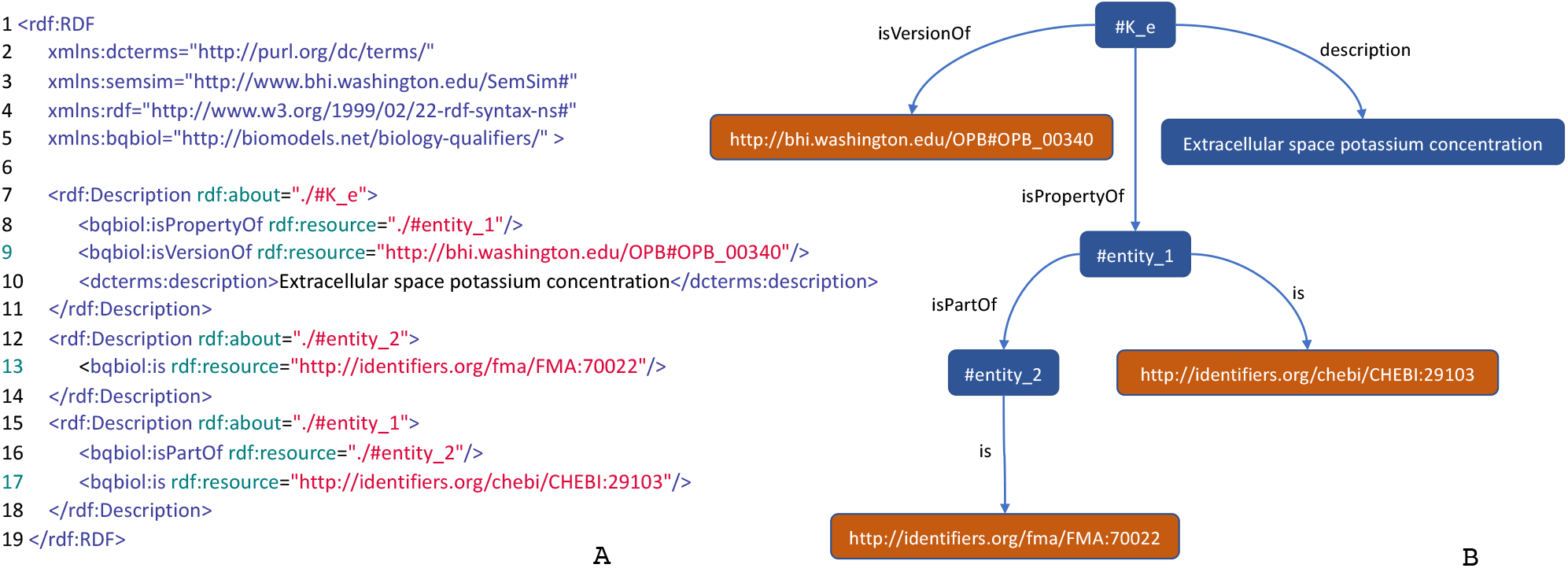
An annotation of a model entity of concentration of potassium in extracellular space. (A) An RDF/XML code describing the model entity. (B) A tree structure representing the RDF code describing the model entity.

The BioModels database has a larger biosimulation model collection consisting of 2,287 SBML files of which 1,039 have been manually annotated. Similar to CellML, model entities are represented as a tree structure but are usually annotated with a simpler format consisting of a single triple linking an entity, a predicate, and an ontology class. Overall, the BioModels database has 13,957 model entities, 46,431 paths connecting roots and leaves, and 2,419 ontology classes.

We utilise BioPortal (Whetzel et al., 2011) to get ontology dictionaries covering ontology classes found in the PMR and most of the ontology classes found in BioModels. In total there are 18 ontology dictionaries collected (Figure 1), of which the most commonly found in the PMR are Gene Ontology (GO), Ontology of Physics for Biology (OPB), Foundational Model of Anatomy (FMA), and Chemical Entities of Biological Interest (ChEBI). Moreover, BioPortal provides the NCBO Annotator tool (Jonquet et al., 2010), which is used as the gold standard to measure NLIMED’s performance. The NCBO Annotator can identify ontology classes in a text based on three features: textual metadata, the number of other ontology classes referring to the ontology class, and the number of classes in the same ontology.

### 2.2 Selecting Features of Ontology Classes to Associate With Query Phrases

Each ontology class collected from BioPortal is described by a set of features such as preferred label, synonym, definition, is a, parents, regex, and comment. These features, then, are used to compute the degree of association of a phrase to an ontology class. However, some features do not have attributes useful to calculate the degree of association, for example, created by, comment, and creation date are aimed to record the class development history, and obsolete, is obsolete, and obsolete since are used to show the class status. Some other features are also limited for specific ontologies, such as regex, part of, example, and Wikipedia, while the last two features tend to consist of a very long text and bias calculations. Therefore, we chose features that can textually represent concepts in the ontology class and are highly related to the user query, i.e. preferred label, synonym, definition, and parent label (Table 1). In addition, many entities in the PMR and BioModel repositories come with textual descriptions that complement the RDF annotations, so we include them as an additional feature.

**Table 1.**
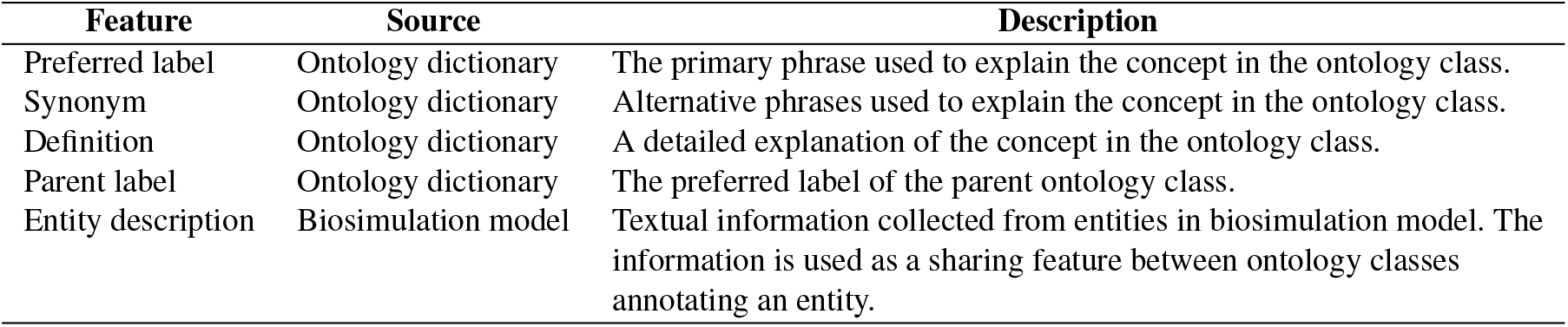
Features representing an ontology class used by the Phrase Annotator to calculate the degree of association of a phrase to the ontology class.

| Feature | Source | Description |
| --- | --- | --- |
| Preferred label | Ontology dictionary | The primary phrase used to explain the concept in the ontology class. |
| Synonym | Ontology dictionary | Alternative phrases used to explain the concept in the ontology class. |
| Definition | Ontology dictionary | A detailed explanation of the concept in the ontology class. |
| Parent label | Ontology dictionary | The preferred label of the parent ontology class. |
| Entity description | Biosimulation model | Textual information collected from entities in biosimulation model. The information is used as a sharing feature between ontology classes annotating an entity. |

### 2.3 Natural Language Query (NLQ) Annotator

The NLQ Annotator is a module for identifying ontology classes with their predicates in an NLQ. It works by chunking the query into candidate phrases using the NLQ Parser, then calculating the degree of association between candidate phrases and ontology classes using the Phrase Annotator (Figure 1). However, calculating the degree of association for all candidate phrases to all ontology classes is inefficient; here, we filter only the likely relevant ontology classes using the Text Feature Index (TFI). The TFI holds a vector space model (Salton et al., 1975) describing ontology classes and adapts an inverted index concept (Harman et al., 1992) to manage features and their relationship to ontology classes. To chunk the query, we investigated the effectiveness of using three NLP tools, CoreNLP (Manning et al., 2014), Benepar (Kitaev and Klein, 2018), and Stanza (Zhang et al., 2020). Stanza is a tool that differs from the other two in that it functions as a Named Entity Recognition (NER) to identify biomedical and clinical entities directly. In contrast, others are parsers that are only used to identify phrases. For simplicity, each approach will be referred to as a parser.

The use of parsers (Figure 3, left side) initially identifies all possible noun phrases, fragments, and sentences as candidate phrases. For example, the NLQ of ‘concentration of potassium in extracellular space’ is chunked to *cp*_1_ : ‘concentration of potassium in extracellular space’, *cp*_2_ : ‘concentration of potassium’, *cp*_3_ : ‘concentration’, *cp*_4_ : ‘potassium’, *cp*_5_ : ‘extracellular space’, and *cp*_6_ : ‘space’ by CoreNLP. The more interesting phrases are those with the highest degree of association to an ontology class while having the most extended terms without overlap while fully covering the NLQ. For the example NLQ, *cp*_3_ : ‘concentration’, *cp*_4_ : ‘potassium’, and *cp*_5_ : ‘extracellular space’ are selected, while the other phrases are considered as irrelevant and removed.

**Figure 3.**
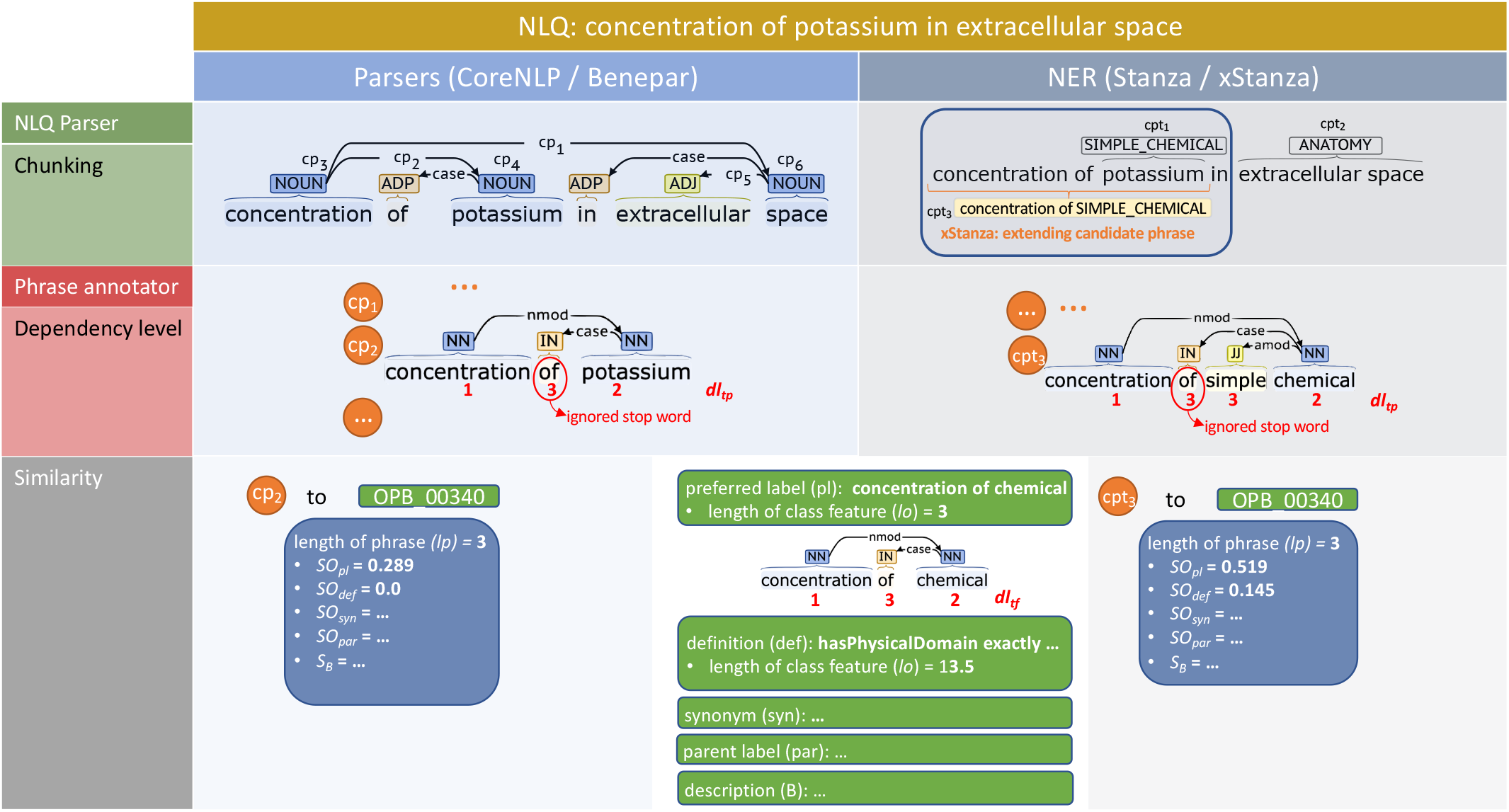
Example NLQ annotation including chunking, dependency level recognition, and similarity calculation for ‘concentration of potassium in extracellular space’. The left side presents the use of parsers (CoreNLP and Benepar) while the right side presents the use of NER (Stanza and xStanza).

The implementation of NER (Figure 3, right side) makes it possible to identify candidate phrases along with entity types. This allows evaluation of the phrase’s context and the identification of additional candidate phrases and related predicates, which are used as extra variables for SPARQL ranking. We utilise three biomedical datasets, AnatEM (Pyysalo and Ananiadou, 2014), BioNLP13CG (Pyysalo et al., 2015), JNLPBA and one clinical dataset (Kim et al., 2003), i2b2 (Uzuner et al., 2011), covering most required entity types including anatomy, chemical, protein, gene, DNA, RNA, cell line, cell type, organ, tissue, amino acid, problem, test, and treatment. Considering the example NLQ; initially we get phrase:entity-type pairs such as *cpt*_1_: ‘potassium - SIMPLE CHEMICAL’ and *cpt*_2_: ‘extracellular space - ANATOMY’. Then the contexts related to each *cpt* are identified, for example the *cpt*_1_ context phrase is ‘concentration of simple chemical’. The context phrase has a higher degree of association with an ontology class rather than a predicate, and so it is selected as an additional phrase *cpt*_3_.

The degree of association between a candidate phrase and an ontology class is the addition of similarity values of the candidate phrase to each feature in the ontology class. Since candidate phrases and features derived from ontology dictionaries are generally short, the similarity is simply the proportion of overlapping terms in candidate phrase *P* and feature *F* normalised by the number of terms in feature *F* (Equation (1)).

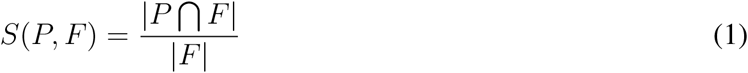

However, longer candidate phrases tend to have high similarity value when the feature text is shorter. For example, *cp*_1_ : ‘concentration of potassium in extracellular space’ and *cp*_5_ : ‘extracellular space’ have the same similarity values when compared to FMA 70022’s preferred label feature, ‘extracellular space’; but intuitively *cp*_5_ should be higher. Therefore, we add the number of terms in the candidate phrase *P* to the normaliser but prevent the excessive similarity value decrease of a longer candidate phrase with 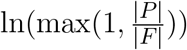 (Equation (2)). Assuming that candidate phrases are always short, we can set the appearance of a term *t* in the candidate phrase to 1, so Equation (2) can be reduced to Equation (3).

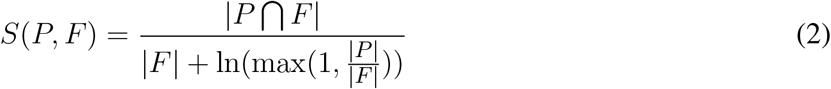

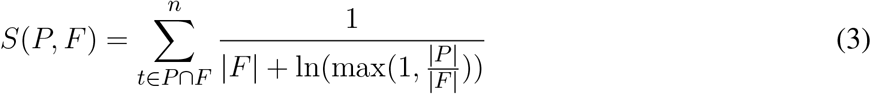

So far all terms in the candidate phrase get the same weight, while based on the level of dependence, different terms ideally have different weights. For example, ‘potassium’ in *cp*_2_: ‘concentration of potassium’ should have lower weight than in *cp*_4_: ‘potassium’, because in *cp*_2_ ‘potassium’ is not the main term and a noun modifier of ‘concentration’, while in *cp*_4_ ‘potassium’ is the main term. Considering that the universal dependencies of the candidate phrase can be constructed as a tree structure, we use the depth of a node plus 1 as the dependency level of a term, so a lower dependency level contributes more to the similarity value. In Figure 3, we can see the example of dependency level determination of *cp*_2_: ‘concentration of potassium’ to 1, 3, 2 and *cpt*_3_: ‘concentration of simple chemical’ to 1, 3, 3, 2. However, when the dependency level of a term in the candidate phrase *dl_tp_* differs from that in the ontology class feature *dl_tf_*, the highest value is chosen. Thus, we introduce the dependency level of a term into Equation (3) so that it becomes Equation (4), where max(*dl_tp_*, *dl_tf_*) selects the highest dependency level.

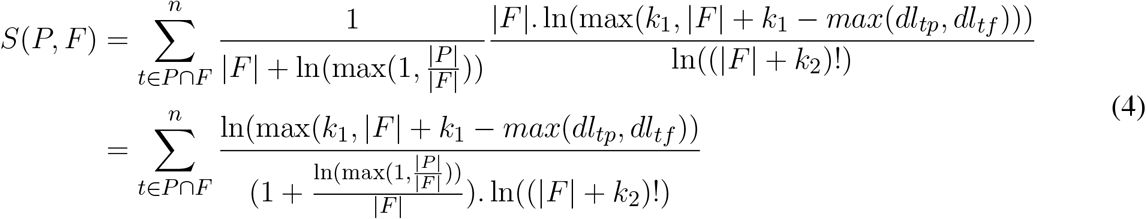

Next, 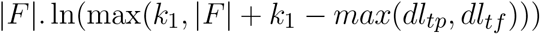 calculates the contribution of the dependency level to the feature’s similarity value, where *k*_1_ is an empirical variable with minimum value of 2. Here, we ensure that a term with a lower dependency level higher than *k*_1_ still has a contribution. Then, this contribution is normalised with ln((|*F*| + *k*_2_)!) where *k*_2_ is an empirical variable with minimal value of 1. For our experiment, *k*_1_ and *k*_2_ were set to 2 and 1, respectively.

For the description feature extracted from the biosimulation model, we use a different similarity calculation because the number of terms in this feature is usually larger than for the other features (Equation (5)). The similarity value is the proportion of overlapping phrases in the candidate phrase and the description feature normalised by the smoothing length of the candidate phrase (1 + ln(|*P*|)) and the smoothing of the total number of terms in the entity descriptions appearing with the ontology class (1 + ln(1 + *ne*_*a*_)). We assume that the more an ontology class is used to annotate entities, the more important that ontology class is. So we multiply by (1 + ln(1 + *ne*_*o*_)), which represents the number of entities annotated by the ontology class.

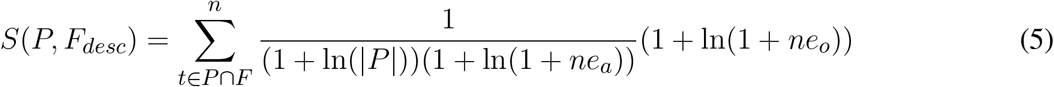

Finally, we combine all features from the similarity results in Equation (6) starting from the preferred label, synonym, definition, parent label, and description in sequence to get the degree of association.

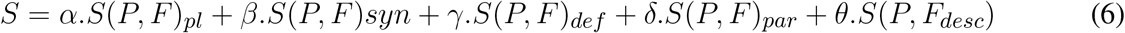

We apply multiple weighting scenarios (Ogilvie et al., 2003; Robertson et al., 2004), so we have multipliers *α*, *β*, *γ*, *δ*, and *θ* for all features. The multipliers are decided empirically to obtain the best performance and will vary based on the training data. At this point, a candidate phrase may relate to multiple ontology classes with various degrees of associations. To get the final phrase, we remove ontology classes with a degree of association lower than a particular threshold value *t*, resulting in candidate phrases with zero or more ontology classes. Hence, the final phrases are those having at least one association with an ontology class.

### 2.4 SPARQL Generator

With the NLQ Annotator result consisting of phrases and their associated ontology classes and predicates, our SPARQL Generator generates SPARQL utilising the RDF Graph Index (RGI) and SPARQL Composer. RGI is an indexing system used to represent entities’ semantic annotation patterns in biosimulation models, where SPARQL Composer is a compiler that locates SPARQL elements in RGI, then constructs and ranks SPARQL.

RGI stores all existing semantic annotation patterns in the repository. For example, the annotated entity in Figure 2 has a pattern consisting of the entities ‘#K_e’ as root described by the ontology classes OPB 00340, FMA_70022, and CHEBI_29103. Between the root and the ontology class, some predicates make up the path. With the ontology classes as the RGI input, we can get the annotation pattern; and then construct SPARQL. This pattern search is done using the indexing system in RGI, which can search for predicate and root sets based on ontology classes (see Supplementary Figure S1 for the RGI creation process).

Each phrase selected by the NLQ Annotator can be associated with multiple ontology classes. For example, in the query we have used throughout this paper, *cp*_3_: ‘concentration’, *cp*_4_: ‘potassium’, and *cp*_5_: ‘extracellular space’ are associated with 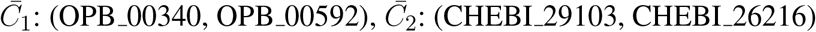, and 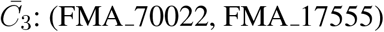, respectively (Figure 4A). The SPARQL Composer initially searches for the available 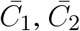, and 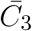 combinations in the RDF Graph by leveraging the RGI’s ontology class to root index. Of the eight combinations, only two have patterns, *ac*_1_: (OPB 00340, FMA 70022, CHEBI 29103) and *ac*_2_: (OPB 00592, FMA 70022, CHEBI 29103) (Figure 4B). The other six possibilities have no matching results in the repositories, and thus, they are discarded. Then, each *ac* is searched for the relationship pattern between its ontology class and root using the RGI ontology class and root to the path. In Figure 4C, we show that *ac*_1_ and *ac*_2_ have one and three relationship patterns, respectively. Next, these patterns are compiled into SPARQL and ranked by summing the degree of phrase association to the ontology class (Figure 4D). Additionally, when using xStanza, the predicate context of a phrase is also used in the ranking. In our example, if the NLQ has predicate terms such as ‘as sink participant’, this can be used to calculate the ranking more accurately.

**Figure 4.**
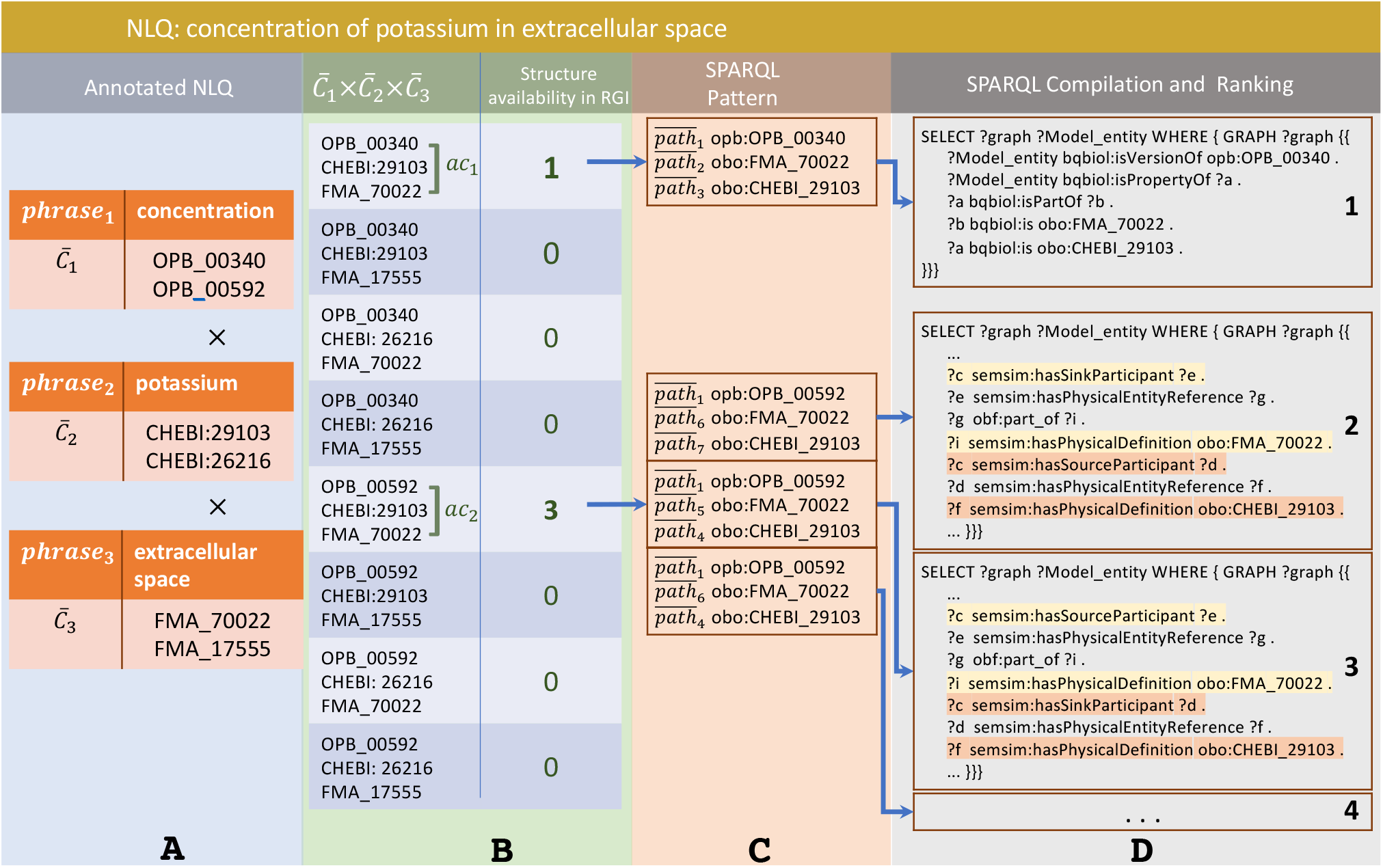
The example of SPARQL generation in SPARQL Composer where (A) phrases and ontology classes as NLQ Annotator result, (B) are combined and checked for their availability in RGI, (C) then are related to available entity annotation patterns, (D) and finally are compiled to SPARQL and ranked.

Arguably, entities can be discovered without constructing SPARQL while their rank can be calculated. However, we still need to verify their availability in repositories along with new entities matching the SPARQL query but not currently indexed in NLIMED.

## 3 EXPERIMENTS AND RESULTS

We performed experiments to measure the performance of our NLQ Annotator in detecting NLQ-related phrases and ontology classes, and to examine NLIMED’s behaviour towards historical query records in the PMR. For the first experiment, we prepared a test dataset examined by experts consisting of 52 NLQ and their associated ontology classes with differing complexity, having 1 – 14 terms and 1 – 4 phrases (Supplementary Table S1). In the second experiment, we collected query-result pairs of from search sessions in the PMR query logs. We obtained 110 pairs by selecting those whose results were models annotated against ontology classes (Supplementary Table S2). An entity identified by NLIMED was considered a relevant result if a model in these pairs contains the entity.

While the preferred label feature is available in all ontology classes, other features may not be present. These empty features may lead to lower degrees of association between a phrase and an ontology class; therefore, adding the preferred label to other features can provide a fairer result. We found that this addition (WPL), along with the use of dependency level to calculate similarity, works well with NLIMED. Furthermore, we varied the measurement on four parsers, i.e. CoreNLP, Benepar, Stanza, xStanza (Stanza with context identification). If a phrase was associated with more than one ontology class, the class with the highest degree of association was selected. We also examined different feature modifications and dependency level implementations more closely. Detailed results are presented in Supplementary Figure S2.

### 3.1 NLQ Annotator performance

The NLQ Annotator performance was measured by calculating precision, recall, and F-measure. Precision is the proportion of correctly identified ontology classes among the number of detected ontology classes, while recall is the proportion of correctly identified classes among the number of ontology classes that count as correct classes. F-measure is the harmonisation of precision and recall obtained by dividing the multiplication of precision and recall by the addition of precision and recall and then multiplying by two. As the gold standard, parsing using the NCBO Annotator is fixed on seven ontologies primarily found in the PMR and prioritised the most prolonged-phrase, resulting in a precision of 0.542 and recall of 0.504.

Table 2 shows experimental results where our approach outperforms the NCBO Annotator. CoreNLP can chunk better than other approaches; moreover, with context identification, xStanza outperforms Stanza by adding new ontology classes, placing this approach second after CoreNLP in recall and F-measure. Furthermore, xStanza provides the ability to detect predicates useful to rank and compose SPARQL. The use of dependency levels and WPL (the distribution of preferred label to other features) can increase the overall performance (see Supplementary Figure S2 with higher *AuC*_*PR*_). Here, dependency levels can improve similarity measurement by giving higher weight to the more critical term. At the same time, WPL can overcome the empty feature problem and balance the participation of each feature by distributing the preferred label, as the essential feature, to other features. As a reference, our most reliable configuration is wpl, mode_3, CoreNLP with multipliers *α* = 3.0, *β* = 3.0, *γ* = 0.0, *δ* = 0.0, and *θ* = 0.38. The complete configurations are available in Supplementary Table S3.

**Table 2.** The performance of NLIMED to annotate NLQ on a test dataset containing 52 NLQs. We modify features by distributing preferred label to other features (WPL). The use of CoreNLP demonstrates the best precision, recall and F-measure. The use of term dependency level in similarity measure can increase recall and F-measure.

| Strategy | Precision | Recall | F-measure |
| --- | --- | --- | --- |
| NCBO Annotator | 0.542 | 0.504 | 0.522 |
| WPL + NoDep + benepar | 0.69 | 0.426 | 0.527 |
| WPL + NoDep + coreNLP | **0.721** | 0.539 | 0.617 |
| WPL + NoDep + stanza | 0.553 | 0.452 | 0.498 |
| WPL + NoDep + xStanza | 0.624 | 0.548 | 0.583 |
| WPL + Dep + benepar | 0.636 | 0.426 | 0.51 |
| WPL + Dep + coreNLP | 0.65 | **0.661** | **0.655** |
| WPL + Dep + stanza | 0.635 | 0.53 | 0.578 |
| WPL + Dep + xStanza | 0.615 | 0.557 | 0.584 |

Considering the NLQ complexity, Figure 5A and 5B present the F-measure for different NLQ lengths. All parsers generally work better than the NCBO Annotator, where CoreNLP is consistently superior to other parsers, and xStanza follows. The analysis based on the number of phrases shows that xStanza is best for one phrase NLQ while CoreNLP is best for the longer NLQ (Figure 5A). The higher performance of CoreNLP is mainly due to its better ability to chunk NLQ, while xStanza’s performance depends on the datasets used to identify named entities. For example, the NLQ ‘luminal antiporter activity’ ideally comprises the two phrases ‘luminal’ and ‘transporter activity’ correctly identified by CoreNLP. In contrast, xStanza identifies ‘luminal’(entity: multi-network structure) or ‘luminal antiporter activity’ (entity: problem). Since the implementation of Stanza avoids any overlapping, ‘luminal’ with the higher degree of association to an ontology class is selected while ‘luminal antiporter activity’ is discarded. The analysis based on the number of terms shows a similar pattern as the analysis based on the number of phrases (Figure 5B), although xStanza is best for two-term NLQ. In general, the high performance of NLIMED on short NLQ shows that the Equation (6) used to calculate the degree of association between a phrase and an ontology class is working well. Hence, NLIMED performance is available to be improved by implementing a better chunker.

**Figure 5.**
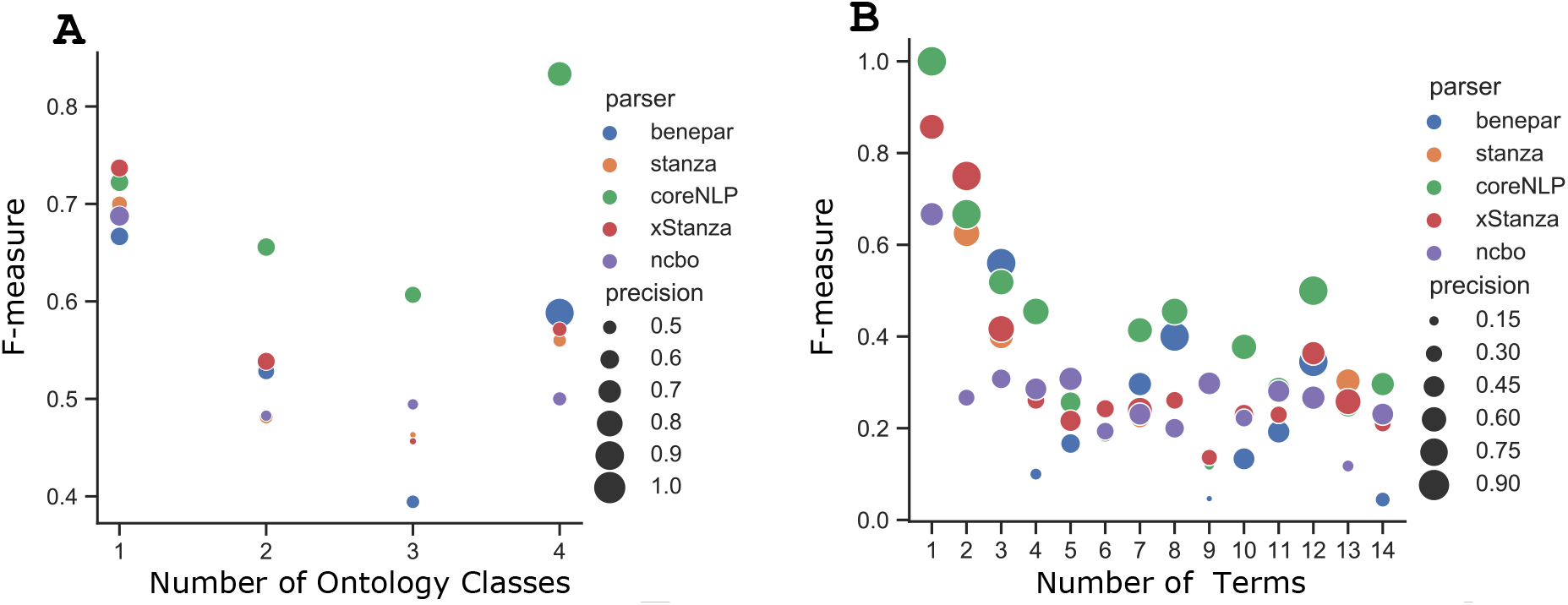
The analysis of NLIMED performance over different number of terms and phrases in NLQ (A) F-measure based on the number of terms in NLQ. (B) F-measure based on the number of phrases in NLQ.

Considering the role of features, the preferred label is the essential feature, and its use with the synonym feature using WPL and CoreNLP could reach higher performance (Supplementary Figure S2A, row 2, column 3). The importance of other features is presented in Supplementary Figure S3 where synonym and description contribute positively for the higher *AuC*_*PR*_.

### 3.2 NLIMED behaviour for historical query records in the PMR

The data used in this experiment has the characteristic that the terms in the ontology classes describing a model are rarely found in the query. It is natural since the available PMR search tools are intended to find biosimulation models rather than entities; therefore, queries tend to use common terms in model descriptions such as author, year, and publication title rather than specific terms in ontology classes. Consequently, NLIMED’s ability to find relevant results based on the provided queries is relatively low. We divided query-result pairs into three groups based on the number of terms that appear together in the query and the results ontology classes divided by the number of terms in the ontology classes. Those groups are G1:”proportion=0”, G2:”0<proportion<=0.5”, and G3:”0.5<proportion<=1.0”. The experiment was conducted using the best multipliers combination stated at Subsection 3.1 and two additional combinations of *α*, *β*, *γ*, *δ*, and *θ* as (3.0, 3.0, 0.5, 0.5, 0.5) and (3.0, 3.0, 1.0, 1.0, 1.0). Using mAP@10 (Mean Average Precision at 10) as a performance measurement, in Supplementary Figure S5, the mAP@10 value continuously increases from G1 to G3. The increase is as expected that a query containing terms in the result’s ontology classes is an advanced query that can find models more precisely. Moreover, in G1, NLIMED can still retrieve a small number of relevant results due to using the entity’s description as a feature.

Contradicting the finding in 3.1, Benepar outperformed other parsers in terms of retrieval, raising F-measure around 0.7 for G3. It seems that Benepar benefits from the relatively higher precision (shown by bigger circles in Figure 5A and 5B), so it can suppress false positives while the number of retrievals is limited. The low performance of CoreNLP and Stanza is probably due to their better ability to identify terms related to ontology classes so that in the chunking process, many common terms are discarded from the query. These common terms help find models that are more general than entities. Meanwhile, the additional ontology classes and predicates by xStanza seem unrelated to the query’s intent, so relevant models found using Stanza have a lower ranking.

## 4 DISCUSSION

### 4.1 NLQ to SPARQL evaluation

We have shown that NLIMED can translate NLQ to SPARQL and find entities annotated to ontology classes. The degree of association equation proposed in the NLQ Annotator combined with the corresponding parser outperformed the NCBO Annotator in terms of associating NLQ with ontology classes. NLIMED performance varies based on the number of terms or the number of phrases in the query (Figure 5), so for future study, it is necessary to consider this query length factor. Next, we discuss NLIMED based on the length and the type of NLQ.

#### 4.1.1 Short NLQ (1-2 phrases)

Most queries have terms that can be annotated correctly, such as ‘chloride’, ‘cardiac myocyte’, ‘apical plasma membrane’, ‘left ventricular wall’, and ‘flux of potassium’. The use of a different parser has its own character on the annotation result; for example, ‘flux of potassium’ is chunked to ‘flux’ and ‘potassium’ by CoreNLP but to ‘flux of simple chemical’ and ‘potassium’ by xStanza, although they are annotated to the same ontology classes, OPB 00593 and CHEBI 29103, respectively. For the same NLQ, Stanza skips the ‘flux’ related phrase because of the lack of a named entity dataset related to the Ontology of Physics for Biology. The use of the NCBO Annotator has a similar issue to Stanza by ignoring some phrases that have a low degree of association to ontology classes. Thus, using information from the annotated entities such as textual description as an additional feature in NLIMED may improve performance.

Nevertheless, recognising phrases not linked by prepositions is quite challenging, for example, noticing that ‘potassium flux’ is similar to ‘flux of potassium’. CoreNLP and Benepar consider it a single phrase that is conceptually correct but does not match the intent of the NLQ, while Stanza only considers the ‘potassium’ term. Extending Stanza, xStanza extracts the context around ‘potassium’ as an additional phrase and successfully associates it as an ontology class. Further, xStanza also identifies additional ontology classes and predicates so that it can construct more specific SPARQL.

#### 4.1.2 Long NLQ (3-4 phrases)

NLQ Annotator’s performance, especially using CoreNLP, is better than NCBO Annotator for NLQ with three and four phrases. Long NLQ are generally well-structured so that determining phrases and ontology classes can be relatively accurate. However, possible drawbacks can occur when there are a lack or excess of identified NLQ phrases. The lack of phrases might not be critical as they only lead to the more general SPARQL, whereas in reality, entities are rarely annotated against more than three ontology classes, so the lack of one ontology class can still generate quite specific SPARQL. On the other hand, the excess of phrases is a critical problem that leads to more specific SPARQL causing NLIMED to miss entities. To overcome this problem, NLIMED implements a filtering threshold based on the degree of association between the phrase and the ontology class.

#### 4.1.3 Question type NLQ

A question type NLQ such as ‘give me models containing a glucose transporter’ usually consists of supporting phrases that are not related to ontology concepts and some relevant phrases which are the core of the NLQ. While Benepar and CoreNLP cannot differentiate accurately, Stanza and xStanza can extract the relevant phrases only as long as the phrase is associated with the available named entity. For the example, Stanza and xStanza can identify ‘glucose transporter’ only, but Benepar and CoreNLP identify an additional phrase ‘give me models’. Nevertheless, for our purposes, since ‘give me models’ contains ‘models’ term which is frequent in biosimulation models, we may consider terms in this phrase as stop words and ignore the phrase. Alternatively, to prevent the same problem, we may apply an inverse document frequency threshold to determine whether to ignore a candidate phrase or not. However, in the future, we may use these non-biological phrases containing terms such as ‘model’, ‘variable’, and ‘constant’ to identify the type of entity to present more accurate results.

### 4.2 NLIMED embedded within other tools

Currently, NLIMED is working over CellML models in the PMR (Yu et al., 2011) and SBML models in BioModels (Chelliah et al., 2015), and has been implemented in EMP (Sarwar et al., 2019a) as an additional interface for discovering model entities. We are confident that NLIMED may also be developed for use with different repositories, e.g. ChEMBL (Gaulton et al., 2012) and BioSamples (Barrett et al., 2012), since they contain models richly annotated with the RDF utilising ontologies. NLIMED has potential to be applied to EMP-like tools such as the Cardiac Electrophysiology Web Lab (Cooper et al., 2016), eSolv (de Boer et al., 2017), and SemGen (Neal et al., 2018). Present annotation tools, e.g. SemGen (Neal et al., 2018), OpenCOR (Garny and Hunter, 2015), and Saint (Lister et al., 2009), which can provide ontology class suggestions based on available ontology databases may take advantage of NLIMED. Following the COMBINE recommendation about standardisation of biosimulation model annotation (Neal et al., 2019), this work can be directed to provide a comprehensive search interface to discover model entities from various biosimulation model repositories.

### 4.3 Limitations and future directions

There are several limitations of NLIMED, some of which will direct our future work. While NLIMED can identify phrases in NLQ and then classify them into ontology classes, we have not yet explored the lexical-semantics; consequently, we cannot differentiate between ‘the role of potassium within cytosol’ and ‘the regulation of K+ in liquid medium contained within a cell’. We predict that the lexical-semantic in NLQ is closely related to the model entity’s tree structure and the dependency between ontology classes.

Currently, NLIMED calculates ranking for all generated SPARQL but not yet for the model entities returned from SPARQL queries. Ranking model entities is quite tricky because all entities retrieved using the same SPARQL have the same ranking and are associated with the same ontology classes, so they do not have differentiation properties. However, the use of the text in the original publication related to the model entities might be helpful, so it may be possible to implement term-weighting strategies such as BM25 (Robertson and Walker, 1994). Furthermore, other entities that surround an entity may also be used to supplement the ranking.

## 5 CONCLUSION

We demonstrated NLIMED, an interface for translating NLQ into SPARQL that consists of NLQ Annotator and SPARQL Generator modules, for model entity discovery. The NLQ Annotator can identify ontology classes in NLQ utilising features extracted from ontologies (preferred label, synonym, definition, and parent label) and biosimulation models’ RDF (description). The ontology class identification performance is relatively high, reaching *AuC*_*PR*_ of 0.690 when using the CoreNLP parser and term dependency level. We also showed that NLIMED could handle a wide range of NLQ types containing one or many terms with one or many phrases. Our SPARQL Generator uses the RDF graphs as indexes into the repositories, and then can generate all possible SPARQL queries (those that have results) based on the provided ontology classes. NLIMED has been implemented in EMP for model entity searching and could potentially be applied as a generic search interface for exploring model entities from numerous biosimulation model repositories, e.g. PMR and BioModels. Further, we are interested in exploring lexical-semantic inside NLQ and semantic concept inside model entities and ontology classes to improve NLIMED performance. By allowing users to easily perform sophisticated search queries over model repositories, we believe that NLIMED will be a valuable model-discovery tool for the biosimulation model community.

## Supporting information

NLIMED Supplementary Material

## AUTHOR CONTRIBUTIONS

YM, KA, and DN contributed to the conception and design of the study. YM implemented NLIMED and performed the analyses. DS embedded NLIMED in an external project. All authors contributed to the testing and evaluation of NLIMED. YM wrote the first draft of the manuscript. YM, AR, and DN wrote sections of the manuscript. All authors contributed to manuscript revision, read, and approved submitted version.

## ACKNOWLEDGMENTS

The authors acknowledge financial support by the Aotearoa Foundation and Auckland Bioengineering Institute.

## DATA AVAILABILITY STATEMENT

The source code, experiment setups and datasets generated for this study can be found in https://github.com/napakalas/NLIMED

