## Supplementary material for "NLIMED: Natural Language Interface for Model Entity Discovery in Biosimulation Model Repositories": NLIMED Supplementary Material

**Table S1:** Test Data: annotation of NLQ to ontology classes in the PMR

| # | Query | Phrases | Annotation |
|---|---|---|---|
| 1 | sodium | sodium | http://purl.obolibrary.org/obo/CHEBI_29101 |
| 2 | chloride | chloride | http://purl.obolibrary.org/obo/CHEBI_17996 |
| 3 | hydron | hydron | http://purl.obolibrary.org/obo/CHEBI_15378 |
| 4 | basolateral membrane | basolateral membrane | http://purl.obolibrary.org/obo/FMA_84669 |
| 5 | apical plasma membrane | apical plasma membrane | http://purl.obolibrary.org/obo/FMA_84666 |
| 6 | Portion of renal filtrate | portion of renal filtrate | http://purl.obolibrary.org/obo/FMA_280587 |
| 7 | Collecting duct of renal tubule | collecting duct of renal tubule | http://purl.obolibrary.org/obo/FMA_15628 |
| 8 | Epithelial cell of proximal tubule | epithelial cell of proximal tubule | http://purl.obolibrary.org/obo/FMA_70973 |
| 9 | sodium/glucose cotransporter 1 (rat) | sodium/glucose cotransporter 1 (rat) | http://purl.obolibrary.org/obo/PR_P53790 |
| 10 | give me model containing glucose transporter | glucose transporter | http://identifiers.org/go/GO:0005355 |
| 11 | chemical concentration flow rate | chemical concentration flow rate | http://identifiers.org/opb/OPB_00593 |
| 12 | voltage gated calcium channel complex | voltage gated calcium channel complex | http://identifiers.org/go/GO:0005891 |
| 13 | van't Hoff law | van't hoff law | http://identifiers.org/opb/OPB_01061 |
| 14 | left ventricular wall | left ventricular wall | http://purl.obolibrary.org/obo/FMA_9556 |
| 15 | oxidative phosphorylation | oxidative phosphorylation | http://identifiers.org/go/GO:0006119 |
| 16 | Cardiac myocyte | cardiac myocyte | http://purl.obolibrary.org/obo/FMA_14067 |
| 17 | calcium driven ADP phosphorylation | calcium | http://identifiers.org/chebi/CHEBI:29108 |
| | | ADP phosphorylation | http://identifiers.org/go/GO:0006757 |
| 18 | electrical potential difference across blood cell | electrical potential difference | http://identifiers.org/opb/OPB_00506 |
| | | blood cell | http://purl.obolibrary.org/obo/FMA_9670 |
| 19 | mechanisms of SGLT2 inhibitors in lowering cytosolic calcium ion transport | sglt2 inhibitors | http://identifiers.org/chebi/CHEBI:73273 |
| | | cytosolic calcium ion transport | http://identifiers.org/go/GO:0060401 |
| 20 | flux of potassium | flux | http://identifiers.org/opb/OPB_00593 |
| | | potassium | http://purl.obolibrary.org/obo/CHEBI_29103 |
| 21 | concentration of sodium | concentration | http://identifiers.org/opb/OPB_00340 |
| | | sodium | http://purl.obolibrary.org/obo/CHEBI_29101 |
| 22 | Nernst reversal potential of sodium | nernst reversal potential | http://identifiers.org/opb/OPB_01581 |
| | | sodium | http://purl.obolibrary.org/obo/CHEBI_29101 |
| 23 | sodium flux | sodium | http://purl.obolibrary.org/obo/CHEBI_29101 |
| | | flux | http://identifiers.org/opb/OPB_00593 |
| 24 | potassium concentration | potassium | http://purl.obolibrary.org/obo/CHEBI_29103 |
| | | concentration | http://identifiers.org/opb/OPB_00340 |
| 25 | luminal antiporter activity | luminal | http://purl.obolibrary.org/obo/FMA_74550 |
| | | antiporter activity | http://identifiers.org/go/GO:0015297 |
| 26 | electrical flow rate incorporating calcium efflux ATPase | electrical flow rate | http://identifiers.org/opb/OPB_00318 |
| | | calcium efflux ATPase | http://identifiers.org/go/GO:0005388 |
| 27 | the mechanisms of calcium channel blocker in ventricular cardiac muscle cell | calcium channel blocker | http://identifiers.org/chebi/CHEBI:38215 |
| | | ventricular cardiac muscle cell | https://identifiers.org/cl/CL:2000046 |

| | | | |
|---|---|---|---|
| 28 | sodium channel in renal intercalated cell | sodium channel | http://purl.obolibrary.org/obo/GO_0005272 |
| | | renal intercalated cell | http://identifiers.org/cl/CL:0005010 |
| 29 | calcium ion significance in mitochondria | calcium ion | http://purl.obolibrary.org/obo/CHEBI_39124 |
| | | mitochondria | http://identifiers.org/go/GO:0005739 |
| 30 | the regulation of mitochondria Na+/Ca2+ antiporter across plasma membrane | the regulation of mitochondria Na+/Ca2+ antiporter | http://identifiers.org/go/GO:0005432 |
| | | plasma membrane | http://purl.obolibrary.org/obo/FMA_63841 |
| 31 | membrane hyperpolarization removes calcium channel blocker | membrane hyperpolarization | http://identifiers.org/go/GO:0060081 |
| | | calcium channel blocker | http://identifiers.org/chebi/CHEBI:38215 |
| 32 | portion of cytosol in epithelial cell of distal tubule via sodium/hydrogen exchanger 3 (human) | portion of cytosol | http://purl.obolibrary.org/obo/FMA_66836 |
| | | epithelial cell of distal tubule | http://purl.obolibrary.org/obo/FMA_70981 |
| | | sodium/hydrogen exchanger 3 (human) | http://purl.obolibrary.org/obo/PR_P48764 |
| 33 | glucose transporter from a kidney that counts for diabetes | glucose transporter | http://identifiers.org/go/GO:0005355 |
| | | kidney | http://purl.obolibrary.org/obo/FMA_7203 |
| | | diabetes | http://identifiers.org/chebi/CHEBI:5931 |
| 34 | Flux of sodium through basolateral plasma membrane | flux | http://identifiers.org/opb/OPB_00593 |
| | | sodium | http://purl.obolibrary.org/obo/CHEBI_29101 |
| | | basolateral plasma membrane | http://purl.obolibrary.org/obo/FMA_84669 |
| 35 | The sodium flow across the apical plasma membrane | sodium | http://purl.obolibrary.org/obo/CHEBI_29101 |
| | | flow | http://identifiers.org/opb/OPB_00593 |
| | | apical plasma membrane | http://purl.obolibrary.org/obo/FMA_84666 |
| 36 | The exchange of chloride with bicarbonate across plasma membrane | chloride | http://purl.obolibrary.org/obo/CHEBI_17996 |
| | | bicarbonate | http://purl.obolibrary.org/obo/CHEBI_17544 |
| | | plasma membrane | http://purl.obolibrary.org/obo/FMA_63841 |
| 37 | concentration of hydron in epithelial cell of proximal tubule compartment | concentration | http://identifiers.org/opb/OPB_00340 |
| | | hydron | http://purl.obolibrary.org/obo/CHEBI_15378 |
| | | epithelial cell of proximal tubule compartment | http://purl.obolibrary.org/obo/FMA_70973 |
| 38 | NBCe1 transports H+ and HCO3- | NBCe1 | http://purl.obolibrary.org/obo/PR_000015160 |
| | | H+ | http://purl.obolibrary.org/obo/CHEBI_15378 |
| | | HCO3- | http://purl.obolibrary.org/obo/CHEBI_17544 |
| 39 | the reverse mode of SGLT1 applies with of a-methyl-alpha-D-glucopyranoside in apical plasma membrane | SGLT1 | http://purl.obolibrary.org/obo/PR_000015165 |
| | | a-methyl-alpha-D-glucopyranoside | http://identifiers.org/chebi/CHEBI:320061 |
| | | tissue fluid | http://purl.obolibrary.org/obo/FMA_84666 |
| 40 | Na/H exchanger 3 rat and anion exchanger 1 rat in the distal convoluted tubule | Na/H exchanger 3 | http://purl.obolibrary.org/obo/PR_P26433 |
| | | anion exchanger | http://purl.obolibrary.org/obo/PR_P23562 |
| | | distal convoluted tubule | http://identifiers.org/fma/FMA:17721 |
| 41 | calcium channel complex and ryanodine receptor in cardiac myocite | calcium channel complex | http://identifiers.org/go/GO:0034704 |
| | | ryanodine receptor | http://identifiers.org/fma/FMA:62492 |
| | | cardiac myocite | http://purl.obolibrary.org/obo/FMA_14067 |
| 42 | Flow rate of Calcium across sarcoplasmic reticulum membrane | flow rate | http://identifiers.org/opb/OPB_00593 |
| | | calcium | http://purl.obolibrary.org/obo/CHEBI_39124 |
| | | sarcoplasmic reticulum membrane | http://identifiers.org/go/GO:0033017 |
| 43 | RyR2 (human) and IP3 related to sarcoplasmic reticulum calcium (human) | RyR2 (human) | http://purl.obolibrary.org/obo/PR_Q92736 |
| | | IP3 | http://purl.obolibrary.org/obo/CHEBI_16595 |

| | | | |
|---|---|---|---|
| | | sarcoplasmic reticulum calcium (human) | http://purl.obolibrary.org/obo/PR_P16615 |
| 44 | All entities containing Mitochondrial Tricarboxylic Acid Cycle, Oxidative Phosphorylation, and isocitrate dehydrogenase complex | mitochondrial tricarboxylic acid cycle | http://identifiers.org/go/GO:0030062 |
| | | oxidative phosphorylation | http://identifiers.org/go/GO:0006119 |
| | | isocitrate dehydrogenase complex | http://identifiers.org/go/GO:0005962 |
| 45 | The calcium state incorporating L-type calcium channel and ryanodine receptor (human) | calcium | http://identifiers.org/chebi/CHEBI:22984 |
| | | L-type calcium channel | http://identifiers.org/go/GO:0008331 |
| | | ryanodine receptor (human) | http://purl.obolibrary.org/obo/PR_Q92736 |
| 46 | Concentration of H+ in the tissue fluid | concentration | http://identifiers.org/opb/OPB_00340 |
| | | H+ | http://purl.obolibrary.org/obo/CHEBI_15378 |
| | | tissue fluid | http://purl.obolibrary.org/obo/FMA_9673 |
| 47 | flux of sodium across apical plasma membrane of the distal convoluted tubule | flux | http://identifiers.org/opb/OPB_00593 |
| | | sodium | http://purl.obolibrary.org/obo/CHEBI_29101 |
| | | apical plasma membrane | http://purl.obolibrary.org/obo/FMA_84666 |
| | | distal convoluted tubule | http://purl.obolibrary.org/obo/FMA_17721 |
| 48 | flux of sodium across apical plasma membrane of the proximal convoluted tubule | flux | http://identifiers.org/opb/OPB_00593 |
| | | sodium | http://purl.obolibrary.org/obo/CHEBI_29101 |
| | | apical plasma membrane | http://purl.obolibrary.org/obo/FMA_84666 |
| | | proximal convoluted tubule | http://purl.obolibrary.org/obo/FMA_17693 |
| 49 | flux of IP3 receptor through P2Y2 purinoceptor and apical plasma membrane | flux | http://identifiers.org/opb/OPB_00593 |
| | | ip3 receptor | http://purl.obolibrary.org/obo/CHEBI_131186 |
| | | p2y2 purinoceptor | http://purl.obolibrary.org/obo/PR_P35383 |
| | | apical plasma membrane | http://purl.obolibrary.org/obo/FMA_84666 |
| 50 | concentration of sodium in the portion of cytosol in epithelial cell of distal tubule | concentration | http://identifiers.org/opb/OPB_00340 |
| | | sodium | http://purl.obolibrary.org/obo/CHEBI_29101 |
| | | portion of cytosol | http://purl.obolibrary.org/obo/FMA_66836 |
| | | epithelial cell of distal tubule | http://purl.obolibrary.org/obo/FMA_70981 |
| 51 | sodium concentration in tissue fluid in epithelial cell of distal tubule | sodium | http://purl.obolibrary.org/obo/CHEBI_29101 |
| | | concentration | http://identifiers.org/opb/OPB_00340 |
| | | tissue fluid | http://purl.obolibrary.org/obo/FMA_9673 |
| | | epithelial cell of distal tubule | http://purl.obolibrary.org/obo/FMA_70981 |
| 52 | K+ flow through potassium channel complex across apical cell membrane | K+ | http://identifiers.org/chebi/CHEBI:29103 |
| | | flow | http://identifiers.org/opb/OPB_00593 |
| | | potassium channel complex | http://identifiers.org/go/GO:0034705 |
| | | apical cell membrane | http://purl.obolibrary.org/obo/FMA_84666 |

**Table S2:** Historical query and result models in the PMR

| # | Query | Model |
|---|---|---|
| 1 | electrical potential difference human | hodgkin_huxley_1952/rawfile/HEAD/hodgkin_huxley_1952.cellml |
| | | hodgkin_huxley_1952/rawfile/HEAD/hodgkin_huxley_1952_variant01.cellml |
| 2 | electrical potential | baylor_hollingworth_chandler_2002/rawfile/HEAD/baylor_hollingworth_chandler_2002_b.cellml |
| | | baylor_hollingworth_chandler_2002/rawfile/HEAD/baylor_hollingworth_chandler_2002_d.cellml |
| | | luo_rudy_1994/rawfile/HEAD/luo_rudy_1994.cellml |
| | | colegrove_albrecht_friel_2000/rawfile/HEAD/colegrove_albrecht_friel_2000.cellml |
| | | dougherty_wright_yew_2005/rawfile/HEAD/dougherty_wright_yew_2005.cellml |
| | | yamaguchi_takaki_matsubara_yasuhara_suga_1996/rawfile/HEAD/yamaguchi_takaki_matsubara_yasuhara_suga_1996.cellml |

| | | |
|---|---|---|
| | | boyett_zhang_garny_holden_2001/rawfile/HEAD/boyett_zhang_garny_holden_2001.cellml |
| | | iribe_kohl_noble_2006/rawfile/HEAD/iribe_kohl_noble_2006.cellml |
| | | izakov_katsnelson_blyakhman_markhasin_shkylar_1991/rawfile/HEAD/izakov_katsnelson_blyakhman_markhasin_shkylar_1991.cellml |
| | | stern_song_sham_yang_boheler_rios_1999/rawfile/HEAD/stern_song_sham_yang_boheler_rios_1999.cellml |
| | | devries_sherman_2000/rawfile/HEAD/devries_sherman_2000.cellml |
| | | marhl_haberichter_brumen_heinrich_2000/rawfile/HEAD/marhl_haberichter_brumen_heinrich_2000.cellml |
| 3 | heart tissue | tentusscher_noble_noble_panfilov_2004/rawfile/HEAD/tentusscher_noble_noble_panfilov_2004_c.cellml |
| | | tentusscher_noble_noble_panfilov_2004/rawfile/HEAD/tentusscher_noble_noble_panfilov_2004_b.cellml |
| | | tentusscher_noble_noble_panfilov_2004/rawfile/HEAD/tentusscher_noble_noble_panfilov_2004_a.cellml |
| 4 | calmodulin | baylor_hollingworth_chandler_2002/rawfile/HEAD/baylor_hollingworth_chandler_2002_b.cellml |
| | | baylor_hollingworth_chandler_2002/rawfile/HEAD/baylor_hollingworth_chandler_2002_d.cellml |
| | | luo_rudy_1994/rawfile/HEAD/luo_rudy_1994.cellml |
| | | colegrove_albrecht_friel_2000/rawfile/HEAD/colegrove_albrecht_friel_2000.cellml |
| | | dougherty_wright_yew_2005/rawfile/HEAD/dougherty_wright_yew_2005.cellml |
| | | yamaguchi_takaki_matsubara_yasuhara_suga_1996/rawfile/HEAD/yamaguchi_takaki_matsubara_yasuhara_suga_1996.cellml |
| | | boyett_zhang_garny_holden_2001/rawfile/HEAD/boyett_zhang_garny_holden_2001.cellml |
| | | iribe_kohl_noble_2006/rawfile/HEAD/iribe_kohl_noble_2006.cellml |
| | | izakov_katsnelson_blyakhman_markhasin_shkylar_1991/rawfile/HEAD/izakov_katsnelson_blyakhman_markhasin_shkylar_1991.cellml |
| | | stern_song_sham_yang_boheler_rios_1999/rawfile/HEAD/stern_song_sham_yang_boheler_rios_1999.cellml |
| | | devries_sherman_2000/rawfile/HEAD/devries_sherman_2000.cellml |
| | | marhl_haberichter_brumen_heinrich_2000/rawfile/HEAD/marhl_haberichter_brumen_heinrich_2000.cellml |
| 5 | a synthetic oscillatory network of transcriptional | baylor_hollingworth_chandler_2002/rawfile/HEAD/baylor_hollingworth_chandler_2002_b.cellml |
| | | baylor_hollingworth_chandler_2002/rawfile/HEAD/baylor_hollingworth_chandler_2002_d.cellml |
| | | luo_rudy_1994/rawfile/HEAD/luo_rudy_1994.cellml |
| | | colegrove_albrecht_friel_2000/rawfile/HEAD/colegrove_albrecht_friel_2000.cellml |
| | | dougherty_wright_yew_2005/rawfile/HEAD/dougherty_wright_yew_2005.cellml |
| | | yamaguchi_takaki_matsubara_yasuhara_suga_1996/rawfile/HEAD/yamaguchi_takaki_matsubara_yasuhara_suga_1996.cellml |
| | | boyett_zhang_garny_holden_2001/rawfile/HEAD/boyett_zhang_garny_holden_2001.cellml |
| | | iribe_kohl_noble_2006/rawfile/HEAD/iribe_kohl_noble_2006.cellml |
| | | izakov_katsnelson_blyakhman_markhasin_shkylar_1991/rawfile/HEAD/izakov_katsnelson_blyakhman_markhasin_shkylar_1991.cellml |
| | | stern_song_sham_yang_boheler_rios_1999/rawfile/HEAD/stern_song_sham_yang_boheler_rios_1999.cellml |
| | | devries_sherman_2000/rawfile/HEAD/devries_sherman_2000.cellml |
| | | marhl_haberichter_brumen_heinrich_2000/rawfile/HEAD/marhl_haberichter_brumen_heinrich_2000.cellml |
| 6 | heart | baylor_hollingworth_chandler_2002/rawfile/HEAD/baylor_hollingworth_chandler_2002_b.cellml |
| | | baylor_hollingworth_chandler_2002/rawfile/HEAD/baylor_hollingworth_chandler_2002_d.cellml |
| | | luo_rudy_1994/rawfile/HEAD/luo_rudy_1994.cellml |
| | | colegrove_albrecht_friel_2000/rawfile/HEAD/colegrove_albrecht_friel_2000.cellml |
| | | dougherty_wright_yew_2005/rawfile/HEAD/dougherty_wright_yew_2005.cellml |
| | | yamaguchi_takaki_matsubara_yasuhara_suga_1996/rawfile/HEAD/yamaguchi_takaki_matsubara_yasuhara_suga_1996.cellml |
| | | boyett_zhang_garny_holden_2001/rawfile/HEAD/boyett_zhang_garny_holden_2001.cellml |
| | | iribe_kohl_noble_2006/rawfile/HEAD/iribe_kohl_noble_2006.cellml |
| | | izakov_katsnelson_blyakhman_markhasin_shkylar_1991/rawfile/HEAD/izakov_katsnelson_blyakhman_markhasin_shkylar_1991.cellml |
| | | stern_song_sham_yang_boheler_rios_1999/rawfile/HEAD/stern_song_sham_yang_boheler_rios_1999.cellml |
| | | devries_sherman_2000/rawfile/HEAD/devries_sherman_2000.cellml |
| | | marhl_haberichter_brumen_heinrich_2000/rawfile/HEAD/marhl_haberichter_brumen_heinrich_2000.cellml |
| 7 | ion | michailova_mcculloch_2001/rawfile/HEAD/michailova_mcculloch_2001.cellml |
| | | jafri_rice_winslow_1998/rawfile/HEAD/jafri_rice_winslow_1998_a.cellml |
| | | winslow_rice_jafri_marban_ororke_1999/rawfile/HEAD/winslow_rice_jafri_marban_ororke_1999.cellml |
| | | jafri_rice_winslow_1998/rawfile/HEAD/jafri_rice_winslow_1998_b.cellml |
| 8 | minimal haemodynamic system model including ventricular interaction and valve dynamics | baylor_hollingworth_chandler_2002/rawfile/HEAD/baylor_hollingworth_chandler_2002_b.cellml |
| | | baylor_hollingworth_chandler_2002/rawfile/HEAD/baylor_hollingworth_chandler_2002_d.cellml |
| | | luo_rudy_1994/rawfile/HEAD/luo_rudy_1994.cellml |
| | | colegrove_albrecht_friel_2000/rawfile/HEAD/colegrove_albrecht_friel_2000.cellml |
| | | dougherty_wright_yew_2005/rawfile/HEAD/dougherty_wright_yew_2005.cellml |
| | | yamaguchi_takaki_matsubara_yasuhara_suga_1996/rawfile/HEAD/yamaguchi_takaki_matsubara_yasuhara_suga_1996.cellml |
| | | boyett_zhang_garny_holden_2001/rawfile/HEAD/boyett_zhang_garny_holden_2001.cellml |
| | | iribe_kohl_noble_2006/rawfile/HEAD/iribe_kohl_noble_2006.cellml |
| | | izakov_katsnelson_blyakhman_markhasin_shkylar_1991/rawfile/HEAD/izakov_katsnelson_blyakhman_markhasin_shkylar_1991.cellml |
| | | stern_song_sham_yang_boheler_rios_1999/rawfile/HEAD/stern_song_sham_yang_boheler_rios_1999.cellml |
| | | devries_sherman_2000/rawfile/HEAD/devries_sherman_2000.cellml |

| | | |
|---|---|---|
| | | marhl_haberichter_brumen_heinrich_2000/rawfile/HEAD/marhl_haberichter_brumen_heinrich_2000.cellml |
| 9 | diffusion | keener_2001/rawfile/HEAD/keener_2001.cellml |
| 10 | calmodulin mediates differential sensitivity of camkii and calcineurin to local | baylor_hollingworth_chandler_2002/rawfile/HEAD/baylor_hollingworth_chandler_2002_b.cellml |
| | | baylor_hollingworth_chandler_2002/rawfile/HEAD/baylor_hollingworth_chandler_2002_d.cellml |
| | | luo_rudy_1994/rawfile/HEAD/luo_rudy_1994.cellml |
| | | colegrove_albrecht_friel_2000/rawfile/HEAD/colegrove_albrecht_friel_2000.cellml |
| | | dougherty_wright_yew_2005/rawfile/HEAD/dougherty_wright_yew_2005.cellml |
| | | yamaguchi_takaki_matsubara_yasuhara_suga_1996/rawfile/HEAD/yamaguchi_takaki_matsubara_yasuhara_suga_1996.cellml |
| | | boyett_zhang_garny_holden_2001/rawfile/HEAD/boyett_zhang_garny_holden_2001.cellml |
| | | iribe_kohl_noble_2006/rawfile/HEAD/iribe_kohl_noble_2006.cellml |
| | | izakov_katsnelson_blyakhman_markhasin_shkylar_1991/rawfile/HEAD/izakov_katsnelson_blyakhman_markhasin_shkylar_1991.cellml |
| | | stern_song_sham_yang_boheler_rios_1999/rawfile/HEAD/stern_song_sham_yang_boheler_rios_1999.cellml |
| | | devries_sherman_2000/rawfile/HEAD/devries_sherman_2000.cellml |
| | | marhl_haberichter_brumen_heinrich_2000/rawfile/HEAD/marhl_haberichter_brumen_heinrich_2000.cellml |
| 11 | a mathematical treatment of integrated ca dynamics within the ventricular myocyte | shannon_wang_puglisi_weber_bers_2004/rawfile/HEAD/shannon_wang_puglisi_weber_bers_2004_a.cellml |
| | | shannon_wang_puglisi_weber_bers_2004/rawfile/HEAD/shannon_wang_puglisi_weber_bers_2004_b.cellml |
| 12 | human atrial | baylor_hollingworth_chandler_2002/rawfile/HEAD/baylor_hollingworth_chandler_2002_b.cellml |
| | | baylor_hollingworth_chandler_2002/rawfile/HEAD/baylor_hollingworth_chandler_2002_d.cellml |
| | | luo_rudy_1994/rawfile/HEAD/luo_rudy_1994.cellml |
| | | colegrove_albrecht_friel_2000/rawfile/HEAD/colegrove_albrecht_friel_2000.cellml |
| | | dougherty_wright_yew_2005/rawfile/HEAD/dougherty_wright_yew_2005.cellml |
| | | yamaguchi_takaki_matsubara_yasuhara_suga_1996/rawfile/HEAD/yamaguchi_takaki_matsubara_yasuhara_suga_1996.cellml |
| | | boyett_zhang_garny_holden_2001/rawfile/HEAD/boyett_zhang_garny_holden_2001.cellml |
| | | iribe_kohl_noble_2006/rawfile/HEAD/iribe_kohl_noble_2006.cellml |
| | | izakov_katsnelson_blyakhman_markhasin_shkylar_1991/rawfile/HEAD/izakov_katsnelson_blyakhman_markhasin_shkylar_1991.cellml |
| | | stern_song_sham_yang_boheler_rios_1999/rawfile/HEAD/stern_song_sham_yang_boheler_rios_1999.cellml |
| | | devries_sherman_2000/rawfile/HEAD/devries_sherman_2000.cellml |
| | | marhl_haberichter_brumen_heinrich_2000/rawfile/HEAD/marhl_haberichter_brumen_heinrich_2000.cellml |
| 13 | thomas r. shannon | shannon_wang_puglisi_weber_bers_2004/rawfile/HEAD/shannon_wang_puglisi_weber_bers_2004_b.cellml |
| | | shannon_wang_puglisi_weber_bers_2004/rawfile/HEAD/shannon_wang_puglisi_weber_bers_2004_a.cellml |
| 14 | atrial fibrillation | baylor_hollingworth_chandler_2002/rawfile/HEAD/baylor_hollingworth_chandler_2002_b.cellml |
| | | baylor_hollingworth_chandler_2002/rawfile/HEAD/baylor_hollingworth_chandler_2002_d.cellml |
| | | luo_rudy_1994/rawfile/HEAD/luo_rudy_1994.cellml |
| | | colegrove_albrecht_friel_2000/rawfile/HEAD/colegrove_albrecht_friel_2000.cellml |
| | | dougherty_wright_yew_2005/rawfile/HEAD/dougherty_wright_yew_2005.cellml |
| | | yamaguchi_takaki_matsubara_yasuhara_suga_1996/rawfile/HEAD/yamaguchi_takaki_matsubara_yasuhara_suga_1996.cellml |
| | | boyett_zhang_garny_holden_2001/rawfile/HEAD/boyett_zhang_garny_holden_2001.cellml |
| | | iribe_kohl_noble_2006/rawfile/HEAD/iribe_kohl_noble_2006.cellml |
| | | izakov_katsnelson_blyakhman_markhasin_shkylar_1991/rawfile/HEAD/izakov_katsnelson_blyakhman_markhasin_shkylar_1991.cellml |
| | | stern_song_sham_yang_boheler_rios_1999/rawfile/HEAD/stern_song_sham_yang_boheler_rios_1999.cellml |
| | | devries_sherman_2000/rawfile/HEAD/devries_sherman_2000.cellml |
| | | marhl_haberichter_brumen_heinrich_2000/rawfile/HEAD/marhl_haberichter_brumen_heinrich_2000.cellml |
| 15 | nhe3 | 267/rawfile/HEAD/weinstein_1995-rabbit.cellml |
| | | 267/rawfile/HEAD/SEDML/weinstein_1995/weinstein_1995.cellml |
| | | 267/rawfile/HEAD/weinstein_1995-mouse.cellml |
| | | 267/rawfile/HEAD/weinstein_1995.cellml |
| 16 | caffeine concentration model | baylor_hollingworth_chandler_2002/rawfile/HEAD/baylor_hollingworth_chandler_2002_b.cellml |
| | | baylor_hollingworth_chandler_2002/rawfile/HEAD/baylor_hollingworth_chandler_2002_d.cellml |
| | | luo_rudy_1994/rawfile/HEAD/luo_rudy_1994.cellml |
| | | colegrove_albrecht_friel_2000/rawfile/HEAD/colegrove_albrecht_friel_2000.cellml |
| | | dougherty_wright_yew_2005/rawfile/HEAD/dougherty_wright_yew_2005.cellml |
| | | yamaguchi_takaki_matsubara_yasuhara_suga_1996/rawfile/HEAD/yamaguchi_takaki_matsubara_yasuhara_suga_1996.cellml |
| | | boyett_zhang_garny_holden_2001/rawfile/HEAD/boyett_zhang_garny_holden_2001.cellml |
| | | iribe_kohl_noble_2006/rawfile/HEAD/iribe_kohl_noble_2006.cellml |
| | | izakov_katsnelson_blyakhman_markhasin_shkylar_1991/rawfile/HEAD/izakov_katsnelson_blyakhman_markhasin_shkylar_1991.cellml |
| | | stern_song_sham_yang_boheler_rios_1999/rawfile/HEAD/stern_song_sham_yang_boheler_rios_1999.cellml |
| | | devries_sherman_2000/rawfile/HEAD/devries_sherman_2000.cellml |

| | | |
|---|---|---|
| | | marhl_haberichter_brumen_heinrich_2000/rawfile/HEAD/marhl_haberichter_brumen_heinrich_2000.cellml |
| 17 | gap junctions | bindschadler_sneyd_2001/rawfile/HEAD/bindschadler_sneyd_2001.cellml |
| | | 267/rawfile/HEAD/bindschadler_sneyd_2001.cellml |
| 18 | gap junction | baylor_hollingworth_chandler_2002/rawfile/HEAD/baylor_hollingworth_chandler_2002_b.cellml |
| | | baylor_hollingworth_chandler_2002/rawfile/HEAD/baylor_hollingworth_chandler_2002_d.cellml |
| | | luo_rudy_1994/rawfile/HEAD/luo_rudy_1994.cellml |
| | | colegrove_albrecht_friel_2000/rawfile/HEAD/colegrove_albrecht_friel_2000.cellml |
| | | dougherty_wright_yew_2005/rawfile/HEAD/dougherty_wright_yew_2005.cellml |
| | | yamaguchi_takaki_matsubara_yasuhara_suga_1996/rawfile/HEAD/yamaguchi_takaki_matsubara_yasuhara_suga_1996.cellml |
| | | boyett_zhang_garny_holden_2001/rawfile/HEAD/boyett_zhang_garny_holden_2001.cellml |
| | | iribe_kohl_noble_2006/rawfile/HEAD/iribe_kohl_noble_2006.cellml |
| | | izakov_katsnelson_blyakhman_markhasin_shkylar_1991/rawfile/HEAD/izakov_katsnelson_blyakhman_markhasin_shkylar_1991.cellml |
| | | stern_song_sham_yang_boheler_rios_1999/rawfile/HEAD/stern_song_sham_yang_boheler_rios_1999.cellml |
| | | devries_sherman_2000/rawfile/HEAD/devries_sherman_2000.cellml |
| | | marhl_haberichter_brumen_heinrich_2000/rawfile/HEAD/marhl_haberichter_brumen_heinrich_2000.cellml |
| 19 | a novel computational model of the human ventricular action potential and ca transient | baylor_hollingworth_chandler_2002/rawfile/HEAD/baylor_hollingworth_chandler_2002_b.cellml |
| | | baylor_hollingworth_chandler_2002/rawfile/HEAD/baylor_hollingworth_chandler_2002_d.cellml |
| | | luo_rudy_1994/rawfile/HEAD/luo_rudy_1994.cellml |
| | | colegrove_albrecht_friel_2000/rawfile/HEAD/colegrove_albrecht_friel_2000.cellml |
| | | dougherty_wright_yew_2005/rawfile/HEAD/dougherty_wright_yew_2005.cellml |
| | | yamaguchi_takaki_matsubara_yasuhara_suga_1996/rawfile/HEAD/yamaguchi_takaki_matsubara_yasuhara_suga_1996.cellml |
| | | boyett_zhang_garny_holden_2001/rawfile/HEAD/boyett_zhang_garny_holden_2001.cellml |
| | | iribe_kohl_noble_2006/rawfile/HEAD/iribe_kohl_noble_2006.cellml |
| | | izakov_katsnelson_blyakhman_markhasin_shkylar_1991/rawfile/HEAD/izakov_katsnelson_blyakhman_markhasin_shkylar_1991.cellml |
| | | stern_song_sham_yang_boheler_rios_1999/rawfile/HEAD/stern_song_sham_yang_boheler_rios_1999.cellml |
| | | devries_sherman_2000/rawfile/HEAD/devries_sherman_2000.cellml |
| | | marhl_haberichter_brumen_heinrich_2000/rawfile/HEAD/marhl_haberichter_brumen_heinrich_2000.cellml |
| 20 | a model for human ventricular tissue | tentusscher_noble_noble_panfilov_2004/rawfile/HEAD/tentusscher_noble_noble_panfilov_2004_c.cellml |
| | | tentusscher_noble_noble_panfilov_2004/rawfile/HEAD/tentusscher_noble_noble_panfilov_2004_b.cellml |
| | | tentusscher_noble_noble_panfilov_2004/rawfile/HEAD/tentusscher_noble_noble_panfilov_2004_a.cellml |
| 21 | a model of the ventricular cardiac action potential: depolarization, repolarization, and their interaction | luo_rudy_1991/rawfile/HEAD/luo_rudy_1991.cellml |
| 22 | cardiomyocyte | baylor_hollingworth_chandler_2002/rawfile/HEAD/baylor_hollingworth_chandler_2002_b.cellml |
| | | baylor_hollingworth_chandler_2002/rawfile/HEAD/baylor_hollingworth_chandler_2002_d.cellml |
| | | luo_rudy_1994/rawfile/HEAD/luo_rudy_1994.cellml |
| | | colegrove_albrecht_friel_2000/rawfile/HEAD/colegrove_albrecht_friel_2000.cellml |
| | | dougherty_wright_yew_2005/rawfile/HEAD/dougherty_wright_yew_2005.cellml |
| | | yamaguchi_takaki_matsubara_yasuhara_suga_1996/rawfile/HEAD/yamaguchi_takaki_matsubara_yasuhara_suga_1996.cellml |
| | | boyett_zhang_garny_holden_2001/rawfile/HEAD/boyett_zhang_garny_holden_2001.cellml |
| | | iribe_kohl_noble_2006/rawfile/HEAD/iribe_kohl_noble_2006.cellml |
| | | izakov_katsnelson_blyakhman_markhasin_shkylar_1991/rawfile/HEAD/izakov_katsnelson_blyakhman_markhasin_shkylar_1991.cellml |
| | | stern_song_sham_yang_boheler_rios_1999/rawfile/HEAD/stern_song_sham_yang_boheler_rios_1999.cellml |
| | | devries_sherman_2000/rawfile/HEAD/devries_sherman_2000.cellml |
| | | marhl_haberichter_brumen_heinrich_2000/rawfile/HEAD/marhl_haberichter_brumen_heinrich_2000.cellml |
| 23 | a mathematical model of plasma membrane electrophysiology and calcium dynamics in vascular endothelial cells python model | baylor_hollingworth_chandler_2002/rawfile/HEAD/baylor_hollingworth_chandler_2002_b.cellml |
| | | baylor_hollingworth_chandler_2002/rawfile/HEAD/baylor_hollingworth_chandler_2002_d.cellml |
| | | luo_rudy_1994/rawfile/HEAD/luo_rudy_1994.cellml |
| | | colegrove_albrecht_friel_2000/rawfile/HEAD/colegrove_albrecht_friel_2000.cellml |
| | | dougherty_wright_yew_2005/rawfile/HEAD/dougherty_wright_yew_2005.cellml |
| | | yamaguchi_takaki_matsubara_yasuhara_suga_1996/rawfile/HEAD/yamaguchi_takaki_matsubara_yasuhara_suga_1996.cellml |
| | | boyett_zhang_garny_holden_2001/rawfile/HEAD/boyett_zhang_garny_holden_2001.cellml |
| | | iribe_kohl_noble_2006/rawfile/HEAD/iribe_kohl_noble_2006.cellml |
| | | izakov_katsnelson_blyakhman_markhasin_shkylar_1991/rawfile/HEAD/izakov_katsnelson_blyakhman_markhasin_shkylar_1991.cellml |
| | | stern_song_sham_yang_boheler_rios_1999/rawfile/HEAD/stern_song_sham_yang_boheler_rios_1999.cellml |
| | | devries_sherman_2000/rawfile/HEAD/devries_sherman_2000.cellml |
| | | marhl_haberichter_brumen_heinrich_2000/rawfile/HEAD/marhl_haberichter_brumen_heinrich_2000.cellml |
| | | baylor_hollingworth_chandler_2002/rawfile/HEAD/baylor_hollingworth_chandler_2002_b.cellml |
| | | baylor_hollingworth_chandler_2002/rawfile/HEAD/baylor_hollingworth_chandler_2002_d.cellml |
| | | luo_rudy_1994/rawfile/HEAD/luo_rudy_1994.cellml |
| 24 | cell cycle | |

| | | |
|---|---|---|
| | | colegrove_albrecht_friel_2000/rawfile/HEAD/colegrove_albrecht_friel_2000.cellml |
| | | dougherty_wright_yew_2005/rawfile/HEAD/dougherty_wright_yew_2005.cellml |
| | | yamaguchi_takaki_matsubara_yasuhara_suga_1996/rawfile/HEAD/yamaguchi_takaki_matsubara_yasuhara_suga_1996.cellml |
| | | boyett_zhang_garny_holden_2001/rawfile/HEAD/boyett_zhang_garny_holden_2001.cellml |
| | | iribe_kohl_noble_2006/rawfile/HEAD/iribe_kohl_noble_2006.cellml |
| | | izakov_katsnelson_blyakhman_markhasin_shkylar_1991/rawfile/HEAD/izakov_katsnelson_blyakhman_markhasin_shkylar_1991.cellml |
| | | stern_song_sham_yang_boheler_rios_1999/rawfile/HEAD/stern_song_sham_yang_boheler_rios_1999.cellml |
| | | devries_sherman_2000/rawfile/HEAD/devries_sherman_2000.cellml |
| | | marhl_haberichter_brumen_heinrich_2000/rawfile/HEAD/marhl_haberichter_brumen_heinrich_2000.cellml |
| 25 | using physiome standards to couple cellular functions for rat cardiac excitation-contraction | terkildsen_niederer_crampin_hunter_smith_2008/rawfile/HEAD/Pandit_Hinch_Niederer.cellml |
| 26 | rat gattoni | terkildsen_niederer_crampin_hunter_smith_2008/rawfile/HEAD/Pandit_Hinch_Niederer.cellml |
| | | baylor_hollingworth_chandler_2002/rawfile/HEAD/baylor_hollingworth_chandler_2002_b.cellml |
| | | baylor_hollingworth_chandler_2002/rawfile/HEAD/baylor_hollingworth_chandler_2002_d.cellml |
| | | luo_rudy_1994/rawfile/HEAD/luo_rudy_1994.cellml |
| | | colegrove_albrecht_friel_2000/rawfile/HEAD/colegrove_albrecht_friel_2000.cellml |
| | | dougherty_wright_yew_2005/rawfile/HEAD/dougherty_wright_yew_2005.cellml |
| | | yamaguchi_takaki_matsubara_yasuhara_suga_1996/rawfile/HEAD/yamaguchi_takaki_matsubara_yasuhara_suga_1996.cellml |
| 27 | action potential | luo_rudy_1991/rawfile/HEAD/luo_rudy_1991.cellml |
| | | boyett_zhang_garny_holden_2001/rawfile/HEAD/boyett_zhang_garny_holden_2001.cellml |
| | | iribe_kohl_noble_2006/rawfile/HEAD/iribe_kohl_noble_2006.cellml |
| | | izakov_katsnelson_blyakhman_markhasin_shkylar_1991/rawfile/HEAD/izakov_katsnelson_blyakhman_markhasin_shkylar_1991.cellml |
| | | stern_song_sham_yang_boheler_rios_1999/rawfile/HEAD/stern_song_sham_yang_boheler_rios_1999.cellml |
| | | devries_sherman_2000/rawfile/HEAD/devries_sherman_2000.cellml |
| | | marhl_haberichter_brumen_heinrich_2000/rawfile/HEAD/marhl_haberichter_brumen_heinrich_2000.cellml |
| | | baylor_hollingworth_chandler_2002/rawfile/HEAD/baylor_hollingworth_chandler_2002_b.cellml |
| | | baylor_hollingworth_chandler_2002/rawfile/HEAD/baylor_hollingworth_chandler_2002_d.cellml |
| | | luo_rudy_1994/rawfile/HEAD/luo_rudy_1994.cellml |
| | | colegrove_albrecht_friel_2000/rawfile/HEAD/colegrove_albrecht_friel_2000.cellml |
| | | dougherty_wright_yew_2005/rawfile/HEAD/dougherty_wright_yew_2005.cellml |
| | | yamaguchi_takaki_matsubara_yasuhara_suga_1996/rawfile/HEAD/yamaguchi_takaki_matsubara_yasuhara_suga_1996.cellml |
| 28 | human fibroblast | boyett_zhang_garny_holden_2001/rawfile/HEAD/boyett_zhang_garny_holden_2001.cellml |
| | | iribe_kohl_noble_2006/rawfile/HEAD/iribe_kohl_noble_2006.cellml |
| | | izakov_katsnelson_blyakhman_markhasin_shkylar_1991/rawfile/HEAD/izakov_katsnelson_blyakhman_markhasin_shkylar_1991.cellml |
| | | stern_song_sham_yang_boheler_rios_1999/rawfile/HEAD/stern_song_sham_yang_boheler_rios_1999.cellml |
| | | devries_sherman_2000/rawfile/HEAD/devries_sherman_2000.cellml |
| | | marhl_haberichter_brumen_heinrich_2000/rawfile/HEAD/marhl_haberichter_brumen_heinrich_2000.cellml |
| | | baylor_hollingworth_chandler_2002/rawfile/HEAD/baylor_hollingworth_chandler_2002_b.cellml |
| | | baylor_hollingworth_chandler_2002/rawfile/HEAD/baylor_hollingworth_chandler_2002_d.cellml |
| | | luo_rudy_1994/rawfile/HEAD/luo_rudy_1994.cellml |
| | | colegrove_albrecht_friel_2000/rawfile/HEAD/colegrove_albrecht_friel_2000.cellml |
| | | dougherty_wright_yew_2005/rawfile/HEAD/dougherty_wright_yew_2005.cellml |
| 29 | minimal hemodynamics heart | yamaguchi_takaki_matsubara_yasuhara_suga_1996/rawfile/HEAD/yamaguchi_takaki_matsubara_yasuhara_suga_1996.cellml |
| | | boyett_zhang_garny_holden_2001/rawfile/HEAD/boyett_zhang_garny_holden_2001.cellml |
| | | iribe_kohl_noble_2006/rawfile/HEAD/iribe_kohl_noble_2006.cellml |
| | | izakov_katsnelson_blyakhman_markhasin_shkylar_1991/rawfile/HEAD/izakov_katsnelson_blyakhman_markhasin_shkylar_1991.cellml |
| | | stern_song_sham_yang_boheler_rios_1999/rawfile/HEAD/stern_song_sham_yang_boheler_rios_1999.cellml |
| | | devries_sherman_2000/rawfile/HEAD/devries_sherman_2000.cellml |
| | | marhl_haberichter_brumen_heinrich_2000/rawfile/HEAD/marhl_haberichter_brumen_heinrich_2000.cellml |
| 30 | di francesco-noble | difrancesco_noble_1985/rawfile/HEAD/difrancesco_noble_1985.cellml |
| | | baylor_hollingworth_chandler_2002/rawfile/HEAD/baylor_hollingworth_chandler_2002_b.cellml |
| | | baylor_hollingworth_chandler_2002/rawfile/HEAD/baylor_hollingworth_chandler_2002_d.cellml |
| | | luo_rudy_1994/rawfile/HEAD/luo_rudy_1994.cellml |
| | | colegrove_albrecht_friel_2000/rawfile/HEAD/colegrove_albrecht_friel_2000.cellml |
| | | dougherty_wright_yew_2005/rawfile/HEAD/dougherty_wright_yew_2005.cellml |
| 31 | smith, crampin | yamaguchi_takaki_matsubara_yasuhara_suga_1996/rawfile/HEAD/yamaguchi_takaki_matsubara_yasuhara_suga_1996.cellml |
| | | boyett_zhang_garny_holden_2001/rawfile/HEAD/boyett_zhang_garny_holden_2001.cellml |
| | | iribe_kohl_noble_2006/rawfile/HEAD/iribe_kohl_noble_2006.cellml |
| | | izakov_katsnelson_blyakhman_markhasin_shkylar_1991/rawfile/HEAD/izakov_katsnelson_blyakhman_markhasin_shkylar_1991.cellml |
| | | stern_song_sham_yang_boheler_rios_1999/rawfile/HEAD/stern_song_sham_yang_boheler_rios_1999.cellml |

| | | |
|---|---|---|
| | | devries_sherman_2000/rawfile/HEAD/devries_sherman_2000.cellml |
| | | marhl_haberichter_brumen_heinrich_2000/rawfile/HEAD/marhl_haberichter_brumen_heinrich_2000.cellml |
| 32 | modeling the effects of caffeine on blood pressure | baylor_hollingworth_chandler_2002/rawfile/HEAD/baylor_hollingworth_chandler_2002_b.cellml |
| | | baylor_hollingworth_chandler_2002/rawfile/HEAD/baylor_hollingworth_chandler_2002_d.cellml |
| | | luo_rudy_1994/rawfile/HEAD/luo_rudy_1994.cellml |
| | | colegrove_albrecht_friel_2000/rawfile/HEAD/colegrove_albrecht_friel_2000.cellml |
| | | dougherty_wright_yew_2005/rawfile/HEAD/dougherty_wright_yew_2005.cellml |
| | | yamaguchi_takaki_matsubara_yasuhara_suga_1996/rawfile/HEAD/yamaguchi_takaki_matsubara_yasuhara_suga_1996.cellml |
| | | boyett_zhang_garny_holden_2001/rawfile/HEAD/boyett_zhang_garny_holden_2001.cellml |
| | | iribe_kohl_noble_2006/rawfile/HEAD/iribe_kohl_noble_2006.cellml |
| | | izakov_katsnelson_blyakhman_markhasin_shkylar_1991/rawfile/HEAD/izakov_katsnelson_blyakhman_markhasin_shkylar_1991.cellml |
| | | stern_song_sham_yang_boheler_rios_1999/rawfile/HEAD/stern_song_sham_yang_boheler_rios_1999.cellml |
| | | devries_sherman_2000/rawfile/HEAD/devries_sherman_2000.cellml |
| | | marhl_haberichter_brumen_heinrich_2000/rawfile/HEAD/marhl_haberichter_brumen_heinrich_2000.cellml |
| 33 | cardiac* | baylor_hollingworth_chandler_2002/rawfile/HEAD/baylor_hollingworth_chandler_2002_b.cellml |
| | | baylor_hollingworth_chandler_2002/rawfile/HEAD/baylor_hollingworth_chandler_2002_d.cellml |
| | | luo_rudy_1994/rawfile/HEAD/luo_rudy_1994.cellml |
| | | colegrove_albrecht_friel_2000/rawfile/HEAD/colegrove_albrecht_friel_2000.cellml |
| | | dougherty_wright_yew_2005/rawfile/HEAD/dougherty_wright_yew_2005.cellml |
| | | yamaguchi_takaki_matsubara_yasuhara_suga_1996/rawfile/HEAD/yamaguchi_takaki_matsubara_yasuhara_suga_1996.cellml |
| | | boyett_zhang_garny_holden_2001/rawfile/HEAD/boyett_zhang_garny_holden_2001.cellml |
| | | iribe_kohl_noble_2006/rawfile/HEAD/iribe_kohl_noble_2006.cellml |
| | | izakov_katsnelson_blyakhman_markhasin_shkylar_1991/rawfile/HEAD/izakov_katsnelson_blyakhman_markhasin_shkylar_1991.cellml |
| | | stern_song_sham_yang_boheler_rios_1999/rawfile/HEAD/stern_song_sham_yang_boheler_rios_1999.cellml |
| | | devries_sherman_2000/rawfile/HEAD/devries_sherman_2000.cellml |
| | | marhl_haberichter_brumen_heinrich_2000/rawfile/HEAD/marhl_haberichter_brumen_heinrich_2000.cellml |
| 34 | insulin | baylor_hollingworth_chandler_2002/rawfile/HEAD/baylor_hollingworth_chandler_2002_b.cellml |
| | | baylor_hollingworth_chandler_2002/rawfile/HEAD/baylor_hollingworth_chandler_2002_d.cellml |
| | | luo_rudy_1994/rawfile/HEAD/luo_rudy_1994.cellml |
| | | gall_susa_1999/rawfile/HEAD/gall_susa_1999_b.cellml |
| | | colegrove_albrecht_friel_2000/rawfile/HEAD/colegrove_albrecht_friel_2000.cellml |
| | | dougherty_wright_yew_2005/rawfile/HEAD/dougherty_wright_yew_2005.cellml |
| | | gall_susa_1999/rawfile/HEAD/gall_susa_1999_c.cellml |
| | | yamaguchi_takaki_matsubara_yasuhara_suga_1996/rawfile/HEAD/yamaguchi_takaki_matsubara_yasuhara_suga_1996.cellml |
| | | boyett_zhang_garny_holden_2001/rawfile/HEAD/boyett_zhang_garny_holden_2001.cellml |
| | | chay_1997/rawfile/HEAD/chay_1997.cellml |
| | | 570/rawfile/HEAD/bertram_satin_pedersen_luciani_sherman_2007.cellml |
| | | keener_2001/rawfile/HEAD/keener_2001.cellml |
| | | iribe_kohl_noble_2006/rawfile/HEAD/iribe_kohl_noble_2006.cellml |
| | | magnus_keizer_1998/rawfile/HEAD/magnus_keizer_1998.cellml |
| | | goforth_bertram_khan_zhang_sherman_satin_2002/rawfile/HEAD/goforth_bertram_khan_zhang_sherman_satin_2002.cellml |
| | | gall_susa_1999/rawfile/HEAD/gall_susa_1999_a.cellml |
| | | izakov_katsnelson_blyakhman_markhasin_shkylar_1991/rawfile/HEAD/izakov_katsnelson_blyakhman_markhasin_shkylar_1991.cellml |
| | | stern_song_sham_yang_boheler_rios_1999/rawfile/HEAD/stern_song_sham_yang_boheler_rios_1999.cellml |
| | | devries_sherman_2000/rawfile/HEAD/devries_sherman_2000.cellml |
| | | marhl_haberichter_brumen_heinrich_2000/rawfile/HEAD/marhl_haberichter_brumen_heinrich_2000.cellml |
| 35 | beta cell | bertram_previte_sherman_kinard_satin_2000/rawfile/HEAD/bertram_previte_sherman_kinard_satin_2000_medium.cellml |
| | | bertram_previte_sherman_kinard_satin_2000/rawfile/HEAD/bertram_previte_sherman_kinard_satin_2000_fast.cellml |
| | | bertram_previte_sherman_kinard_satin_2000/rawfile/HEAD/bertram_previte_sherman_kinard_satin_2000_slow.cellml |
| 36 | camkii | baylor_hollingworth_chandler_2002/rawfile/HEAD/baylor_hollingworth_chandler_2002_b.cellml |
| | | baylor_hollingworth_chandler_2002/rawfile/HEAD/baylor_hollingworth_chandler_2002_d.cellml |
| | | luo_rudy_1994/rawfile/HEAD/luo_rudy_1994.cellml |
| | | colegrove_albrecht_friel_2000/rawfile/HEAD/colegrove_albrecht_friel_2000.cellml |
| | | dougherty_wright_yew_2005/rawfile/HEAD/dougherty_wright_yew_2005.cellml |
| | | yamaguchi_takaki_matsubara_yasuhara_suga_1996/rawfile/HEAD/yamaguchi_takaki_matsubara_yasuhara_suga_1996.cellml |
| | | boyett_zhang_garny_holden_2001/rawfile/HEAD/boyett_zhang_garny_holden_2001.cellml |
| | | iribe_kohl_noble_2006/rawfile/HEAD/iribe_kohl_noble_2006.cellml |
| | | izakov_katsnelson_blyakhman_markhasin_shkylar_1991/rawfile/HEAD/izakov_katsnelson_blyakhman_markhasin_shkylar_1991.cellml |
| | | stern_song_sham_yang_boheler_rios_1999/rawfile/HEAD/stern_song_sham_yang_boheler_rios_1999.cellml |
| | | devries_sherman_2000/rawfile/HEAD/devries_sherman_2000.cellml |

| | | |
|---|---|---|
| | | marhl_haberichter_brumen_heinrich_2000/rawfile/HEAD/marhl_haberichter_brumen_heinrich_2000.cellml |
| 37 | frog intact muscle | baylor_hollingworth_chandler_2002/rawfile/HEAD/baylor_hollingworth_chandler_2002_f.cellml |
| 38 | christopher data-driven computer | baylor_hollingworth_chandler_2002/rawfile/HEAD/baylor_hollingworth_chandler_2002_b.cellml |
| | | baylor_hollingworth_chandler_2002/rawfile/HEAD/baylor_hollingworth_chandler_2002_d.cellml |
| | | luo_rudy_1994/rawfile/HEAD/luo_rudy_1994.cellml |
| | | colegrove_albrecht_friel_2000/rawfile/HEAD/colegrove_albrecht_friel_2000.cellml |
| | | dougherty_wright_yew_2005/rawfile/HEAD/dougherty_wright_yew_2005.cellml |
| | | yamaguchi_takaki_matsubara_yasuhara_suga_1996/rawfile/HEAD/yamaguchi_takaki_matsubara_yasuhara_suga_1996.cellml |
| | | boyett_zhang_garny_holden_2001/rawfile/HEAD/boyett_zhang_garny_holden_2001.cellml |
| | | iribe_kohl_noble_2006/rawfile/HEAD/iribe_kohl_noble_2006.cellml |
| | | izakov_katsnelson_blyakhman_markhasin_shkylar_1991/rawfile/HEAD/izakov_katsnelson_blyakhman_markhasin_shkylar_1991.cellml |
| | | stern_song_sham_yang_boheler_rios_1999/rawfile/HEAD/stern_song_sham_yang_boheler_rios_1999.cellml |
| | | devries_sherman_2000/rawfile/HEAD/devries_sherman_2000.cellml |
| | | marhl_haberichter_brumen_heinrich_2000/rawfile/HEAD/marhl_haberichter_brumen_heinrich_2000.cellml |
| 39 | brain model | 546/rawfile/HEAD/cloutier_2009.cellml |
| 40 | socrates dokos, branko celler, and nigel lovell | baylor_hollingworth_chandler_2002/rawfile/HEAD/baylor_hollingworth_chandler_2002_b.cellml |
| | | baylor_hollingworth_chandler_2002/rawfile/HEAD/baylor_hollingworth_chandler_2002_d.cellml |
| | | luo_rudy_1994/rawfile/HEAD/luo_rudy_1994.cellml |
| | | colegrove_albrecht_friel_2000/rawfile/HEAD/colegrove_albrecht_friel_2000.cellml |
| | | dougherty_wright_yew_2005/rawfile/HEAD/dougherty_wright_yew_2005.cellml |
| | | yamaguchi_takaki_matsubara_yasuhara_suga_1996/rawfile/HEAD/yamaguchi_takaki_matsubara_yasuhara_suga_1996.cellml |
| | | boyett_zhang_garny_holden_2001/rawfile/HEAD/boyett_zhang_garny_holden_2001.cellml |
| | | iribe_kohl_noble_2006/rawfile/HEAD/iribe_kohl_noble_2006.cellml |
| | | izakov_katsnelson_blyakhman_markhasin_shkylar_1991/rawfile/HEAD/izakov_katsnelson_blyakhman_markhasin_shkylar_1991.cellml |
| | | stern_song_sham_yang_boheler_rios_1999/rawfile/HEAD/stern_song_sham_yang_boheler_rios_1999.cellml |
| | | devries_sherman_2000/rawfile/HEAD/devries_sherman_2000.cellml |
| | | marhl_haberichter_brumen_heinrich_2000/rawfile/HEAD/marhl_haberichter_brumen_heinrich_2000.cellml |
| 41 | rabbit | baylor_hollingworth_chandler_2002/rawfile/HEAD/baylor_hollingworth_chandler_2002_b.cellml |
| | | baylor_hollingworth_chandler_2002/rawfile/HEAD/baylor_hollingworth_chandler_2002_d.cellml |
| | | luo_rudy_1994/rawfile/HEAD/luo_rudy_1994.cellml |
| | | colegrove_albrecht_friel_2000/rawfile/HEAD/colegrove_albrecht_friel_2000.cellml |
| | | dougherty_wright_yew_2005/rawfile/HEAD/dougherty_wright_yew_2005.cellml |
| | | yamaguchi_takaki_matsubara_yasuhara_suga_1996/rawfile/HEAD/yamaguchi_takaki_matsubara_yasuhara_suga_1996.cellml |
| | | boyett_zhang_garny_holden_2001/rawfile/HEAD/boyett_zhang_garny_holden_2001.cellml |
| | | iribe_kohl_noble_2006/rawfile/HEAD/iribe_kohl_noble_2006.cellml |
| | | izakov_katsnelson_blyakhman_markhasin_shkylar_1991/rawfile/HEAD/izakov_katsnelson_blyakhman_markhasin_shkylar_1991.cellml |
| | | stern_song_sham_yang_boheler_rios_1999/rawfile/HEAD/stern_song_sham_yang_boheler_rios_1999.cellml |
| | | devries_sherman_2000/rawfile/HEAD/devries_sherman_2000.cellml |
| | | marhl_haberichter_brumen_heinrich_2000/rawfile/HEAD/marhl_haberichter_brumen_heinrich_2000.cellml |
| 42 | kidney | baylor_hollingworth_chandler_2002/rawfile/HEAD/baylor_hollingworth_chandler_2002_b.cellml |
| | | baylor_hollingworth_chandler_2002/rawfile/HEAD/baylor_hollingworth_chandler_2002_d.cellml |
| | | luo_rudy_1994/rawfile/HEAD/luo_rudy_1994.cellml |
| | | colegrove_albrecht_friel_2000/rawfile/HEAD/colegrove_albrecht_friel_2000.cellml |
| | | 267/rawfile/HEAD/mackenzie_1996-mouse-baso.cellml |
| | | 267/rawfile/HEAD/SEDML/mackenzie_1996/mackenzie_1996.cellml |
| | | dougherty_wright_yew_2005/rawfile/HEAD/dougherty_wright_yew_2005.cellml |
| | | yamaguchi_takaki_matsubara_yasuhara_suga_1996/rawfile/HEAD/yamaguchi_takaki_matsubara_yasuhara_suga_1996.cellml |
| | | 267/rawfile/HEAD/mackenzie_1996.cellml |
| | | boyett_zhang_garny_holden_2001/rawfile/HEAD/boyett_zhang_garny_holden_2001.cellml |
| | | iribe_kohl_noble_2006/rawfile/HEAD/iribe_kohl_noble_2006.cellml |
| | | izakov_katsnelson_blyakhman_markhasin_shkylar_1991/rawfile/HEAD/izakov_katsnelson_blyakhman_markhasin_shkylar_1991.cellml |
| | | stern_song_sham_yang_boheler_rios_1999/rawfile/HEAD/stern_song_sham_yang_boheler_rios_1999.cellml |
| | | devries_sherman_2000/rawfile/HEAD/devries_sherman_2000.cellml |
| | | marhl_haberichter_brumen_heinrich_2000/rawfile/HEAD/marhl_haberichter_brumen_heinrich_2000.cellml |
| 43 | protein | baylor_hollingworth_chandler_2002/rawfile/HEAD/baylor_hollingworth_chandler_2002_b.cellml |
| | | baylor_hollingworth_chandler_2002/rawfile/HEAD/baylor_hollingworth_chandler_2002_d.cellml |
| | | luo_rudy_1994/rawfile/HEAD/luo_rudy_1994.cellml |
| | | colegrove_albrecht_friel_2000/rawfile/HEAD/colegrove_albrecht_friel_2000.cellml |
| | | dougherty_wright_yew_2005/rawfile/HEAD/dougherty_wright_yew_2005.cellml |
| | | yamaguchi_takaki_matsubara_yasuhara_suga_1996/rawfile/HEAD/yamaguchi_takaki_matsubara_yasuhara_suga_1996.cellml |

| | | |
|---|---|---|
| | | boyett_zhang_garny_holden_2001/rawfile/HEAD/boyett_zhang_garny_holden_2001.cellml |
| | | iribe_kohl_noble_2006/rawfile/HEAD/iribe_kohl_noble_2006.cellml |
| | | izakov_katsnelson_blyakhman_markhasin_shkylar_1991/rawfile/HEAD/izakov_katsnelson_blyakhman_markhasin_shkylar_1991.cellml |
| | | stern_song_sham_yang_boheler_rios_1999/rawfile/HEAD/stern_song_sham_yang_boheler_rios_1999.cellml |
| | | devries_sherman_2000/rawfile/HEAD/devries_sherman_2000.cellml |
| | | marhl_haberichter_brumen_heinrich_2000/rawfile/HEAD/marhl_haberichter_brumen_heinrich_2000.cellml |
| 44 | sodium channel | baylor_hollingworth_chandler_2002/rawfile/HEAD/baylor_hollingworth_chandler_2002_b.cellml |
| | | baylor_hollingworth_chandler_2002/rawfile/HEAD/baylor_hollingworth_chandler_2002_d.cellml |
| | | luo_rudy_1994/rawfile/HEAD/luo_rudy_1994.cellml |
| | | colegrove_albrecht_friel_2000/rawfile/HEAD/colegrove_albrecht_friel_2000.cellml |
| | | dougherty_wright_yew_2005/rawfile/HEAD/dougherty_wright_yew_2005.cellml |
| | | yamaguchi_takaki_matsubara_yasuhara_suga_1996/rawfile/HEAD/yamaguchi_takaki_matsubara_yasuhara_suga_1996.cellml |
| | | boyett_zhang_garny_holden_2001/rawfile/HEAD/boyett_zhang_garny_holden_2001.cellml |
| | | iribe_kohl_noble_2006/rawfile/HEAD/iribe_kohl_noble_2006.cellml |
| | | izakov_katsnelson_blyakhman_markhasin_shkylar_1991/rawfile/HEAD/izakov_katsnelson_blyakhman_markhasin_shkylar_1991.cellml |
| | | stern_song_sham_yang_boheler_rios_1999/rawfile/HEAD/stern_song_sham_yang_boheler_rios_1999.cellml |
| | | devries_sherman_2000/rawfile/HEAD/devries_sherman_2000.cellml |
| | | marhl_haberichter_brumen_heinrich_2000/rawfile/HEAD/marhl_haberichter_brumen_heinrich_2000.cellml |
| 45 | mitochondrial modulation of intracellular | fall_keizer_2001/rawfile/HEAD/fall_keizer_2001.cellml |
| 46 | a simplified model for mitochondrial atp production | baylor_hollingworth_chandler_2002/rawfile/HEAD/baylor_hollingworth_chandler_2002_b.cellml |
| | | baylor_hollingworth_chandler_2002/rawfile/HEAD/baylor_hollingworth_chandler_2002_d.cellml |
| | | luo_rudy_1994/rawfile/HEAD/luo_rudy_1994.cellml |
| | | colegrove_albrecht_friel_2000/rawfile/HEAD/colegrove_albrecht_friel_2000.cellml |
| | | dougherty_wright_yew_2005/rawfile/HEAD/dougherty_wright_yew_2005.cellml |
| | | yamaguchi_takaki_matsubara_yasuhara_suga_1996/rawfile/HEAD/yamaguchi_takaki_matsubara_yasuhara_suga_1996.cellml |
| | | boyett_zhang_garny_holden_2001/rawfile/HEAD/boyett_zhang_garny_holden_2001.cellml |
| | | iribe_kohl_noble_2006/rawfile/HEAD/iribe_kohl_noble_2006.cellml |
| | | izakov_katsnelson_blyakhman_markhasin_shkylar_1991/rawfile/HEAD/izakov_katsnelson_blyakhman_markhasin_shkylar_1991.cellml |
| | | stern_song_sham_yang_boheler_rios_1999/rawfile/HEAD/stern_song_sham_yang_boheler_rios_1999.cellml |
| | | devries_sherman_2000/rawfile/HEAD/devries_sherman_2000.cellml |
| | | marhl_haberichter_brumen_heinrich_2000/rawfile/HEAD/marhl_haberichter_brumen_heinrich_2000.cellml |
| 47 | an integrated model of cardiac mitochondrial energy metabolism and calcium dynamics | cortassa_aon_marban_winslow_orourke_2003/rawfile/HEAD/cortassa_aon_marban_winslow_orourke_2003.cellml |
| 48 | complex calcium oscillations and the role of mitochondria and cytosolic proteins | baylor_hollingworth_chandler_2002/rawfile/HEAD/baylor_hollingworth_chandler_2002_b.cellml |
| | | baylor_hollingworth_chandler_2002/rawfile/HEAD/baylor_hollingworth_chandler_2002_d.cellml |
| | | luo_rudy_1994/rawfile/HEAD/luo_rudy_1994.cellml |
| | | colegrove_albrecht_friel_2000/rawfile/HEAD/colegrove_albrecht_friel_2000.cellml |
| | | dougherty_wright_yew_2005/rawfile/HEAD/dougherty_wright_yew_2005.cellml |
| | | yamaguchi_takaki_matsubara_yasuhara_suga_1996/rawfile/HEAD/yamaguchi_takaki_matsubara_yasuhara_suga_1996.cellml |
| | | boyett_zhang_garny_holden_2001/rawfile/HEAD/boyett_zhang_garny_holden_2001.cellml |
| | | iribe_kohl_noble_2006/rawfile/HEAD/iribe_kohl_noble_2006.cellml |
| | | izakov_katsnelson_blyakhman_markhasin_shkylar_1991/rawfile/HEAD/izakov_katsnelson_blyakhman_markhasin_shkylar_1991.cellml |
| | | stern_song_sham_yang_boheler_rios_1999/rawfile/HEAD/stern_song_sham_yang_boheler_rios_1999.cellml |
| | | devries_sherman_2000/rawfile/HEAD/devries_sherman_2000.cellml |
| | | marhl_haberichter_brumen_heinrich_2000/rawfile/HEAD/marhl_haberichter_brumen_heinrich_2000.cellml |
| 49 | a mitochondrial oscillator dependent on reactive oxygen species | cortassa_aon_marban_winslow_orourke_2003/rawfile/HEAD/cortassa_aon_marban_winslow_orourke_2003.cellml |
| 50 | computer modeling of mitochondrial tricarboxylic acid cycle, oxidative phosphorylation, metabolite transport, and electrophysiology | baylor_hollingworth_chandler_2002/rawfile/HEAD/baylor_hollingworth_chandler_2002_b.cellml |
| | | baylor_hollingworth_chandler_2002/rawfile/HEAD/baylor_hollingworth_chandler_2002_d.cellml |
| | | luo_rudy_1994/rawfile/HEAD/luo_rudy_1994.cellml |
| | | colegrove_albrecht_friel_2000/rawfile/HEAD/colegrove_albrecht_friel_2000.cellml |
| | | dougherty_wright_yew_2005/rawfile/HEAD/dougherty_wright_yew_2005.cellml |
| | | yamaguchi_takaki_matsubara_yasuhara_suga_1996/rawfile/HEAD/yamaguchi_takaki_matsubara_yasuhara_suga_1996.cellml |
| | | boyett_zhang_garny_holden_2001/rawfile/HEAD/boyett_zhang_garny_holden_2001.cellml |
| | | iribe_kohl_noble_2006/rawfile/HEAD/iribe_kohl_noble_2006.cellml |
| | | izakov_katsnelson_blyakhman_markhasin_shkylar_1991/rawfile/HEAD/izakov_katsnelson_blyakhman_markhasin_shkylar_1991.cellml |
| | | stern_song_sham_yang_boheler_rios_1999/rawfile/HEAD/stern_song_sham_yang_boheler_rios_1999.cellml |
| | | devries_sherman_2000/rawfile/HEAD/devries_sherman_2000.cellml |
| | | marhl_haberichter_brumen_heinrich_2000/rawfile/HEAD/marhl_haberichter_brumen_heinrich_2000.cellml |
| | | baylor_hollingworth_chandler_2002/rawfile/HEAD/baylor_hollingworth_chandler_2002_b.cellml |



51 mitochondria

| | | |
|---|---|---|
| | | baylor_hollingworth_chandler_2002/rawfile/HEAD/baylor_hollingworth_chandler_2002_d.cellml |
| | | luo_rudy_1994/rawfile/HEAD/luo_rudy_1994.cellml |
| | | colegrove_albrecht_friel_2000/rawfile/HEAD/colegrove_albrecht_friel_2000.cellml |
| | | cortassa_aon_marban_winslow_orourke_2003/rawfile/HEAD/cortassa_aon_marban_winslow_orourke_2003.cellml |
| | | dougherty_wright_yew_2005/rawfile/HEAD/dougherty_wright_yew_2005.cellml |
| | | yamaguchi_takaki_matsubara_yasuhara_suga_1996/rawfile/HEAD/yamaguchi_takaki_matsubara_yasuhara_suga_1996.cellml |
| | | boyett_zhang_garny_holden_2001/rawfile/HEAD/boyett_zhang_garny_holden_2001.cellml |
| | | fall_keizer_2001/rawfile/HEAD/fall_keizer_2001.cellml |
| | | iribe_kohl_noble_2006/rawfile/HEAD/iribe_kohl_noble_2006.cellml |
| | | 546/rawfile/HEAD/cloutier_2009.cellml |
| | | albrecht_colegrove_friel_2002/rawfile/HEAD/albrecht_colegrove_friel_2002.cellml |
| | | izakov_katsnelson_blyakhman_markhasin_shkylar_1991/rawfile/HEAD/izakov_katsnelson_blyakhman_markhasin_shkylar_1991.cellml |
| | | stern_song_sham_yang_boheler_rios_1999/rawfile/HEAD/stern_song_sham_yang_boheler_rios_1999.cellml |
| | | devries_sherman_2000/rawfile/HEAD/devries_sherman_2000.cellml |
| | | marhl_haberichter_brumen_heinrich_2000/rawfile/HEAD/marhl_haberichter_brumen_heinrich_2000.cellml |
| 52 | t cell receptors | baylor_hollingworth_chandler_2002/rawfile/HEAD/baylor_hollingworth_chandler_2002_b.cellml |
| | | baylor_hollingworth_chandler_2002/rawfile/HEAD/baylor_hollingworth_chandler_2002_d.cellml |
| | | luo_rudy_1994/rawfile/HEAD/luo_rudy_1994.cellml |
| | | colegrove_albrecht_friel_2000/rawfile/HEAD/colegrove_albrecht_friel_2000.cellml |
| | | dougherty_wright_yew_2005/rawfile/HEAD/dougherty_wright_yew_2005.cellml |
| | | yamaguchi_takaki_matsubara_yasuhara_suga_1996/rawfile/HEAD/yamaguchi_takaki_matsubara_yasuhara_suga_1996.cellml |
| | | boyett_zhang_garny_holden_2001/rawfile/HEAD/boyett_zhang_garny_holden_2001.cellml |
| | | iribe_kohl_noble_2006/rawfile/HEAD/iribe_kohl_noble_2006.cellml |
| | | izakov_katsnelson_blyakhman_markhasin_shkylar_1991/rawfile/HEAD/izakov_katsnelson_blyakhman_markhasin_shkylar_1991.cellml |
| | | stern_song_sham_yang_boheler_rios_1999/rawfile/HEAD/stern_song_sham_yang_boheler_rios_1999.cellml |
| | | devries_sherman_2000/rawfile/HEAD/devries_sherman_2000.cellml |
| | | marhl_haberichter_brumen_heinrich_2000/rawfile/HEAD/marhl_haberichter_brumen_heinrich_2000.cellml |
| 53 | respiratory | baylor_hollingworth_chandler_2002/rawfile/HEAD/baylor_hollingworth_chandler_2002_b.cellml |
| | | baylor_hollingworth_chandler_2002/rawfile/HEAD/baylor_hollingworth_chandler_2002_d.cellml |
| | | luo_rudy_1994/rawfile/HEAD/luo_rudy_1994.cellml |
| | | colegrove_albrecht_friel_2000/rawfile/HEAD/colegrove_albrecht_friel_2000.cellml |
| | | dougherty_wright_yew_2005/rawfile/HEAD/dougherty_wright_yew_2005.cellml |
| | | yamaguchi_takaki_matsubara_yasuhara_suga_1996/rawfile/HEAD/yamaguchi_takaki_matsubara_yasuhara_suga_1996.cellml |
| | | boyett_zhang_garny_holden_2001/rawfile/HEAD/boyett_zhang_garny_holden_2001.cellml |
| | | iribe_kohl_noble_2006/rawfile/HEAD/iribe_kohl_noble_2006.cellml |
| | | izakov_katsnelson_blyakhman_markhasin_shkylar_1991/rawfile/HEAD/izakov_katsnelson_blyakhman_markhasin_shkylar_1991.cellml |
| | | stern_song_sham_yang_boheler_rios_1999/rawfile/HEAD/stern_song_sham_yang_boheler_rios_1999.cellml |
| | | devries_sherman_2000/rawfile/HEAD/devries_sherman_2000.cellml |
| | | marhl_haberichter_brumen_heinrich_2000/rawfile/HEAD/marhl_haberichter_brumen_heinrich_2000.cellml |
| 54 | atrial ap | baylor_hollingworth_chandler_2002/rawfile/HEAD/baylor_hollingworth_chandler_2002_b.cellml |
| | | baylor_hollingworth_chandler_2002/rawfile/HEAD/baylor_hollingworth_chandler_2002_d.cellml |
| | | luo_rudy_1994/rawfile/HEAD/luo_rudy_1994.cellml |
| | | colegrove_albrecht_friel_2000/rawfile/HEAD/colegrove_albrecht_friel_2000.cellml |
| | | dougherty_wright_yew_2005/rawfile/HEAD/dougherty_wright_yew_2005.cellml |
| | | yamaguchi_takaki_matsubara_yasuhara_suga_1996/rawfile/HEAD/yamaguchi_takaki_matsubara_yasuhara_suga_1996.cellml |
| | | boyett_zhang_garny_holden_2001/rawfile/HEAD/boyett_zhang_garny_holden_2001.cellml |
| | | iribe_kohl_noble_2006/rawfile/HEAD/iribe_kohl_noble_2006.cellml |
| | | izakov_katsnelson_blyakhman_markhasin_shkylar_1991/rawfile/HEAD/izakov_katsnelson_blyakhman_markhasin_shkylar_1991.cellml |
| | | stern_song_sham_yang_boheler_rios_1999/rawfile/HEAD/stern_song_sham_yang_boheler_rios_1999.cellml |
| | | devries_sherman_2000/rawfile/HEAD/devries_sherman_2000.cellml |
| | | marhl_haberichter_brumen_heinrich_2000/rawfile/HEAD/marhl_haberichter_brumen_heinrich_2000.cellml |
| 55 | calcium dynamics | baylor_hollingworth_chandler_2002/rawfile/HEAD/baylor_hollingworth_chandler_2002_b.cellml |
| | | baylor_hollingworth_chandler_2002/rawfile/HEAD/baylor_hollingworth_chandler_2002_d.cellml |
| | | luo_rudy_1994/rawfile/HEAD/luo_rudy_1994.cellml |
| | | colegrove_albrecht_friel_2000/rawfile/HEAD/colegrove_albrecht_friel_2000.cellml |
| | | cortassa_aon_marban_winslow_orourke_2003/rawfile/HEAD/cortassa_aon_marban_winslow_orourke_2003.cellml |
| | | dougherty_wright_yew_2005/rawfile/HEAD/dougherty_wright_yew_2005.cellml |
| | | yamaguchi_takaki_matsubara_yasuhara_suga_1996/rawfile/HEAD/yamaguchi_takaki_matsubara_yasuhara_suga_1996.cellml |
| | | noble_noble_2001/rawfile/HEAD/noble_noble_2001.cellml |
| | | boyett_zhang_garny_holden_2001/rawfile/HEAD/boyett_zhang_garny_holden_2001.cellml |

| | | |
|---|---|---|
| | | iribe_kohl_noble_2006/rawfile/HEAD/iribe_kohl_noble_2006.cellml |
| | | shannon_wang_puglisi_weber_bers_2004/rawfile/HEAD/shannon_wang_puglisi_weber_bers_2004_a.cellml |
| | | izakov_katsnelson_blyakhman_markhasin_shkylar_1991/rawfile/HEAD/izakov_katsnelson_blyakhman_markhasin_shkylar_1991.cellml |
| | | stern_song_sham_yang_boheler_rios_1999/rawfile/HEAD/stern_song_sham_yang_boheler_rios_1999.cellml |
| | | shiferaw_watanabe_garfinkel_weiss_karma_2003/rawfile/HEAD/shiferaw_watanabe_garfinkel_weiss_karma_2003.cellml |
| | | shannon_wang_puglisi_weber_bers_2004/rawfile/HEAD/shannon_wang_puglisi_weber_bers_2004_b.cellml |
| | | devries_sherman_2000/rawfile/HEAD/devries_sherman_2000.cellml |
| | | marhl_haberichter_brumen_heinrich_2000/rawfile/HEAD/marhl_haberichter_brumen_heinrich_2000.cellml |
| 56 | luo rudy phase -ii model | baylor_hollingworth_chandler_2002/rawfile/HEAD/baylor_hollingworth_chandler_2002_b.cellml |
| | | baylor_hollingworth_chandler_2002/rawfile/HEAD/baylor_hollingworth_chandler_2002_d.cellml |
| | | luo_rudy_1994/rawfile/HEAD/luo_rudy_1994.cellml |
| | | colegrove_albrecht_friel_2000/rawfile/HEAD/colegrove_albrecht_friel_2000.cellml |
| | | dougherty_wright_yew_2005/rawfile/HEAD/dougherty_wright_yew_2005.cellml |
| | | yamaguchi_takaki_matsubara_yasuhara_suga_1996/rawfile/HEAD/yamaguchi_takaki_matsubara_yasuhara_suga_1996.cellml |
| | | boyett_zhang_garny_holden_2001/rawfile/HEAD/boyett_zhang_garny_holden_2001.cellml |
| | | iribe_kohl_noble_2006/rawfile/HEAD/iribe_kohl_noble_2006.cellml |
| | | izakov_katsnelson_blyakhman_markhasin_shkylar_1991/rawfile/HEAD/izakov_katsnelson_blyakhman_markhasin_shkylar_1991.cellml |
| | | stern_song_sham_yang_boheler_rios_1999/rawfile/HEAD/stern_song_sham_yang_boheler_rios_1999.cellml |
| | | devries_sherman_2000/rawfile/HEAD/devries_sherman_2000.cellml |
| | | marhl_haberichter_brumen_heinrich_2000/rawfile/HEAD/marhl_haberichter_brumen_heinrich_2000.cellml |
| 57 | magnus, keizer | fall_keizer_2001/rawfile/HEAD/fall_keizer_2001.cellml |
| 58 | purkinje cell model | baylor_hollingworth_chandler_2002/rawfile/HEAD/baylor_hollingworth_chandler_2002_b.cellml |
| | | baylor_hollingworth_chandler_2002/rawfile/HEAD/baylor_hollingworth_chandler_2002_d.cellml |
| | | luo_rudy_1994/rawfile/HEAD/luo_rudy_1994.cellml |
| | | colegrove_albrecht_friel_2000/rawfile/HEAD/colegrove_albrecht_friel_2000.cellml |
| | | dougherty_wright_yew_2005/rawfile/HEAD/dougherty_wright_yew_2005.cellml |
| | | yamaguchi_takaki_matsubara_yasuhara_suga_1996/rawfile/HEAD/yamaguchi_takaki_matsubara_yasuhara_suga_1996.cellml |
| | | boyett_zhang_garny_holden_2001/rawfile/HEAD/boyett_zhang_garny_holden_2001.cellml |
| | | iribe_kohl_noble_2006/rawfile/HEAD/iribe_kohl_noble_2006.cellml |
| | | izakov_katsnelson_blyakhman_markhasin_shkylar_1991/rawfile/HEAD/izakov_katsnelson_blyakhman_markhasin_shkylar_1991.cellml |
| | | stern_song_sham_yang_boheler_rios_1999/rawfile/HEAD/stern_song_sham_yang_boheler_rios_1999.cellml |
| | | devries_sherman_2000/rawfile/HEAD/devries_sherman_2000.cellml |
| | | marhl_haberichter_brumen_heinrich_2000/rawfile/HEAD/marhl_haberichter_brumen_heinrich_2000.cellml |
| 59 | human atrial cell model | baylor_hollingworth_chandler_2002/rawfile/HEAD/baylor_hollingworth_chandler_2002_b.cellml |
| | | baylor_hollingworth_chandler_2002/rawfile/HEAD/baylor_hollingworth_chandler_2002_d.cellml |
| | | luo_rudy_1994/rawfile/HEAD/luo_rudy_1994.cellml |
| | | colegrove_albrecht_friel_2000/rawfile/HEAD/colegrove_albrecht_friel_2000.cellml |
| | | dougherty_wright_yew_2005/rawfile/HEAD/dougherty_wright_yew_2005.cellml |
| | | yamaguchi_takaki_matsubara_yasuhara_suga_1996/rawfile/HEAD/yamaguchi_takaki_matsubara_yasuhara_suga_1996.cellml |
| | | boyett_zhang_garny_holden_2001/rawfile/HEAD/boyett_zhang_garny_holden_2001.cellml |
| | | iribe_kohl_noble_2006/rawfile/HEAD/iribe_kohl_noble_2006.cellml |
| | | izakov_katsnelson_blyakhman_markhasin_shkylar_1991/rawfile/HEAD/izakov_katsnelson_blyakhman_markhasin_shkylar_1991.cellml |
| | | stern_song_sham_yang_boheler_rios_1999/rawfile/HEAD/stern_song_sham_yang_boheler_rios_1999.cellml |
| | | devries_sherman_2000/rawfile/HEAD/devries_sherman_2000.cellml |
| | | marhl_haberichter_brumen_heinrich_2000/rawfile/HEAD/marhl_haberichter_brumen_heinrich_2000.cellml |
| 60 | purkinje fibre human | baylor_hollingworth_chandler_2002/rawfile/HEAD/baylor_hollingworth_chandler_2002_b.cellml |
| | | baylor_hollingworth_chandler_2002/rawfile/HEAD/baylor_hollingworth_chandler_2002_d.cellml |
| | | luo_rudy_1994/rawfile/HEAD/luo_rudy_1994.cellml |
| | | colegrove_albrecht_friel_2000/rawfile/HEAD/colegrove_albrecht_friel_2000.cellml |
| | | dougherty_wright_yew_2005/rawfile/HEAD/dougherty_wright_yew_2005.cellml |
| | | yamaguchi_takaki_matsubara_yasuhara_suga_1996/rawfile/HEAD/yamaguchi_takaki_matsubara_yasuhara_suga_1996.cellml |
| | | boyett_zhang_garny_holden_2001/rawfile/HEAD/boyett_zhang_garny_holden_2001.cellml |
| | | iribe_kohl_noble_2006/rawfile/HEAD/iribe_kohl_noble_2006.cellml |
| | | izakov_katsnelson_blyakhman_markhasin_shkylar_1991/rawfile/HEAD/izakov_katsnelson_blyakhman_markhasin_shkylar_1991.cellml |
| | | stern_song_sham_yang_boheler_rios_1999/rawfile/HEAD/stern_song_sham_yang_boheler_rios_1999.cellml |
| | | devries_sherman_2000/rawfile/HEAD/devries_sherman_2000.cellml |
| | | marhl_haberichter_brumen_heinrich_2000/rawfile/HEAD/marhl_haberichter_brumen_heinrich_2000.cellml |
| 61 | human purkinje | baylor_hollingworth_chandler_2002/rawfile/HEAD/baylor_hollingworth_chandler_2002_b.cellml |
| | | baylor_hollingworth_chandler_2002/rawfile/HEAD/baylor_hollingworth_chandler_2002_d.cellml |
| | | luo_rudy_1994/rawfile/HEAD/luo_rudy_1994.cellml |

| | | |
|---|---|---|
| | | colegrove_albrecht_friel_2000/rawfile/HEAD/colegrove_albrecht_friel_2000.cellml |
| | | dougherty_wright_yew_2005/rawfile/HEAD/dougherty_wright_yew_2005.cellml |
| | | yamaguchi_takaki_matsubara_yasuhara_suga_1996/rawfile/HEAD/yamaguchi_takaki_matsubara_yasuhara_suga_1996.cellml |
| | | boyett_zhang_garny_holden_2001/rawfile/HEAD/boyett_zhang_garny_holden_2001.cellml |
| | | iribe_kohl_noble_2006/rawfile/HEAD/iribe_kohl_noble_2006.cellml |
| | | izakov_katsnelson_blyakhman_markhasin_shkylar_1991/rawfile/HEAD/izakov_katsnelson_blyakhman_markhasin_shkylar_1991.cellml |
| | | stern_song_sham_yang_boheler_rios_1999/rawfile/HEAD/stern_song_sham_yang_boheler_rios_1999.cellml |
| | | devries_sherman_2000/rawfile/HEAD/devries_sherman_2000.cellml |
| | | marhl_haberichter_brumen_heinrich_2000/rawfile/HEAD/marhl_haberichter_brumen_heinrich_2000.cellml |
| 62 | matlab calcium | faber_rudy_2000/rawfile/HEAD/faber_rudy_modified_version_2000.cellml |
| 63 | an integrative dynamic model of brain energy metabolism using in vivo neurochemical measurements | 546/rawfile/HEAD/cloutier_2009.cellml |
| 64 | ryanodine | baylor_hollingworth_chandler_2002/rawfile/HEAD/baylor_hollingworth_chandler_2002_b.cellml |
| | | baylor_hollingworth_chandler_2002/rawfile/HEAD/baylor_hollingworth_chandler_2002_d.cellml |
| | | luo_rudy_1994/rawfile/HEAD/luo_rudy_1994.cellml |
| | | michailova_mcculloch_2001/rawfile/HEAD/michailova_mcculloch_2001.cellml |
| | | colegrove_albrecht_friel_2000/rawfile/HEAD/colegrove_albrecht_friel_2000.cellml |
| | | dougherty_wright_yew_2005/rawfile/HEAD/dougherty_wright_yew_2005.cellml |
| | | jafri_rice_winslow_1998/rawfile/HEAD/jafri_rice_winslow_1998_a.cellml |
| | | jafri_rice_winslow_1998/rawfile/HEAD/jafri_rice_winslow_1998_b.cellml |
| | | yamaguchi_takaki_matsubara_yasuhara_suga_1996/rawfile/HEAD/yamaguchi_takaki_matsubara_yasuhara_suga_1996.cellml |
| | | boyett_zhang_garny_holden_2001/rawfile/HEAD/boyett_zhang_garny_holden_2001.cellml |
| | | iribe_kohl_noble_2006/rawfile/HEAD/iribe_kohl_noble_2006.cellml |
| | | winslow_rice_jafri_marban_ororke_1999/rawfile/HEAD/winslow_rice_jafri_marban_ororke_1999.cellml |
| | | izakov_katsnelson_blyakhman_markhasin_shkylar_1991/rawfile/HEAD/izakov_katsnelson_blyakhman_markhasin_shkylar_1991.cellml |
| | | stern_song_sham_yang_boheler_rios_1999/rawfile/HEAD/stern_song_sham_yang_boheler_rios_1999.cellml |
| | | devries_sherman_2000/rawfile/HEAD/devries_sherman_2000.cellml |
| | | marhl_haberichter_brumen_heinrich_2000/rawfile/HEAD/marhl_haberichter_brumen_heinrich_2000.cellml |
| 65 | magnus, keizer, 1998 | magnus_keizer_1998/rawfile/HEAD/magnus_keizer_1998.cellml |
| | | fall_keizer_2001/rawfile/HEAD/fall_keizer_2001.cellml |
| 66 | brain | 546/rawfile/HEAD/cloutier_2009.cellml |
| 67 | noble'98 | noble_noble_2001/rawfile/HEAD/noble_noble_2001.cellml |
| 68 | nuclear erythyroid 2 | baylor_hollingworth_chandler_2002/rawfile/HEAD/baylor_hollingworth_chandler_2002_b.cellml |
| | | baylor_hollingworth_chandler_2002/rawfile/HEAD/baylor_hollingworth_chandler_2002_d.cellml |
| | | luo_rudy_1994/rawfile/HEAD/luo_rudy_1994.cellml |
| | | colegrove_albrecht_friel_2000/rawfile/HEAD/colegrove_albrecht_friel_2000.cellml |
| | | dougherty_wright_yew_2005/rawfile/HEAD/dougherty_wright_yew_2005.cellml |
| | | yamaguchi_takaki_matsubara_yasuhara_suga_1996/rawfile/HEAD/yamaguchi_takaki_matsubara_yasuhara_suga_1996.cellml |
| | | boyett_zhang_garny_holden_2001/rawfile/HEAD/boyett_zhang_garny_holden_2001.cellml |
| | | iribe_kohl_noble_2006/rawfile/HEAD/iribe_kohl_noble_2006.cellml |
| | | izakov_katsnelson_blyakhman_markhasin_shkylar_1991/rawfile/HEAD/izakov_katsnelson_blyakhman_markhasin_shkylar_1991.cellml |
| | | stern_song_sham_yang_boheler_rios_1999/rawfile/HEAD/stern_song_sham_yang_boheler_rios_1999.cellml |
| | | devries_sherman_2000/rawfile/HEAD/devries_sherman_2000.cellml |
| | | marhl_haberichter_brumen_heinrich_2000/rawfile/HEAD/marhl_haberichter_brumen_heinrich_2000.cellml |
| 69 | cytosolic inhibitor | baylor_hollingworth_chandler_2002/rawfile/HEAD/baylor_hollingworth_chandler_2002_b.cellml |
| | | baylor_hollingworth_chandler_2002/rawfile/HEAD/baylor_hollingworth_chandler_2002_d.cellml |
| | | 267/rawfile/HEAD/eskandari_2005.cellml |
| | | luo_rudy_1994/rawfile/HEAD/luo_rudy_1994.cellml |
| | | colegrove_albrecht_friel_2000/rawfile/HEAD/colegrove_albrecht_friel_2000.cellml |
| | | dougherty_wright_yew_2005/rawfile/HEAD/dougherty_wright_yew_2005.cellml |
| | | yamaguchi_takaki_matsubara_yasuhara_suga_1996/rawfile/HEAD/yamaguchi_takaki_matsubara_yasuhara_suga_1996.cellml |
| | | boyett_zhang_garny_holden_2001/rawfile/HEAD/boyett_zhang_garny_holden_2001.cellml |
| | | 267/rawfile/HEAD/SEDML/eskandari_2005/eskandari_2005.cellml |
| | | iribe_kohl_noble_2006/rawfile/HEAD/iribe_kohl_noble_2006.cellml |
| | | izakov_katsnelson_blyakhman_markhasin_shkylar_1991/rawfile/HEAD/izakov_katsnelson_blyakhman_markhasin_shkylar_1991.cellml |
| | | stern_song_sham_yang_boheler_rios_1999/rawfile/HEAD/stern_song_sham_yang_boheler_rios_1999.cellml |
| | | devries_sherman_2000/rawfile/HEAD/devries_sherman_2000.cellml |
| | | marhl_haberichter_brumen_heinrich_2000/rawfile/HEAD/marhl_haberichter_brumen_heinrich_2000.cellml |
| 70 | lumped-parameter cardiovascular | baylor_hollingworth_chandler_2002/rawfile/HEAD/baylor_hollingworth_chandler_2002_b.cellml |

| | | |
|---|---|---|
| | | baylor_hollingworth_chandler_2002/rawfile/HEAD/baylor_hollingworth_chandler_2002_d.cellml |
| | | luo_rudy_1994/rawfile/HEAD/luo_rudy_1994.cellml |
| | | colegrove_albrecht_friel_2000/rawfile/HEAD/colegrove_albrecht_friel_2000.cellml |
| | | dougherty_wright_yew_2005/rawfile/HEAD/dougherty_wright_yew_2005.cellml |
| | | yamaguchi_takaki_matsubara_yasuhara_suga_1996/rawfile/HEAD/yamaguchi_takaki_matsubara_yasuhara_suga_1996.cellml |
| | | boyett_zhang_garny_holden_2001/rawfile/HEAD/boyett_zhang_garny_holden_2001.cellml |
| | | iribe_kohl_noble_2006/rawfile/HEAD/iribe_kohl_noble_2006.cellml |
| | | izakov_katsnelson_blyakhman_markhasin_shkylar_1991/rawfile/HEAD/izakov_katsnelson_blyakhman_markhasin_shkylar_1991.cellml |
| | | stern_song_sham_yang_boheler_rios_1999/rawfile/HEAD/stern_song_sham_yang_boheler_rios_1999.cellml |
| | | devries_sherman_2000/rawfile/HEAD/devries_sherman_2000.cellml |
| | | marhl_haberichter_brumen_heinrich_2000/rawfile/HEAD/marhl_haberichter_brumen_heinrich_2000.cellml |
| 71 | mackenzie, loo, panayotova-heiermann and wright | 267/rawfile/HEAD/mackenzie_1996-mouse-baso.cellml |
| | | 267/rawfile/HEAD/SEDML/mackenzie_1996/mackenzie_1996.cellml |
| | | 267/rawfile/HEAD/mackenzie_1996.cellml |
| 72 | calcium cerebral palsy | bertram_satin_zhang_smolen_sherman_2004/rawfile/HEAD/bertram_satin_zhang_smolen_sherman_2004_b.cellml |
| | | baylor_hollingworth_chandler_2002/rawfile/HEAD/baylor_hollingworth_chandler_2002_b.cellml |
| | | baylor_hollingworth_chandler_2002/rawfile/HEAD/baylor_hollingworth_chandler_2002_d.cellml |
| | | luo_rudy_1994/rawfile/HEAD/luo_rudy_1994.cellml |
| | | colegrove_albrecht_friel_2000/rawfile/HEAD/colegrove_albrecht_friel_2000.cellml |
| | | dougherty_wright_yew_2005/rawfile/HEAD/dougherty_wright_yew_2005.cellml |
| | | yamaguchi_takaki_matsubara_yasuhara_suga_1996/rawfile/HEAD/yamaguchi_takaki_matsubara_yasuhara_suga_1996.cellml |
| | | boyett_zhang_garny_holden_2001/rawfile/HEAD/boyett_zhang_garny_holden_2001.cellml |
| | | 267/rawfile/HEAD/bindschadler_sneyd_2001.cellml |
| | | iribe_kohl_noble_2006/rawfile/HEAD/iribe_kohl_noble_2006.cellml |
| | | magnus_keizer_1998/rawfile/HEAD/magnus_keizer_1998.cellml |
| | | bertram_satin_zhang_smolen_sherman_2004/rawfile/HEAD/bertram_satin_zhang_smolen_sherman_2004_a.cellml |
| | | izakov_katsnelson_blyakhman_markhasin_shkylar_1991/rawfile/HEAD/izakov_katsnelson_blyakhman_markhasin_shkylar_1991.cellml |
| | | stern_song_sham_yang_boheler_rios_1999/rawfile/HEAD/stern_song_sham_yang_boheler_rios_1999.cellml |
| | | bindschadler_sneyd_2001/rawfile/HEAD/bindschadler_sneyd_2001.cellml |
| | | devries_sherman_2000/rawfile/HEAD/devries_sherman_2000.cellml |
| | | marhl_haberichter_brumen_heinrich_2000/rawfile/HEAD/marhl_haberichter_brumen_heinrich_2000.cellml |
| 73 | ventricular | baylor_hollingworth_chandler_2002/rawfile/HEAD/baylor_hollingworth_chandler_2002_b.cellml |
| | | baylor_hollingworth_chandler_2002/rawfile/HEAD/baylor_hollingworth_chandler_2002_d.cellml |
| | | luo_rudy_1994/rawfile/HEAD/luo_rudy_1994.cellml |
| | | colegrove_albrecht_friel_2000/rawfile/HEAD/colegrove_albrecht_friel_2000.cellml |
| | | dougherty_wright_yew_2005/rawfile/HEAD/dougherty_wright_yew_2005.cellml |
| | | yamaguchi_takaki_matsubara_yasuhara_suga_1996/rawfile/HEAD/yamaguchi_takaki_matsubara_yasuhara_suga_1996.cellml |
| | | boyett_zhang_garny_holden_2001/rawfile/HEAD/boyett_zhang_garny_holden_2001.cellml |
| | | iribe_kohl_noble_2006/rawfile/HEAD/iribe_kohl_noble_2006.cellml |
| | | izakov_katsnelson_blyakhman_markhasin_shkylar_1991/rawfile/HEAD/izakov_katsnelson_blyakhman_markhasin_shkylar_1991.cellml |
| | | stern_song_sham_yang_boheler_rios_1999/rawfile/HEAD/stern_song_sham_yang_boheler_rios_1999.cellml |
| | | shiferaw_watanabe_garfinkel_weiss_karma_2003/rawfile/HEAD/shiferaw_watanabe_garfinkel_weiss_karma_2003.cellml |
| | | devries_sherman_2000/rawfile/HEAD/devries_sherman_2000.cellml |
| | | marhl_haberichter_brumen_heinrich_2000/rawfile/HEAD/marhl_haberichter_brumen_heinrich_2000.cellml |
| 74 | calcium heart | michailova_mcculloch_2001/rawfile/HEAD/michailova_mcculloch_2001.cellml |
| | | jafri_rice_winslow_1998/rawfile/HEAD/jafri_rice_winslow_1998_a.cellml |
| | | winslow_rice_jafri_marban_ororke_1999/rawfile/HEAD/winslow_rice_jafri_marban_ororke_1999.cellml |
| | | jafri_rice_winslow_1998/rawfile/HEAD/jafri_rice_winslow_1998_b.cellml |
| 75 | noble ventricle 2000 | tentusscher_noble_noble_panfilov_2004/rawfile/HEAD/tentusscher_noble_noble_panfilov_2004_c.cellml |
| | | tentusscher_noble_noble_panfilov_2004/rawfile/HEAD/tentusscher_noble_noble_panfilov_2004_b.cellml |
| | | tentusscher_noble_noble_panfilov_2004/rawfile/HEAD/tentusscher_noble_noble_panfilov_2004_a.cellml |
| 76 | mapk human | baylor_hollingworth_chandler_2002/rawfile/HEAD/baylor_hollingworth_chandler_2002_b.cellml |
| | | baylor_hollingworth_chandler_2002/rawfile/HEAD/baylor_hollingworth_chandler_2002_d.cellml |
| | | luo_rudy_1994/rawfile/HEAD/luo_rudy_1994.cellml |
| | | colegrove_albrecht_friel_2000/rawfile/HEAD/colegrove_albrecht_friel_2000.cellml |
| | | dougherty_wright_yew_2005/rawfile/HEAD/dougherty_wright_yew_2005.cellml |
| | | yamaguchi_takaki_matsubara_yasuhara_suga_1996/rawfile/HEAD/yamaguchi_takaki_matsubara_yasuhara_suga_1996.cellml |
| | | boyett_zhang_garny_holden_2001/rawfile/HEAD/boyett_zhang_garny_holden_2001.cellml |
| | | iribe_kohl_noble_2006/rawfile/HEAD/iribe_kohl_noble_2006.cellml |

| | | |
|---|---|---|
| | | izakov_katsnelson_blyakhman_markhasin_shkylar_1991/rawfile/HEAD/izakov_katsnelson_blyakhman_markhasin_shkylar_1991.cellml |
| | | stern_song_sham_yang_boheler_rios_1999/rawfile/HEAD/stern_song_sham_yang_boheler_rios_1999.cellml |
| | | devries_sherman_2000/rawfile/HEAD/devries_sherman_2000.cellml |
| | | marhl_haberichter_brumen_heinrich_2000/rawfile/HEAD/marhl_haberichter_brumen_heinrich_2000.cellml |
| 77 | mapk | baylor_hollingworth_chandler_2002/rawfile/HEAD/baylor_hollingworth_chandler_2002_b.cellml |
| | | baylor_hollingworth_chandler_2002/rawfile/HEAD/baylor_hollingworth_chandler_2002_d.cellml |
| | | luo_rudy_1994/rawfile/HEAD/luo_rudy_1994.cellml |
| | | colegrove_albrecht_friel_2000/rawfile/HEAD/colegrove_albrecht_friel_2000.cellml |
| | | dougherty_wright_yew_2005/rawfile/HEAD/dougherty_wright_yew_2005.cellml |
| | | yamaguchi_takaki_matsubara_yasuhara_suga_1996/rawfile/HEAD/yamaguchi_takaki_matsubara_yasuhara_suga_1996.cellml |
| | | boyett_zhang_garny_holden_2001/rawfile/HEAD/boyett_zhang_garny_holden_2001.cellml |
| | | iribe_kohl_noble_2006/rawfile/HEAD/iribe_kohl_noble_2006.cellml |
| | | izakov_katsnelson_blyakhman_markhasin_shkylar_1991/rawfile/HEAD/izakov_katsnelson_blyakhman_markhasin_shkylar_1991.cellml |
| | | stern_song_sham_yang_boheler_rios_1999/rawfile/HEAD/stern_song_sham_yang_boheler_rios_1999.cellml |
| | | devries_sherman_2000/rawfile/HEAD/devries_sherman_2000.cellml |
| | | marhl_haberichter_brumen_heinrich_2000/rawfile/HEAD/marhl_haberichter_brumen_heinrich_2000.cellml |
| 78 | electrolytes disturbence with complete heart block | baylor_hollingworth_chandler_2002/rawfile/HEAD/baylor_hollingworth_chandler_2002_b.cellml |
| | | baylor_hollingworth_chandler_2002/rawfile/HEAD/baylor_hollingworth_chandler_2002_d.cellml |
| | | luo_rudy_1994/rawfile/HEAD/luo_rudy_1994.cellml |
| | | colegrove_albrecht_friel_2000/rawfile/HEAD/colegrove_albrecht_friel_2000.cellml |
| | | dougherty_wright_yew_2005/rawfile/HEAD/dougherty_wright_yew_2005.cellml |
| | | yamaguchi_takaki_matsubara_yasuhara_suga_1996/rawfile/HEAD/yamaguchi_takaki_matsubara_yasuhara_suga_1996.cellml |
| | | boyett_zhang_garny_holden_2001/rawfile/HEAD/boyett_zhang_garny_holden_2001.cellml |
| | | iribe_kohl_noble_2006/rawfile/HEAD/iribe_kohl_noble_2006.cellml |
| | | izakov_katsnelson_blyakhman_markhasin_shkylar_1991/rawfile/HEAD/izakov_katsnelson_blyakhman_markhasin_shkylar_1991.cellml |
| | | stern_song_sham_yang_boheler_rios_1999/rawfile/HEAD/stern_song_sham_yang_boheler_rios_1999.cellml |
| | | devries_sherman_2000/rawfile/HEAD/devries_sherman_2000.cellml |
| | | marhl_haberichter_brumen_heinrich_2000/rawfile/HEAD/marhl_haberichter_brumen_heinrich_2000.cellml |
| 79 | mitochondrial | fall_keizer_2001/rawfile/HEAD/fall_keizer_2001.cellml |
| 80 | atp | baylor_hollingworth_chandler_2002/rawfile/HEAD/baylor_hollingworth_chandler_2002_b.cellml |
| | | baylor_hollingworth_chandler_2002/rawfile/HEAD/baylor_hollingworth_chandler_2002_d.cellml |
| | | luo_rudy_1994/rawfile/HEAD/luo_rudy_1994.cellml |
| | | colegrove_albrecht_friel_2000/rawfile/HEAD/colegrove_albrecht_friel_2000.cellml |
| | | dougherty_wright_yew_2005/rawfile/HEAD/dougherty_wright_yew_2005.cellml |
| | | yamaguchi_takaki_matsubara_yasuhara_suga_1996/rawfile/HEAD/yamaguchi_takaki_matsubara_yasuhara_suga_1996.cellml |
| | | boyett_zhang_garny_holden_2001/rawfile/HEAD/boyett_zhang_garny_holden_2001.cellml |
| | | iribe_kohl_noble_2006/rawfile/HEAD/iribe_kohl_noble_2006.cellml |
| | | izakov_katsnelson_blyakhman_markhasin_shkylar_1991/rawfile/HEAD/izakov_katsnelson_blyakhman_markhasin_shkylar_1991.cellml |
| | | stern_song_sham_yang_boheler_rios_1999/rawfile/HEAD/stern_song_sham_yang_boheler_rios_1999.cellml |
| | | devries_sherman_2000/rawfile/HEAD/devries_sherman_2000.cellml |
| | | marhl_haberichter_brumen_heinrich_2000/rawfile/HEAD/marhl_haberichter_brumen_heinrich_2000.cellml |
| 81 | luo-rudy model | difrancesco_noble_1985/rawfile/HEAD/difrancesco_noble_1985.cellml |
| 82 | cancer metabolism | baylor_hollingworth_chandler_2002/rawfile/HEAD/baylor_hollingworth_chandler_2002_b.cellml |
| | | baylor_hollingworth_chandler_2002/rawfile/HEAD/baylor_hollingworth_chandler_2002_d.cellml |
| | | luo_rudy_1994/rawfile/HEAD/luo_rudy_1994.cellml |
| | | colegrove_albrecht_friel_2000/rawfile/HEAD/colegrove_albrecht_friel_2000.cellml |
| | | dougherty_wright_yew_2005/rawfile/HEAD/dougherty_wright_yew_2005.cellml |
| | | yamaguchi_takaki_matsubara_yasuhara_suga_1996/rawfile/HEAD/yamaguchi_takaki_matsubara_yasuhara_suga_1996.cellml |
| | | boyett_zhang_garny_holden_2001/rawfile/HEAD/boyett_zhang_garny_holden_2001.cellml |
| | | iribe_kohl_noble_2006/rawfile/HEAD/iribe_kohl_noble_2006.cellml |
| | | izakov_katsnelson_blyakhman_markhasin_shkylar_1991/rawfile/HEAD/izakov_katsnelson_blyakhman_markhasin_shkylar_1991.cellml |
| | | stern_song_sham_yang_boheler_rios_1999/rawfile/HEAD/stern_song_sham_yang_boheler_rios_1999.cellml |
| | | devries_sherman_2000/rawfile/HEAD/devries_sherman_2000.cellml |
| | | marhl_haberichter_brumen_heinrich_2000/rawfile/HEAD/marhl_haberichter_brumen_heinrich_2000.cellml |
| 83 | protein regulation | baylor_hollingworth_chandler_2002/rawfile/HEAD/baylor_hollingworth_chandler_2002_b.cellml |
| | | baylor_hollingworth_chandler_2002/rawfile/HEAD/baylor_hollingworth_chandler_2002_d.cellml |
| | | luo_rudy_1994/rawfile/HEAD/luo_rudy_1994.cellml |
| | | colegrove_albrecht_friel_2000/rawfile/HEAD/colegrove_albrecht_friel_2000.cellml |
| | | dougherty_wright_yew_2005/rawfile/HEAD/dougherty_wright_yew_2005.cellml |
| | | yamaguchi_takaki_matsubara_yasuhara_suga_1996/rawfile/HEAD/yamaguchi_takaki_matsubara_yasuhara_suga_1996.cellml |

| | | |
|---|---|---|
| | | boyett_zhang_garny_holden_2001/rawfile/HEAD/boyett_zhang_garny_holden_2001.cellml |
| | | iribe_kohl_noble_2006/rawfile/HEAD/iribe_kohl_noble_2006.cellml |
| | | izakov_katsnelson_blyakhman_markhasin_shkylar_1991/rawfile/HEAD/izakov_katsnelson_blyakhman_markhasin_shkylar_1991.cellml |
| | | stern_song_sham_yang_boheler_rios_1999/rawfile/HEAD/stern_song_sham_yang_boheler_rios_1999.cellml |
| | | devries_sherman_2000/rawfile/HEAD/devries_sherman_2000.cellml |
| | | marhl_haberichter_brumen_heinrich_2000/rawfile/HEAD/marhl_haberichter_brumen_heinrich_2000.cellml |
| 84 | interaction of glycolysis and mitochondrial respiration in metabolic oscillations of pancreatic islets | 570/rawfile/HEAD/bertram_satin_pedersen_luciani_sherman_2007.cellml |
| 85 | luo rudy model | baylor_hollingworth_chandler_2002/rawfile/HEAD/baylor_hollingworth_chandler_2002_b.cellml |
| | | baylor_hollingworth_chandler_2002/rawfile/HEAD/baylor_hollingworth_chandler_2002_d.cellml |
| | | luo_rudy_1994/rawfile/HEAD/luo_rudy_1994.cellml |
| | | colegrove_albrecht_friel_2000/rawfile/HEAD/colegrove_albrecht_friel_2000.cellml |
| | | dougherty_wright_yew_2005/rawfile/HEAD/dougherty_wright_yew_2005.cellml |
| | | yamaguchi_takaki_matsubara_yasuhara_suga_1996/rawfile/HEAD/yamaguchi_takaki_matsubara_yasuhara_suga_1996.cellml |
| | | boyett_zhang_garny_holden_2001/rawfile/HEAD/boyett_zhang_garny_holden_2001.cellml |
| | | iribe_kohl_noble_2006/rawfile/HEAD/iribe_kohl_noble_2006.cellml |
| | | izakov_katsnelson_blyakhman_markhasin_shkylar_1991/rawfile/HEAD/izakov_katsnelson_blyakhman_markhasin_shkylar_1991.cellml |
| | | stern_song_sham_yang_boheler_rios_1999/rawfile/HEAD/stern_song_sham_yang_boheler_rios_1999.cellml |
| | | devries_sherman_2000/rawfile/HEAD/devries_sherman_2000.cellml |
| | | marhl_haberichter_brumen_heinrich_2000/rawfile/HEAD/marhl_haberichter_brumen_heinrich_2000.cellml |
| 86 | di francesco-noble purkinje fibre model 1985 | difrancesco_noble_1985/rawfile/HEAD/difrancesco_noble_1985.cellml |
| 87 | computational models of ventricular | iyer_mazhari_winslow_2004/rawfile/HEAD/iyer_mazhari_winslow_2004.cellml |
| 88 | cardiac | baylor_hollingworth_chandler_2002/rawfile/HEAD/baylor_hollingworth_chandler_2002_b.cellml |
| | | baylor_hollingworth_chandler_2002/rawfile/HEAD/baylor_hollingworth_chandler_2002_d.cellml |
| | | luo_rudy_1994/rawfile/HEAD/luo_rudy_1994.cellml |
| | | colegrove_albrecht_friel_2000/rawfile/HEAD/colegrove_albrecht_friel_2000.cellml |
| | | dougherty_wright_yew_2005/rawfile/HEAD/dougherty_wright_yew_2005.cellml |
| | | yamaguchi_takaki_matsubara_yasuhara_suga_1996/rawfile/HEAD/yamaguchi_takaki_matsubara_yasuhara_suga_1996.cellml |
| | | boyett_zhang_garny_holden_2001/rawfile/HEAD/boyett_zhang_garny_holden_2001.cellml |
| | | iribe_kohl_noble_2006/rawfile/HEAD/iribe_kohl_noble_2006.cellml |
| | | izakov_katsnelson_blyakhman_markhasin_shkylar_1991/rawfile/HEAD/izakov_katsnelson_blyakhman_markhasin_shkylar_1991.cellml |
| | | stern_song_sham_yang_boheler_rios_1999/rawfile/HEAD/stern_song_sham_yang_boheler_rios_1999.cellml |
| | | devries_sherman_2000/rawfile/HEAD/devries_sherman_2000.cellml |
| | | marhl_haberichter_brumen_heinrich_2000/rawfile/HEAD/marhl_haberichter_brumen_heinrich_2000.cellml |
| 89 | human ventricular | tentusscher_noble_noble_panfilov_2004/rawfile/HEAD/tentusscher_noble_noble_panfilov_2004_c.cellml |
| | | tentusscher_noble_noble_panfilov_2004/rawfile/HEAD/tentusscher_noble_noble_panfilov_2004_b.cellml |
| | | tentusscher_noble_noble_panfilov_2004/rawfile/HEAD/tentusscher_noble_noble_panfilov_2004_a.cellml |
| 90 | bueno-orovio-cherry-fenton model | difrancesco_noble_1985/rawfile/HEAD/difrancesco_noble_1985.cellml |
| 91 | temperature | baylor_hollingworth_chandler_2002/rawfile/HEAD/baylor_hollingworth_chandler_2002_b.cellml |
| | | baylor_hollingworth_chandler_2002/rawfile/HEAD/baylor_hollingworth_chandler_2002_d.cellml |
| | | luo_rudy_1994/rawfile/HEAD/luo_rudy_1994.cellml |
| | | colegrove_albrecht_friel_2000/rawfile/HEAD/colegrove_albrecht_friel_2000.cellml |
| | | dougherty_wright_yew_2005/rawfile/HEAD/dougherty_wright_yew_2005.cellml |
| | | yamaguchi_takaki_matsubara_yasuhara_suga_1996/rawfile/HEAD/yamaguchi_takaki_matsubara_yasuhara_suga_1996.cellml |
| | | boyett_zhang_garny_holden_2001/rawfile/HEAD/boyett_zhang_garny_holden_2001.cellml |
| | | iribe_kohl_noble_2006/rawfile/HEAD/iribe_kohl_noble_2006.cellml |
| | | izakov_katsnelson_blyakhman_markhasin_shkylar_1991/rawfile/HEAD/izakov_katsnelson_blyakhman_markhasin_shkylar_1991.cellml |
| | | stern_song_sham_yang_boheler_rios_1999/rawfile/HEAD/stern_song_sham_yang_boheler_rios_1999.cellml |
| | | devries_sherman_2000/rawfile/HEAD/devries_sherman_2000.cellml |
| | | marhl_haberichter_brumen_heinrich_2000/rawfile/HEAD/marhl_haberichter_brumen_heinrich_2000.cellml |
| 92 | adenosine brain | 546/rawfile/HEAD/cloutier_2009.cellml |
| 93 | sglt1 | 267/rawfile/HEAD/SEDML/mackenzie_1996/mackenzie_1996.cellml |
| | | 267/rawfile/HEAD/mackenzie_1996.cellml |
| | | 267/rawfile/HEAD/SEDML/eskandari_2005/eskandari_2005.cellml |
| | | 267/rawfile/HEAD/eskandari_2005.cellml |
| | | 267/rawfile/HEAD/mackenzie_1996-mouse-baso.cellml |

| | | |
|---|---|---|
| 94 | a model of cardiac electrical activity incorporating ionic pumps and concentration changes | difrancesco_noble_1985/rawfile/HEAD/difrancesco_noble_1985.cellml |
| 95 | oxidative phosphorylation | baylor_hollingworth_chandler_2002/rawfile/HEAD/baylor_hollingworth_chandler_2002_b.cellml |
| | | baylor_hollingworth_chandler_2002/rawfile/HEAD/baylor_hollingworth_chandler_2002_d.cellml |
| | | luo_rudy_1994/rawfile/HEAD/luo_rudy_1994.cellml |
| | | colegrove_albrecht_friel_2000/rawfile/HEAD/colegrove_albrecht_friel_2000.cellml |
| | | dougherty_wright_yew_2005/rawfile/HEAD/dougherty_wright_yew_2005.cellml |
| | | yamaguchi_takaki_matsubara_yasuhara_suga_1996/rawfile/HEAD/yamaguchi_takaki_matsubara_yasuhara_suga_1996.cellml |
| | | boyett_zhang_garny_holden_2001/rawfile/HEAD/boyett_zhang_garny_holden_2001.cellml |
| | | iribe_kohl_noble_2006/rawfile/HEAD/iribe_kohl_noble_2006.cellml |
| | | 546/rawfile/HEAD/cloutier_2009.cellml |
| | | izakov_katsnelson_blyakhman_markhasin_shkylar_1991/rawfile/HEAD/izakov_katsnelson_blyakhman_markhasin_shkylar_1991.cellml |
| | | stern_song_sham_yang_boheler_rios_1999/rawfile/HEAD/stern_song_sham_yang_boheler_rios_1999.cellml |
| | | devries_sherman_2000/rawfile/HEAD/devries_sherman_2000.cellml |
| | | marhl_haberichter_brumen_heinrich_2000/rawfile/HEAD/marhl_haberichter_brumen_heinrich_2000.cellml |
| 96 | rat | baylor_hollingworth_chandler_2002/rawfile/HEAD/baylor_hollingworth_chandler_2002_b.cellml |
| | | baylor_hollingworth_chandler_2002/rawfile/HEAD/baylor_hollingworth_chandler_2002_d.cellml |
| | | luo_rudy_1994/rawfile/HEAD/luo_rudy_1994.cellml |
| | | colegrove_albrecht_friel_2000/rawfile/HEAD/colegrove_albrecht_friel_2000.cellml |
| | | dougherty_wright_yew_2005/rawfile/HEAD/dougherty_wright_yew_2005.cellml |
| | | yamaguchi_takaki_matsubara_yasuhara_suga_1996/rawfile/HEAD/yamaguchi_takaki_matsubara_yasuhara_suga_1996.cellml |
| | | boyett_zhang_garny_holden_2001/rawfile/HEAD/boyett_zhang_garny_holden_2001.cellml |
| | | iribe_kohl_noble_2006/rawfile/HEAD/iribe_kohl_noble_2006.cellml |
| | | izakov_katsnelson_blyakhman_markhasin_shkylar_1991/rawfile/HEAD/izakov_katsnelson_blyakhman_markhasin_shkylar_1991.cellml |
| | | stern_song_sham_yang_boheler_rios_1999/rawfile/HEAD/stern_song_sham_yang_boheler_rios_1999.cellml |
| | | devries_sherman_2000/rawfile/HEAD/devries_sherman_2000.cellml |
| | | marhl_haberichter_brumen_heinrich_2000/rawfile/HEAD/marhl_haberichter_brumen_heinrich_2000.cellml |
| 97 | glucose | baylor_hollingworth_chandler_2002/rawfile/HEAD/baylor_hollingworth_chandler_2002_b.cellml |
| | | baylor_hollingworth_chandler_2002/rawfile/HEAD/baylor_hollingworth_chandler_2002_d.cellml |
| | | 267/rawfile/HEAD/eskandari_2005.cellml |
| | | luo_rudy_1994/rawfile/HEAD/luo_rudy_1994.cellml |
| | | colegrove_albrecht_friel_2000/rawfile/HEAD/colegrove_albrecht_friel_2000.cellml |
| | | dougherty_wright_yew_2005/rawfile/HEAD/dougherty_wright_yew_2005.cellml |
| | | yamaguchi_takaki_matsubara_yasuhara_suga_1996/rawfile/HEAD/yamaguchi_takaki_matsubara_yasuhara_suga_1996.cellml |
| | | boyett_zhang_garny_holden_2001/rawfile/HEAD/boyett_zhang_garny_holden_2001.cellml |
| | | 267/rawfile/HEAD/SEDML/eskandari_2005/eskandari_2005.cellml |
| | | 570/rawfile/HEAD/bertram_satin_pedersen_luciani_sherman_2007.cellml |
| | | iribe_kohl_noble_2006/rawfile/HEAD/iribe_kohl_noble_2006.cellml |
| | | izakov_katsnelson_blyakhman_markhasin_shkylar_1991/rawfile/HEAD/izakov_katsnelson_blyakhman_markhasin_shkylar_1991.cellml |
| | | stern_song_sham_yang_boheler_rios_1999/rawfile/HEAD/stern_song_sham_yang_boheler_rios_1999.cellml |
| | | devries_sherman_2000/rawfile/HEAD/devries_sherman_2000.cellml |
| | | marhl_haberichter_brumen_heinrich_2000/rawfile/HEAD/marhl_haberichter_brumen_heinrich_2000.cellml |
| 98 | pancreatic alpha cells | baylor_hollingworth_chandler_2002/rawfile/HEAD/baylor_hollingworth_chandler_2002_b.cellml |
| | | baylor_hollingworth_chandler_2002/rawfile/HEAD/baylor_hollingworth_chandler_2002_d.cellml |
| | | luo_rudy_1994/rawfile/HEAD/luo_rudy_1994.cellml |
| | | colegrove_albrecht_friel_2000/rawfile/HEAD/colegrove_albrecht_friel_2000.cellml |
| | | dougherty_wright_yew_2005/rawfile/HEAD/dougherty_wright_yew_2005.cellml |
| | | yamaguchi_takaki_matsubara_yasuhara_suga_1996/rawfile/HEAD/yamaguchi_takaki_matsubara_yasuhara_suga_1996.cellml |
| | | boyett_zhang_garny_holden_2001/rawfile/HEAD/boyett_zhang_garny_holden_2001.cellml |
| | | iribe_kohl_noble_2006/rawfile/HEAD/iribe_kohl_noble_2006.cellml |
| | | izakov_katsnelson_blyakhman_markhasin_shkylar_1991/rawfile/HEAD/izakov_katsnelson_blyakhman_markhasin_shkylar_1991.cellml |
| | | stern_song_sham_yang_boheler_rios_1999/rawfile/HEAD/stern_song_sham_yang_boheler_rios_1999.cellml |
| | | devries_sherman_2000/rawfile/HEAD/devries_sherman_2000.cellml |
| | | marhl_haberichter_brumen_heinrich_2000/rawfile/HEAD/marhl_haberichter_brumen_heinrich_2000.cellml |
| | | iyer_mazhari_winslow_2004/rawfile/HEAD/iyer_mazhari_winslow_2004.cellml |
| | | baylor_hollingworth_chandler_2002/rawfile/HEAD/baylor_hollingworth_chandler_2002_b.cellml |
| | | tentusscher_noble_noble_panfilov_2004/rawfile/HEAD/tentusscher_noble_noble_panfilov_2004_b.cellml |
| | | baylor_hollingworth_chandler_2002/rawfile/HEAD/baylor_hollingworth_chandler_2002_d.cellml |

| | | |
|---|---|---|
| | | faber_rudy_2000/rawfile/HEAD/faber_rudy_2000.cellml |
| | | luo_rudy_1994/rawfile/HEAD/luo_rudy_1994.cellml |
| | | colegrove_albrecht_friel_2000/rawfile/HEAD/colegrove_albrecht_friel_2000.cellml |
| | | 55c/rawfile/HEAD/Hinch_et_al_2004.cellml |
| | | dougherty_wright_yew_2005/rawfile/HEAD/dougherty_wright_yew_2005.cellml |
| | | 556/rawfile/HEAD/niederer_hunter_smith_2006.cellml |
| | | yamaguchi_takaki_matsubara_yasuhara_suga_1996/rawfile/HEAD/yamaguchi_takaki_matsubara_yasuhara_suga_1996.cellml |
| | | boyett_zhang_garny_holden_2001/rawfile/HEAD/boyett_zhang_garny_holden_2001.cellml |
| | | tentusscher_noble_noble_panfilov_2004/rawfile/HEAD/tentusscher_noble_noble_panfilov_2004_a.cellml |
| | | tentusscher_noble_noble_panfilov_2004/rawfile/HEAD/tentusscher_noble_noble_panfilov_2004_c.cellml |
| | | terkildsen_niederer_crampin_hunter_smith_2008/rawfile/HEAD/Hinch_et_al_2004.cellml |
| | | hinch_greenstein_tanskanen_xu_winslow_2004/rawfile/HEAD/hinch_greenstein_tanskanen_xu_winslow_2004.cellml |
| | | iribe_kohl_noble_2006/rawfile/HEAD/iribe_kohl_noble_2006.cellml |
| | | shannon_wang_puglisi_weber_bers_2004/rawfile/HEAD/shannon_wang_puglisi_weber_bers_2004_a.cellml |
| | | niederer_hunter_smith_2006/rawfile/HEAD/niederer_hunter_smith_2006.cellml |
| | | izakov_katsnelson_blyakhman_markhasin_shkylar_1991/rawfile/HEAD/izakov_katsnelson_blyakhman_markhasin_shkylar_1991.cellml |
| | | 563/rawfile/HEAD/saucerman_brunton_michailova_mcculloch_2003.cellml |
| | | stern_song_sham_yang_boheler_rios_1999/rawfile/HEAD/stern_song_sham_yang_boheler_rios_1999.cellml |
| | | shannon_wang_puglisi_weber_bers_2004/rawfile/HEAD/shannon_wang_puglisi_weber_bers_2004_b.cellml |
| | | devries_sherman_2000/rawfile/HEAD/devries_sherman_2000.cellml |
| | | marhl_haberichter_brumen_heinrich_2000/rawfile/HEAD/marhl_haberichter_brumen_heinrich_2000.cellml |
| 100 | network | baylor_hollingworth_chandler_2002/rawfile/HEAD/baylor_hollingworth_chandler_2002_b.cellml |
| | | baylor_hollingworth_chandler_2002/rawfile/HEAD/baylor_hollingworth_chandler_2002_d.cellml |
| | | luo_rudy_1994/rawfile/HEAD/luo_rudy_1994.cellml |
| | | colegrove_albrecht_friel_2000/rawfile/HEAD/colegrove_albrecht_friel_2000.cellml |
| | | dougherty_wright_yew_2005/rawfile/HEAD/dougherty_wright_yew_2005.cellml |
| | | yamaguchi_takaki_matsubara_yasuhara_suga_1996/rawfile/HEAD/yamaguchi_takaki_matsubara_yasuhara_suga_1996.cellml |
| | | boyett_zhang_garny_holden_2001/rawfile/HEAD/boyett_zhang_garny_holden_2001.cellml |
| | | iribe_kohl_noble_2006/rawfile/HEAD/iribe_kohl_noble_2006.cellml |
| | | izakov_katsnelson_blyakhman_markhasin_shkylar_1991/rawfile/HEAD/izakov_katsnelson_blyakhman_markhasin_shkylar_1991.cellml |
| | | stern_song_sham_yang_boheler_rios_1999/rawfile/HEAD/stern_song_sham_yang_boheler_rios_1999.cellml |
| | | devries_sherman_2000/rawfile/HEAD/devries_sherman_2000.cellml |
| | | marhl_haberichter_brumen_heinrich_2000/rawfile/HEAD/marhl_haberichter_brumen_heinrich_2000.cellml |
| 101 | graph | baylor_hollingworth_chandler_2002/rawfile/HEAD/baylor_hollingworth_chandler_2002_b.cellml |
| | | baylor_hollingworth_chandler_2002/rawfile/HEAD/baylor_hollingworth_chandler_2002_d.cellml |
| | | luo_rudy_1994/rawfile/HEAD/luo_rudy_1994.cellml |
| | | colegrove_albrecht_friel_2000/rawfile/HEAD/colegrove_albrecht_friel_2000.cellml |
| | | dougherty_wright_yew_2005/rawfile/HEAD/dougherty_wright_yew_2005.cellml |
| | | yamaguchi_takaki_matsubara_yasuhara_suga_1996/rawfile/HEAD/yamaguchi_takaki_matsubara_yasuhara_suga_1996.cellml |
| | | boyett_zhang_garny_holden_2001/rawfile/HEAD/boyett_zhang_garny_holden_2001.cellml |
| | | iribe_kohl_noble_2006/rawfile/HEAD/iribe_kohl_noble_2006.cellml |
| | | izakov_katsnelson_blyakhman_markhasin_shkylar_1991/rawfile/HEAD/izakov_katsnelson_blyakhman_markhasin_shkylar_1991.cellml |
| | | stern_song_sham_yang_boheler_rios_1999/rawfile/HEAD/stern_song_sham_yang_boheler_rios_1999.cellml |
| | | devries_sherman_2000/rawfile/HEAD/devries_sherman_2000.cellml |
| | | marhl_haberichter_brumen_heinrich_2000/rawfile/HEAD/marhl_haberichter_brumen_heinrich_2000.cellml |
| 102 | mouse ventricular myocyte action potential calcium transients cpvt | 563/rawfile/HEAD/saucerman_brunton_michailova_mcculloch_2003.cellml |
| 103 | growth hormone | baylor_hollingworth_chandler_2002/rawfile/HEAD/baylor_hollingworth_chandler_2002_b.cellml |
| | | baylor_hollingworth_chandler_2002/rawfile/HEAD/baylor_hollingworth_chandler_2002_d.cellml |
| | | luo_rudy_1994/rawfile/HEAD/luo_rudy_1994.cellml |
| | | colegrove_albrecht_friel_2000/rawfile/HEAD/colegrove_albrecht_friel_2000.cellml |
| | | dougherty_wright_yew_2005/rawfile/HEAD/dougherty_wright_yew_2005.cellml |
| | | yamaguchi_takaki_matsubara_yasuhara_suga_1996/rawfile/HEAD/yamaguchi_takaki_matsubara_yasuhara_suga_1996.cellml |
| | | boyett_zhang_garny_holden_2001/rawfile/HEAD/boyett_zhang_garny_holden_2001.cellml |
| | | iribe_kohl_noble_2006/rawfile/HEAD/iribe_kohl_noble_2006.cellml |
| | | izakov_katsnelson_blyakhman_markhasin_shkylar_1991/rawfile/HEAD/izakov_katsnelson_blyakhman_markhasin_shkylar_1991.cellml |
| | | stern_song_sham_yang_boheler_rios_1999/rawfile/HEAD/stern_song_sham_yang_boheler_rios_1999.cellml |
| | | devries_sherman_2000/rawfile/HEAD/devries_sherman_2000.cellml |
| | | marhl_haberichter_brumen_heinrich_2000/rawfile/HEAD/marhl_haberichter_brumen_heinrich_2000.cellml |
| | | baylor_hollingworth_chandler_2002/rawfile/HEAD/baylor_hollingworth_chandler_2002_b.cellml |



104 shorten wall

| | | |
|---|---|---|
| | | baylor_hollingworth_chandler_2002/rawfile/HEAD/baylor_hollingworth_chandler_2002_d.cellml |
| | | luo_rudy_1994/rawfile/HEAD/luo_rudy_1994.cellml |
| | | colegrove_albrecht_friel_2000/rawfile/HEAD/colegrove_albrecht_friel_2000.cellml |
| | | dougherty_wright_yew_2005/rawfile/HEAD/dougherty_wright_yew_2005.cellml |
| | | yamaguchi_takaki_matsubara_yasuhara_suga_1996/rawfile/HEAD/yamaguchi_takaki_matsubara_yasuhara_suga_1996.cellml |
| | | boyett_zhang_garny_holden_2001/rawfile/HEAD/boyett_zhang_garny_holden_2001.cellml |
| | | iribe_kohl_noble_2006/rawfile/HEAD/iribe_kohl_noble_2006.cellml |
| | | shorten_robson_mckinnon_wall_2000/rawfile/HEAD/shorten_robson_mckinnon_wall_2000.cellml |
| | | izakov_katsnelson_blyakhman_markhasin_shkylar_1991/rawfile/HEAD/izakov_katsnelson_blyakhman_markhasin_shkylar_1991.cellml |
| | | stern_song_sham_yang_boheler_rios_1999/rawfile/HEAD/stern_song_sham_yang_boheler_rios_1999.cellml |
| | | devries_sherman_2000/rawfile/HEAD/devries_sherman_2000.cellml |
| | | marhl_haberichter_brumen_heinrich_2000/rawfile/HEAD/marhl_haberichter_brumen_heinrich_2000.cellml |
| 105 | zhang h, holden av, kodama i, honjo h, lei m, varghese t, boyett mr | baylor_hollingworth_chandler_2002/rawfile/HEAD/baylor_hollingworth_chandler_2002_b.cellml |
| | | baylor_hollingworth_chandler_2002/rawfile/HEAD/baylor_hollingworth_chandler_2002_d.cellml |
| | | luo_rudy_1994/rawfile/HEAD/luo_rudy_1994.cellml |
| | | colegrove_albrecht_friel_2000/rawfile/HEAD/colegrove_albrecht_friel_2000.cellml |
| | | dougherty_wright_yew_2005/rawfile/HEAD/dougherty_wright_yew_2005.cellml |
| | | yamaguchi_takaki_matsubara_yasuhara_suga_1996/rawfile/HEAD/yamaguchi_takaki_matsubara_yasuhara_suga_1996.cellml |
| | | boyett_zhang_garny_holden_2001/rawfile/HEAD/boyett_zhang_garny_holden_2001.cellml |
| | | iribe_kohl_noble_2006/rawfile/HEAD/iribe_kohl_noble_2006.cellml |
| | | izakov_katsnelson_blyakhman_markhasin_shkylar_1991/rawfile/HEAD/izakov_katsnelson_blyakhman_markhasin_shkylar_1991.cellml |
| | | stern_song_sham_yang_boheler_rios_1999/rawfile/HEAD/stern_song_sham_yang_boheler_rios_1999.cellml |
| | | devries_sherman_2000/rawfile/HEAD/devries_sherman_2000.cellml |
| | | marhl_haberichter_brumen_heinrich_2000/rawfile/HEAD/marhl_haberichter_brumen_heinrich_2000.cellml |
| 106 | a synthetic oscillatory network of transciptional regulators | baylor_hollingworth_chandler_2002/rawfile/HEAD/baylor_hollingworth_chandler_2002_b.cellml |
| | | baylor_hollingworth_chandler_2002/rawfile/HEAD/baylor_hollingworth_chandler_2002_d.cellml |
| | | luo_rudy_1994/rawfile/HEAD/luo_rudy_1994.cellml |
| | | colegrove_albrecht_friel_2000/rawfile/HEAD/colegrove_albrecht_friel_2000.cellml |
| | | dougherty_wright_yew_2005/rawfile/HEAD/dougherty_wright_yew_2005.cellml |
| | | yamaguchi_takaki_matsubara_yasuhara_suga_1996/rawfile/HEAD/yamaguchi_takaki_matsubara_yasuhara_suga_1996.cellml |
| | | boyett_zhang_garny_holden_2001/rawfile/HEAD/boyett_zhang_garny_holden_2001.cellml |
| | | iribe_kohl_noble_2006/rawfile/HEAD/iribe_kohl_noble_2006.cellml |
| | | izakov_katsnelson_blyakhman_markhasin_shkylar_1991/rawfile/HEAD/izakov_katsnelson_blyakhman_markhasin_shkylar_1991.cellml |
| | | stern_song_sham_yang_boheler_rios_1999/rawfile/HEAD/stern_song_sham_yang_boheler_rios_1999.cellml |
| | | devries_sherman_2000/rawfile/HEAD/devries_sherman_2000.cellml |
| | | marhl_haberichter_brumen_heinrich_2000/rawfile/HEAD/marhl_haberichter_brumen_heinrich_2000.cellml |
| 107 | a model of the ventricular cardiac action | luo_rudy_1991/rawfile/HEAD/luo_rudy_1991.cellml |
| 108 | the ord human ventricular action potential model | w/andre/SAN-ORd/rawfile/HEAD/Ohara_Rudy_2011.cellml |
| 109 | computational model for emergent dynamics in the heart | baylor_hollingworth_chandler_2002/rawfile/HEAD/baylor_hollingworth_chandler_2002_b.cellml |
| | | baylor_hollingworth_chandler_2002/rawfile/HEAD/baylor_hollingworth_chandler_2002_d.cellml |
| | | luo_rudy_1994/rawfile/HEAD/luo_rudy_1994.cellml |
| | | colegrove_albrecht_friel_2000/rawfile/HEAD/colegrove_albrecht_friel_2000.cellml |
| | | dougherty_wright_yew_2005/rawfile/HEAD/dougherty_wright_yew_2005.cellml |
| | | yamaguchi_takaki_matsubara_yasuhara_suga_1996/rawfile/HEAD/yamaguchi_takaki_matsubara_yasuhara_suga_1996.cellml |
| | | boyett_zhang_garny_holden_2001/rawfile/HEAD/boyett_zhang_garny_holden_2001.cellml |
| | | iribe_kohl_noble_2006/rawfile/HEAD/iribe_kohl_noble_2006.cellml |
| | | difrancesco_noble_1985/rawfile/HEAD/difrancesco_noble_1985.cellml |
| | | izakov_katsnelson_blyakhman_markhasin_shkylar_1991/rawfile/HEAD/izakov_katsnelson_blyakhman_markhasin_shkylar_1991.cellml |
| | | stern_song_sham_yang_boheler_rios_1999/rawfile/HEAD/stern_song_sham_yang_boheler_rios_1999.cellml |
| | | devries_sherman_2000/rawfile/HEAD/devries_sherman_2000.cellml |
| | | marhl_haberichter_brumen_heinrich_2000/rawfile/HEAD/marhl_haberichter_brumen_heinrich_2000.cellml |
| 110 | a model of excitation and adaptation in bacterial chemotaxis | baylor_hollingworth_chandler_2002/rawfile/HEAD/baylor_hollingworth_chandler_2002_b.cellml |
| | | baylor_hollingworth_chandler_2002/rawfile/HEAD/baylor_hollingworth_chandler_2002_d.cellml |
| | | luo_rudy_1994/rawfile/HEAD/luo_rudy_1994.cellml |
| | | colegrove_albrecht_friel_2000/rawfile/HEAD/colegrove_albrecht_friel_2000.cellml |
| | | dougherty_wright_yew_2005/rawfile/HEAD/dougherty_wright_yew_2005.cellml |
| | | yamaguchi_takaki_matsubara_yasuhara_suga_1996/rawfile/HEAD/yamaguchi_takaki_matsubara_yasuhara_suga_1996.cellml |
| | | boyett_zhang_garny_holden_2001/rawfile/HEAD/boyett_zhang_garny_holden_2001.cellml |
| | | iribe_kohl_noble_2006/rawfile/HEAD/iribe_kohl_noble_2006.cellml |
| | | izakov_katsnelson_blyakhman_markhasin_shkylar_1991/rawfile/HEAD/izakov_katsnelson_blyakhman_markhasin_shkylar_1991.cellml |

| | | |
|---|---|---|
| | | stern_song_sham_yang_boheler_rios_1999/rawfile/HEAD/stern_song_sham_yang_boheler_rios_1999.cellml |
| | | devries_sherman_2000/rawfile/HEAD/devries_sherman_2000.cellml |
| | | marhl_haberichter_brumen_heinrich_2000/rawfile/HEAD/marhl_haberichter_brumen_heinrich_2000.cellml |

**Table S3.** The best multipliers setup and results. We examine three feature modifications, i.e. without modification (WPURE), adding the preferred label to other features (WPL), and adding the preferred label to empty features only (WPLE). We also examine three additional scenarios to calculate the degree of association, i.e. considering all features with dependency level (mode_1), considering all features without dependency level (mode_2), and considering one feature only from ontology dictionaries with the highest weight combined with description feature from the model entity and dependency level (mode_3).

| Feature modification | Similarity measure | Parser | $(\alpha\ \beta\ \gamma\ \theta\ \delta)$ | $AUC_{PR}$ |
|---|---|---|---|---|
| wple | mode_1 | benepar | (3.0, 0.0, 0.0, 0.0, 0.81) | 0.387 |
| | | stanza | (3.0, 0.4, 0.0, 0.0, 0.13) | 0.452 |
| | | coreNLP | (3.0, 0.0, 0.0, 0.0, 0.16) | 0.528 |
| | | xStanza | (3.0, 0.4, 0.0, 0.0, 0.26) | 0.472 |
| | mode_2 | benepar | (3.0, 0.0, 0.0, 0.0, 1.41) | 0.344 |
| | | stanza | (3.0, 0.7, 0.0, 0.0, 2.67) | 0.333 |
| | | coreNLP | (3.0, 1.0, 0.3, 0.3, 1.67) | 0.405 |
| | | xStanza | (3.0, 0.0, 0.0, 0.0, 0.19) | 0.371 |
| | mode_3 | benepar | (3.0, 0.5, 0.0, 0.1, 0.86) | 0.39 |
| | | stanza | (3.0, 0.5, 0.0, 0.0, 0.21) | 0.454 |
| | | coreNLP | (3.0, 0.5, 0.0, 0.0, 0.31) | 0.517 |
| | | xStanza | (3.0, 0.5, 0.0, 0.0, 0.21) | 0.474 |
| wpl | mode_1 | benepar | (3.0, 1.1, 0.0, 0.4, 1.12) | 0.395 |
| | | stanza | (3.0, 0.4, 0.0, 0.2, 0.12) | **0.463** |
| | | coreNLP | (3.0, 0.4, 0.0, 0.2, 0.21) | 0.546 |
| | | xStanza | (3.0, 0.3, 0.1, 0.2, 0.12) | 0.476 |
| | mode_2 | benepar | (3.0, 0.9, 0.0, 0.0, 1.8) | 0.362 |
| | | stanza | (3.0, 0.7, 0.0, 0.0, 2.67) | 0.332 |
| | | coreNLP | (3.0, 1.0, 0.0, 0.3, 2.0) | 0.421 |
| | | xStanza | (3.0, 0.1, 0.7, 0.1, 0.29) | 0.376 |
| | mode_3 | benepar | (3.0, 3.0, 0.0, 0.0, 1.5) | **0.408** |
| | | stanza | (3.0, 0.5, 0.0, 0.1, 0.11) | **0.463** |
| | | coreNLP | (3.0, 3.0, 0.0, 0.0, 0.38) | **0.553** |
| | | xStanza | (3.0, 1.0, 0.1, 0.2, 0.12) | **0.478** |
| wpure | mode_1 | benepar | (3.0, 0.0, 0.0, 0.0, 0.81) | 0.387 |
| | | stanza | (3.0, 0.0, 0.0, 0.0, 0.16) | 0.441 |
| | | coreNLP | (3.0, 0.0, 0.0, 0.0, 0.16) | 0.528 |
| | | xStanza | (3.0, 0.0, 0.0, 0.0, 0.16) | 0.457 |
| | mode_2 | benepar | (3.0, 0.0, 0.0, 0.0, 1.41) | 0.344 |
| | | stanza | (3.0, 0.6, 0.0, 0.3, 2.4) | 0.33 |
| | | coreNLP | (3.0, 0.0, 0.2, 0.0, 1.5) | 0.402 |
| | | xStanza | (3.0, 0.0, 0.0, 0.0, 0.19) | 0.371 |
| | mode_3 | benepar | (3.0, 0.9, 0.0, 0.0, 0.94) | 0.372 |
| | | stanza | (3.0, 0.5, 0.0, 0.0, 0.21) | 0.409 |
| | | coreNLP | (3.0, 0.5, 0.0, 0.0, 0.43) | 0.477 |
| | | xStanza | (3.0, 0.6, 0.2, 0.0, 0.6) | 0.436 |

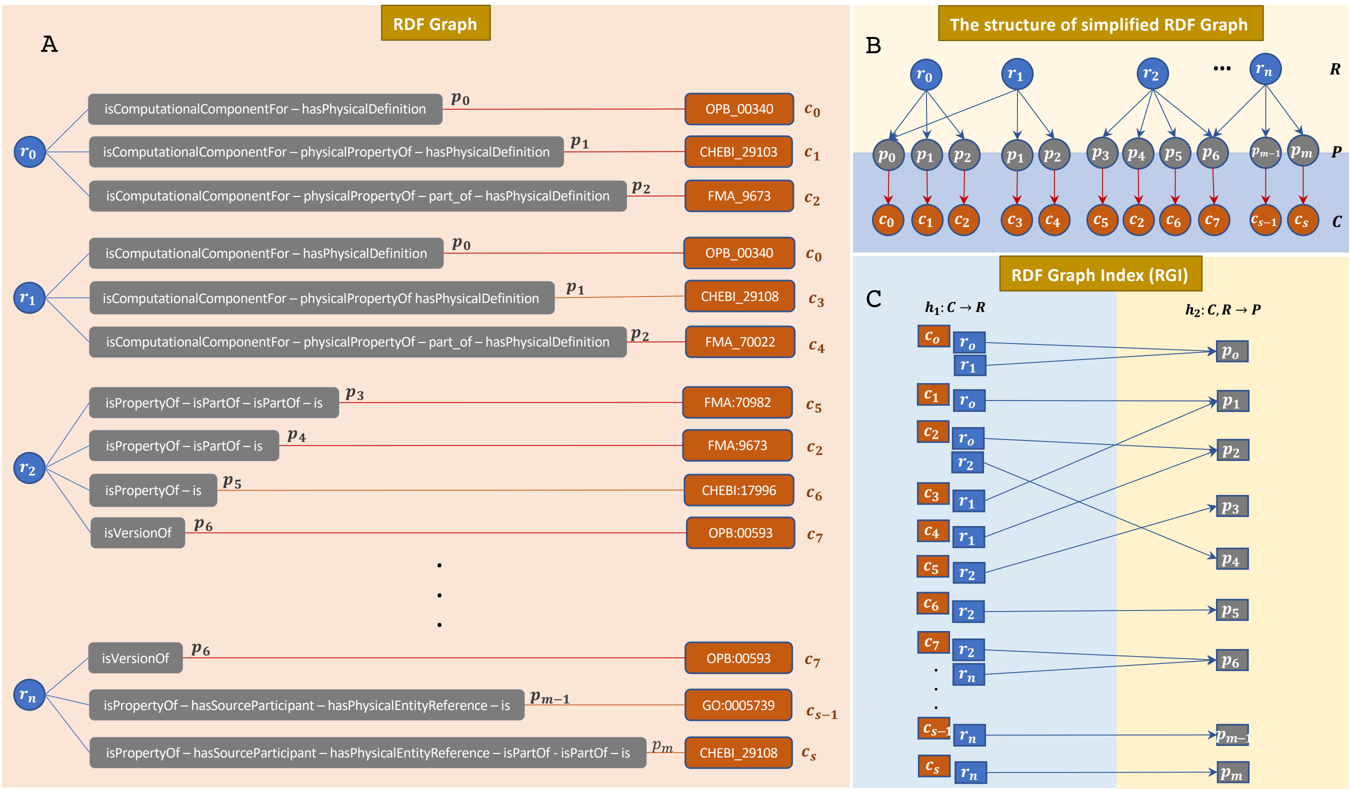


**Figure S1:** The creation of the RDF Graph Index (RGI). (A) The structure of RDF annotation in repositories. There are multiple trees with entities as roots $R$, ontology classes $C$ describing entities $R$, and paths $P$ defining how ontology classes $C$ describing entities $R$. (B) Since a pair of path and ontology class can appear in multiple trees, the structure of the RDF Graph can be simplified by providing a single entry for each pair referenced by multiple roots $\bar{R}$. (C) RDF Graph Index (RGI) consisting of indexes mapping ontology classes $C$ to entities $R$ and $(C, R)$ to paths $P$.

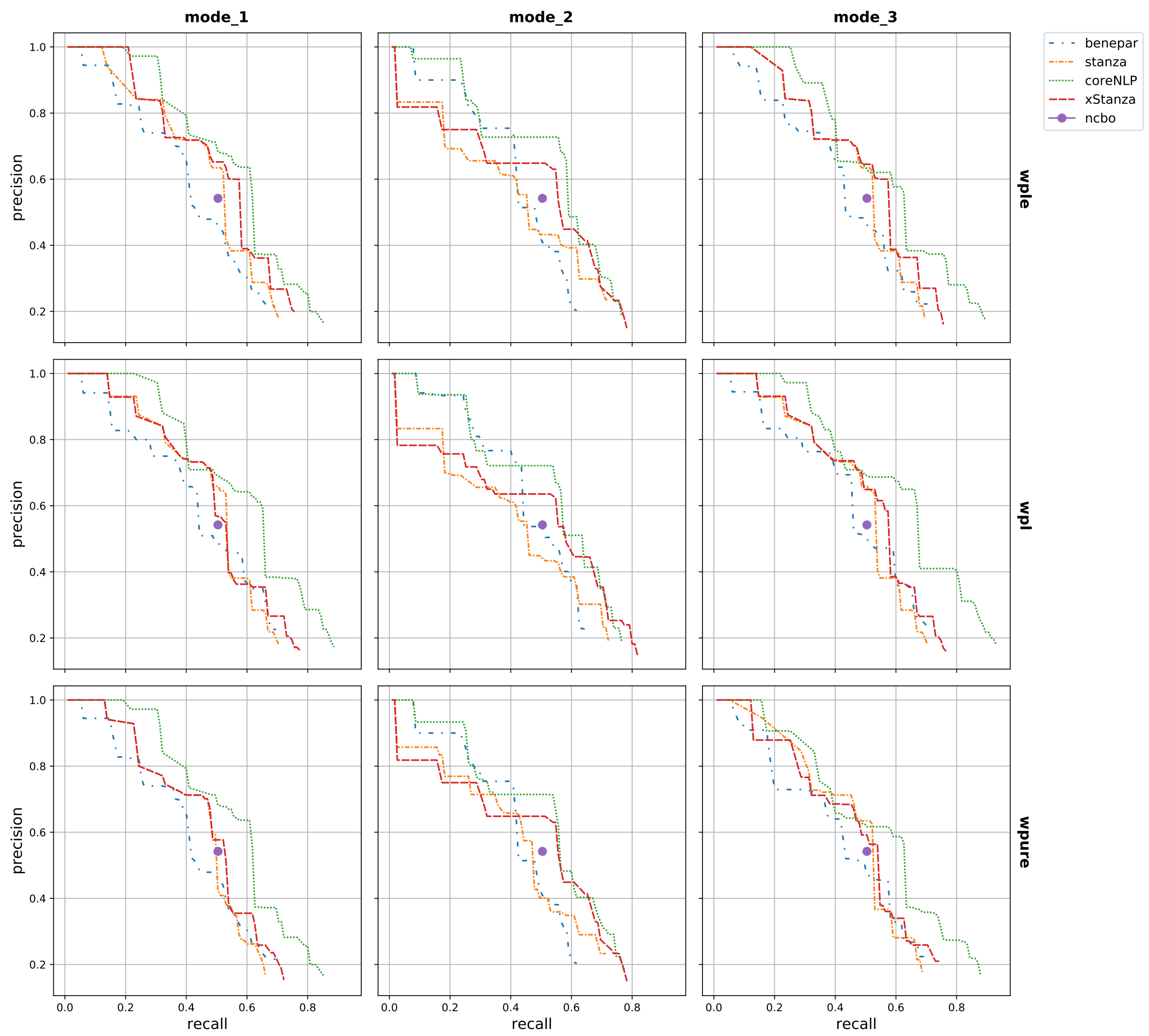


**Figure S2:** The performance of NLIMED to annotate NLQ on a test data containing 52 NLQ. The use of CoreNLP demonstrates the best $AuC_{PR}$ (Precision-Recall AuC, a single value representing the combination of different precision and recall pairs), followed by xStanza. Compared to NCBO Annotator where its precision and recall is 0.542 and 0.504, respectively, almost all parsers perform better. Interpolated $AuC_{PR}$ of NLIMED differentiate with four parsers (CoreNLP, Benepar, Stanza, xStanza), three feature modifications (WPURE, WPL, WPLE), and three scenarios to calculate the degree of association (mode_1, mode_2, mode_3).

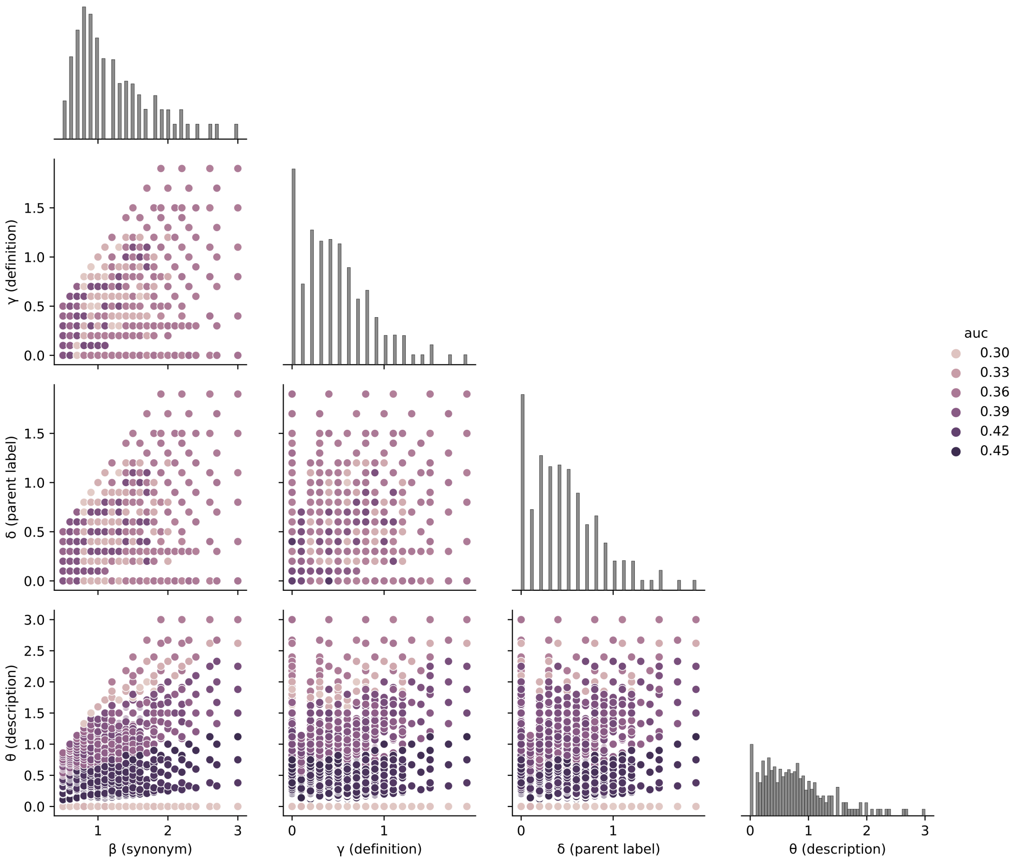


**Figure S3:** The interaction of features and their role in annotating a query to ontology classes using Stanza, WPL, mode_3, and $\alpha$=3.0. The most important feature is the preferred label followed by the description and synonym, whereas the parent label and definition have the least contributions.

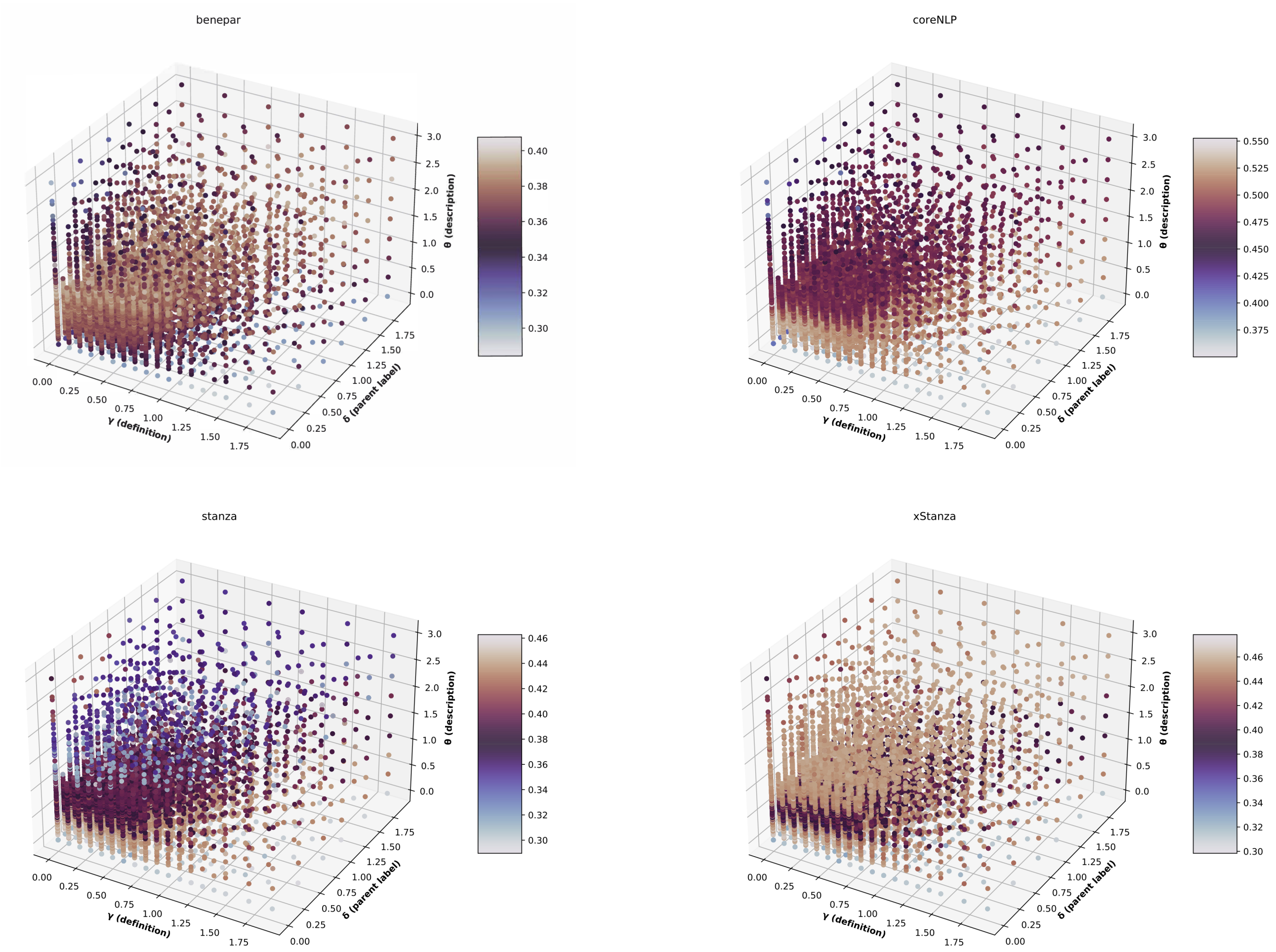


**Figure S4:** The interaction of $\gamma$(definition), $\delta$(parent label), and $\theta$(description) using wpl and mode_3 along with their $AuC_{PR}$

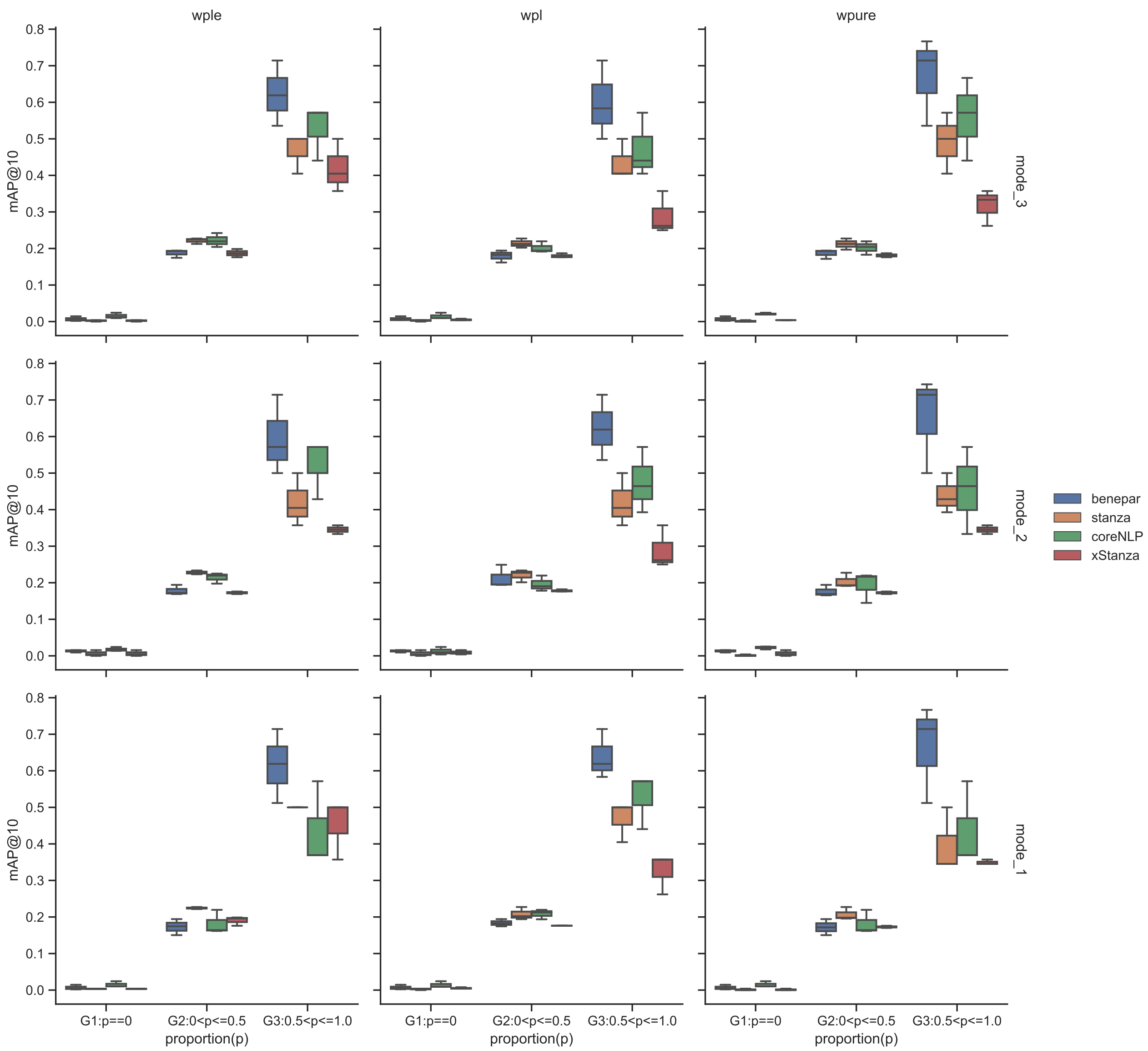


**Figure S5:** The mAP@10 of NLIMED on historical query-results records in the PMR differentiate with the highest proportion of terms in one of their ontology classes appear in the query. The multiplier combinations used are the best combination stated at Subsection **??** with additional combinations of $\alpha$, $\beta$, $\gamma$, $\delta$, and $\theta$ as (3.0, 3.0, 0.5, 0.5, 0.5) and (3.0, 3.0, 1.0, 1.0, 1.0).
